# Cellular stemness identifies high-risk ductal carcinoma *in situ* and offers a therapeutic interception opportunity

**DOI:** 10.64898/2026.05.13.724882

**Authors:** E. Schueddig, V. Kochat, E. Arslan, Y. Dallas, P. Yang, W. Padron, Z. Li, R. Henry, J. Lin, M. Mattohti, R. Madan, T. Fields, S. Golem, S. A. Khan, J. L. Wagner, K. E. Larson, C. Balanoff, A. Aripoli, A. Huppe, O. Winblad, J. Peterson, M. Hill, C. Smith, E. J. Jeffers, L.J. Kilgore, Chenxuan Zang, Peng Wei, N. Navin, C. Fabian, M. T. Lewis, Q. Zhu, A. Thompson, A. K. Godwin, D. C. Koestler, K. Rai, F. Behbod

**Affiliations:** Department of Biostatistics & Data Science, University of Kansas Medical Center, 3901 Rainbow Boulevard, Kansas City, KS 66160, USA; Department of Genomic Medicine and MDACC Epigenomics Therapy Initiative, The University of Texas MD Anderson Cancer Center,1515 Holcombe Boulevard, Houston, TX 77030, USA; Oncology Research and Development, Takeda Pharmaceuticals, 650 East Kendall Street, Cambridge, MA 02142, USA; Department of Pathology and Laboratory Medicine, University of Kansas Medical Center, 3901 Rainbow Boulevard, Kansas City, KS 66160, USA; Department of Biostatistics, The University of Texas MD Anderson Cancer Center,1515 Holcombe Boulevard, Houston, TX 77030, USA; Department of Systems Biology, The University of Texas MD Anderson Cancer Center,1515 Holcombe Boulevard, Houston, TX 77030, USA; Department of Pathology, Feinberg School of Medicine, Northwestern University, 303 East Superior Street, Chicago, IL 60611, USA; Department of Surgery, Immunobiology and Transplant Science Center, Houston Methodist Research Institute, Houston Methodist Hospital, Houston, TX 77030, USA; Department of Surgery, Weill Cornell Medicine, Cornell Univeristy New York, NY 14853, USA; Department of Surgery, University of Kansas Medical Center, 3901 Rainbow Boulevard, Kansas City, KS 66160, USA; Department of Radiology, University of Kansas Medical Center, 3901 Rainbow Boulevard, Kansas City, KS 66160, USA; Department of Medical Oncology, University of Kansas Medical Center, 3901 Rainbow Boulevard, Kansas City, KS 66160, USA; Department of Molecular and Cellular Biology, Baylor College of Medicine, One Baylor Plaza, Houston, TX 77030, USA; Lester and Sue Smith Breast Center, Department of Molecular Genetics, Baylor College of Medicine, One Baylor Plaza, Houston, TX 77030, USA; Department of Surgical Oncology, Division of Breast Surgery, Baylor College of Medicine, One Baylor Plaza, Houston, TX 77030, USA; Kansas Institute for Precision Medicine, University of Kansas Medical Center, 3901 Rainbow Boulevard, Kansas City, KS 66160, USA

**Keywords:** Ductal carcinoma in situ, Invasive ductal carcinoma, stemness, epigenomics, breast cancer, biomarker, therapeutic target

## Abstract

Ductal carcinoma in situ (DCIS) exhibits substantial heterogeneity in its risk of progression to invasive breast cancer, yet the cellular and molecular determinants of high-risk lesions remain incompletely defined. Using spatially resolved single-cell transcriptomic and epigenomic profiling of 43 patient-derived DCIS and DCIS/invasive ductal carcinoma (IDC) samples, we delineate cellular programs, spatial organization, and epigenetic regulatory mechanisms associated with invasive potential. We identify an epithelial population with stemness features within luminal hormone-responsive (LumHR) cells that progressively expands from benign tissue to DCIS and IDC, and is strongly associated with invasive progression and recurrence-linked transcriptional programs. Spatial mapping reveals discrete DCIS niches enriched for stem-like LumHR cells, characterized by elevated CEACAM6 expression and enhanced ligand–receptor interactions, including CEACAM6–EGFR signaling between epithelial and stromal compartments, including cancer-associated fibroblasts, macrophages (*APOC1*-positive) and perivascular cells. These niches define a microenvironmental context that supports stemness and invasive potential. Epigenomic analyses implicate FOXA1 as a key regulator of these stem-like transcriptional states. Pharmacologic disruption of FOXA1-regulatory network using LSD1 inhibition suppresses stemness-associated transcriptional programs in vitro and significantly restrains tumor growth in vivo. Collectively, these findings define high-risk DCIS as a stemness-driven disease embedded within specialized microenvironments, and identify associated regulatory networks as candidate biomarkers and therapeutic vulnerabilities.

## Introduction

Ductal carcinoma in situ (DCIS) is the most common form of non-invasive breast cancer and is widely regarded as a precursor to invasive ductal carcinoma (IDC). Long-term observational studies of untreated DCIS report progression to invasive disease in approximately 14–53% of cases ^1–6^. Despite this heterogeneous natural history, most patients with DCIS undergo multimodality treatment, including surgery, radiation, and endocrine therapy. While these interventions reduce local recurrence, they do not reduce breast cancer-specific mortality ^7^, raising concern that a substantial proportion of patients are overtreated.

Despite decades of investigation, the clinical management of DCIS remains constrained by several unresolved challenges. First, there are no reliable molecular or histopathologic markers that prospectively identify which lesions will progress to invasive or metastatic disease. Second, although current therapies reduce local recurrence, invasive progression and breast cancer–related death still occur ^7–9^. Biomarker discovery efforts, including transcriptomic profiling of DCIS specimens with known outcomes, have identified gene expression signatures that stratify recurrence risk ^10,11^, but these approaches have not yet yielded mechanistically actionable targets for aggressive DCIS.

Recent advances in single-cell transcriptomics and computational biology now enable inference of cellular stemness states at high resolution. CytoTRACE, a validated algorithm for predicting cellular differentiation states from single-cell RNA sequencing data, has demonstrated robust performance across diverse tissues, lineages, platforms, and species ^12^. Integration with single-cell chromatin accessibility profiling has further established concordance between epigenomic organization and CytoTRACE-inferred stemness, implicating chromatin regulation as a key determinant of stem-like states ^12^.

We investigated whether stem-like epithelial states drive the invasive potential of DCIS using integrated single-cell transcriptomic, epigenomic, and spatial profiling. DCIS lesions with invasive potential were enriched for stem-like epithelial cells compared to non-invasive DCIS, accompanied by a shift toward luminal hormone-responsive (LumHR) cells, which emerged as a key lineage in disease progression. These cells exhibited the highest stemness scores, particularly in lesions associated with invasive progression.

Copy number variation analysis showed that although genomic instability increases in progressed models, it does not directly drive stemness. Instead, stem-like states were associated with transcriptional programs involving chromatin organization, MYC signaling, metabolism, and epithelial–mesenchymal transition and were largely independent of proliferation. Stemness also correlated with genes linked to DCIS recurrence.

Spatial analysis identified niches enriched for high-stemness LumHR cells, with evidence of CEACAM6–EGFR signaling interactions in HER2-positive (HER2^+^) samples, suggesting a microenvironment that supports epithelial plasticity. Integrative analyses implicated FOXA1 as a key regulator of this state. Pharmacologic targeting this axis using the LSD1 inhibitor ORY-1001 reduced FOXA1 activity and downstream targets, suppressing tumor growth and metastasis in vivo.

Together, these findings identify stem-like LumHR cells, their spatial niches, and FOXA1-driven programs as central features of aggressive DCIS and as potential therapeutic vulnerabilities.

## Results

### Sample Cohort and Analytical Framework

To characterize cellular and spatial features associated with invasive potential in DCIS, we analyzed 43 patient-derived samples using complementary single-cell and spatial profiling approaches: 17 by single-cell RNA sequencing (scRNA-seq), 16 by spatial transcriptomics on the 10x Genomics Xenium platform, and 10 by joint single-cell ATAC and RNA sequencing (scATAC/RNA-seq). **Figure 1a** summarizes the study design and the relationships among analysis type, progression status, sample type, and molecular subtype.

**Figure 1.**
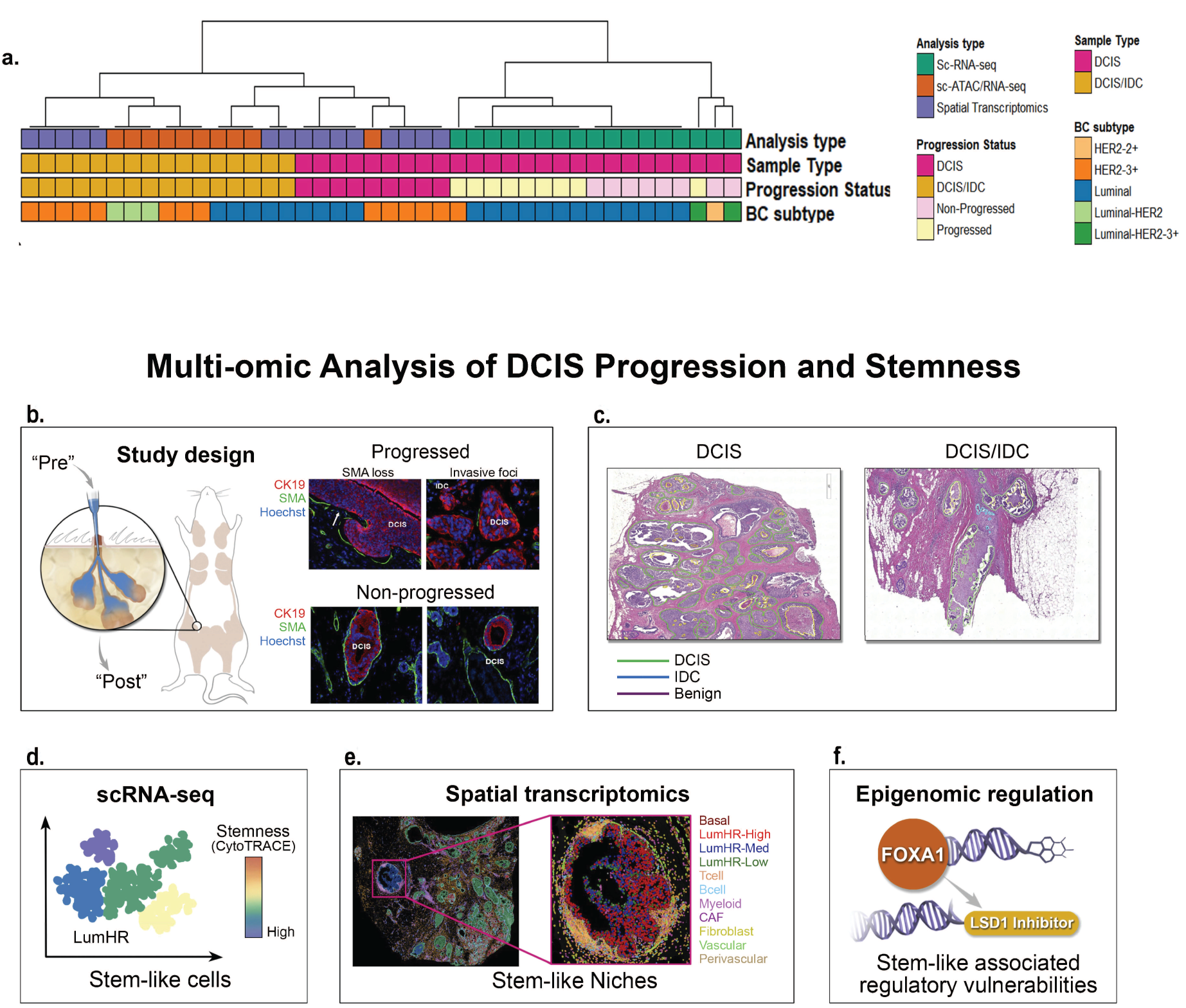
Multi-omic analysis of DCIS progression and stemness. **a,** Overview of the cohort and multi-omic profiling strategy. A total of 43 DCIS samples were subjected to scRNA-seq, scATAC/RNA-seq, or spatial transcriptomics. Annotation tracks indicate analysis type, sample type (DCIS or DCIS/IDC), progression status (progressed or non-progressed), and breast cancer subtype. **b,** Study design for intraductal (MIND) transplantation experiments. Pure DCIS lesions were profiled prior to transplantation (Pre) and again 9 months post-transplantation (Post) to enable longitudinal analysis of molecular programs associated with invasive progression. **c,** Histologic context of analyzed lesions, illustrating pure DCIS and DCIS with associated invasive ductal carcinoma (IDC). **d,** scRNA-seq analysis of DCIS epithelial cells identifying stem-like populations, with stemness inferred using CytoTRACE scores. **e,** Spatial transcriptomics of pure DCIS and DCIS/IDC localizes stem-like cells and defines stem-like niches, including their cellular composition and cell–cell interactions. Representative stemness-associated markers are shown. **f,** Integrated epigenomic analysis identifies stemness-associated regulatory vulnerabilities. scATAC/RNA-seq highlights transcriptional regulators of stemness, and pharmacologic inhibition of the stemness-associated regulator LSD1 suppresses metastatic progression in HER2+ DCIS models.

The 17 scRNA-seq samples were evaluated for invasive progression using the Mouse INtraDuctal (MIND) model, in which primary human DCIS epithelial cells are transplanted directly into mouse mammary ducts to enable longitudinal assessment in an anatomically relevant microenvironment. Of these, nine samples exhibited invasive progression, whereas eight remained non-invasive over a nine-month follow-up period (**Figure 1b**). We and others have shown that this model faithfully recapitulates the natural progression of untreated patient-derived DCIS, preserves the biomarker expression profiles and histopathologic features of the corresponding patient lesions, and reflects breast cancer metastatic patterns more accurately than conventional cleared fat pad transplantation models ^13–15^. The invasion rate in our MIND models (∼46%) closely matches the reported progression rate of untreated moderate-grade DCIS in patients ^5^.

Samples were profiled by scRNA-seq before transplantation and again at 9 months, allowing us to track how epithelial and stromal programs, including stemness-associated features, evolved in progressing versus non-progressing lesions (**Figure 1b**,**d**). Spatial transcriptomics defined the distribution of stem-like cells within DCIS lesions, characterized their local microenvironments, and identified ligand–receptor interactions enriched in stemness-associated niches (**Figure 1c,e**). Joint scATAC/RNA-seq linked transcriptional states to chromatin accessibility in single-cells, enabling identification of high-stemness cell populations and their regulatory features (**Figure 1f**). Analyses focused on comparing HER2^+^ and luminal DCIS subtypes, given the association of HER2^+^ with more aggressive clinical behavior ^11,16,17^. Sample-level details, including biomarker status, histologic and nuclear grade, patient demographics, and race/ethnicity, are provided in **Supplementary Table 1.**

### Integrated Single-cell and Spatial Analysis Delineates Epithelial Cell States in DCIS and DCIS/IDC

We annotated cell types in our DCIS datasets using the Human Breast Cell Atlas (HBCA) from Kumar and colleagues, who profiled 714,331 cells and 117,346 nuclei across 220 tissue samples from 132 women using single-cell transcriptomics and spatial proteomics to define the epithelial, stromal, and immune cell types of the adult human breast ^18^. **Supplementary Table 2a** lists the Xenium samples with their subtypes and metastasis status, and **Supplementary Table 2b** catalogues the 380 genes in the Xenium panel.

We first transferred the HBCA major cell type labels onto each Xenium sample, then integrated 2.6 million cells from all 16 samples to build a reference map of cell types in patient DCIS and DCIS/IDC samples (**Figure 2a**). Label assignments were validated against the marker genes reported by Kumar et al. (**Supplementary Table 2c**), with concordant expression patterns confirmed in **Figure 2b**. A pathologist reviewed all 16 H&E images and annotated benign, DCIS, and IDC epithelium (**Supplementary Figure 1**). Unbiased sub-clustering of the epithelial compartment yielded fifteen distinct subclusters (**Figure 2c–f**, **Supplementary Table 2d**). Each subcluster retained its Basal, LumSec, or LumHR identity and was further classified as benign-associated, DCIS-associated, or non-specific (present in both regions), based on its frequency across HER2^+^ versus Luminal subtypes (**Figure 2d,e**) and across pathologist-annotated regions (**Supplementary Figure 1**). **Supplementary Figure 1** provides a comprehensive view of the 16 Xenium samples: pathologist-annotated regions (a), proportional distribution of epithelial subclusters per region (b), their spatial organization (c), and a colocalization-based graph of inter-subcluster interactions (d). **Figure 2f** is a heatmap of pathways and molecular programs enriched within each epithelial cell subcluster.

**Figure 2.**
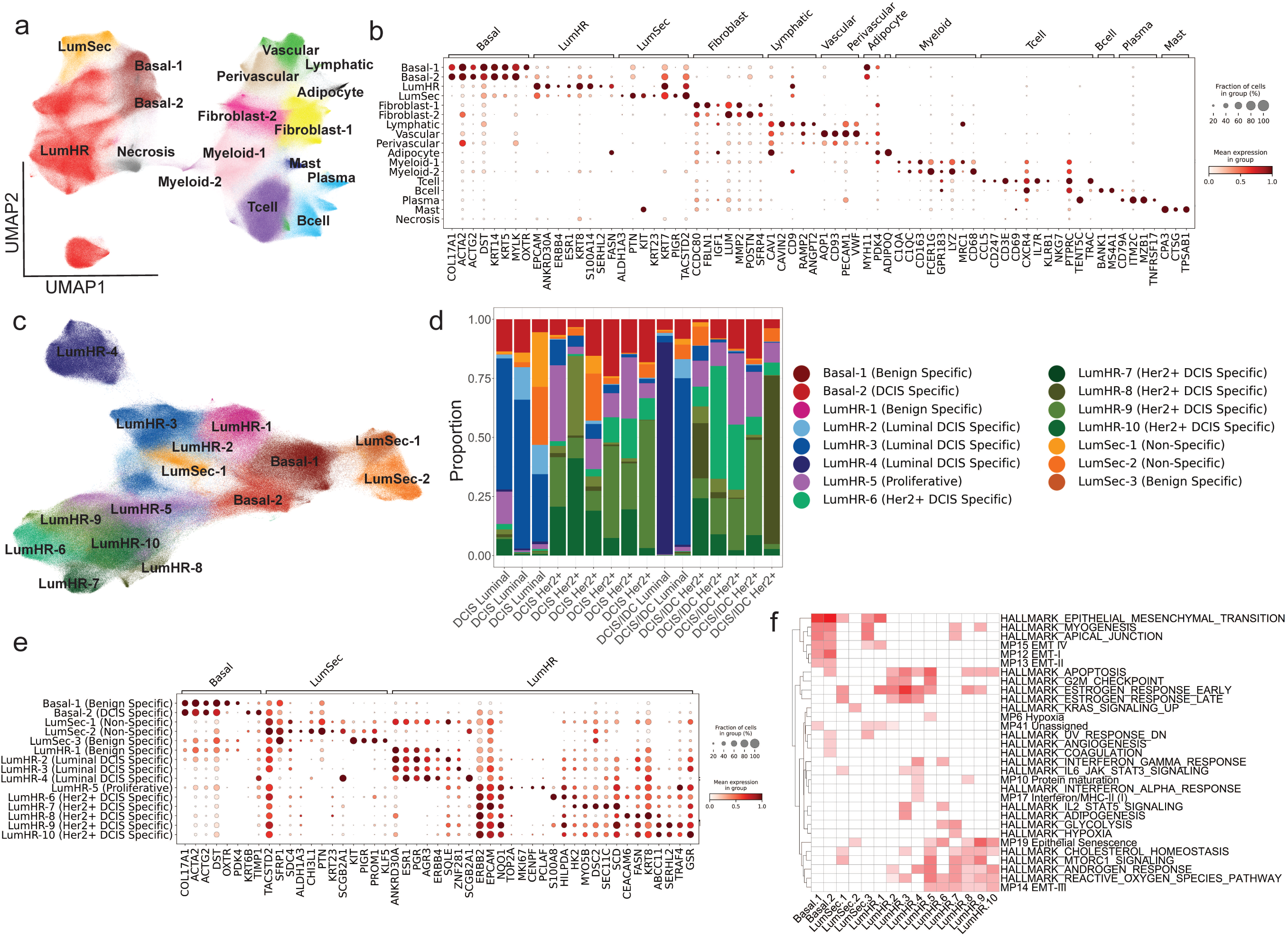
Spatially resolved epithelial reprogramming during DCIS progression. **a,** UMAP of 2.6 million cells integrated from 16 Xenium samples. Major cell types were annotated by label transfer from the Human Breast Cell Atlas (HBCA) reference dataset published by Kumar *et al*. **b,** Dot plot showing expression of canonical marker genes used to validate transferred HBCA cell-type annotations. **c,** UMAP of epithelial cells only, revealing transcriptionally distinct epithelial subclusters. **d,** Relative frequencies of epithelial subclusters across samples stratified by molecular subtype (Luminal versus HER2⁺). **e,** Dot plot of marker genes defining epithelial subclusters shown in **d**. **f,** Heatmap of pathway and molecular program enrichment based on subcluster-specific marker genes. White indicates no significant enrichment; red indicates significant enrichment, with darker shades representing increased gene overlap.

### Functional Reprogramming of Basal Myoepithelial Cells in DCIS Reveals Loss of Physiologic Signaling and Gain of Tumor-Promoting Phenotypes

We identified two distinct basal subclusters: Basal-1, associated with benign epithelium, and Basal-2, associated with DCIS lesions (**Figure 3b,c,f,g**, **& Supplementary Figure 1**). Both expressed classic basal/myoepithelial markers including *COL17A1*, *ACTA2*, *ACTG2*, and *DST*. Basal-1 additionally expressed *OXTR*, which mediates milk ejection during lactation ^19^, and *PDK4*, which regulates the switch between glucose and fatty acid utilization during late pregnancy and lactation ^20^. Both genes were significantly downregulated in Basal-2, suggesting a loss of normal myoepithelial physiological function during the transition to DCIS (**Figure 2e**).

**Figure 3.**
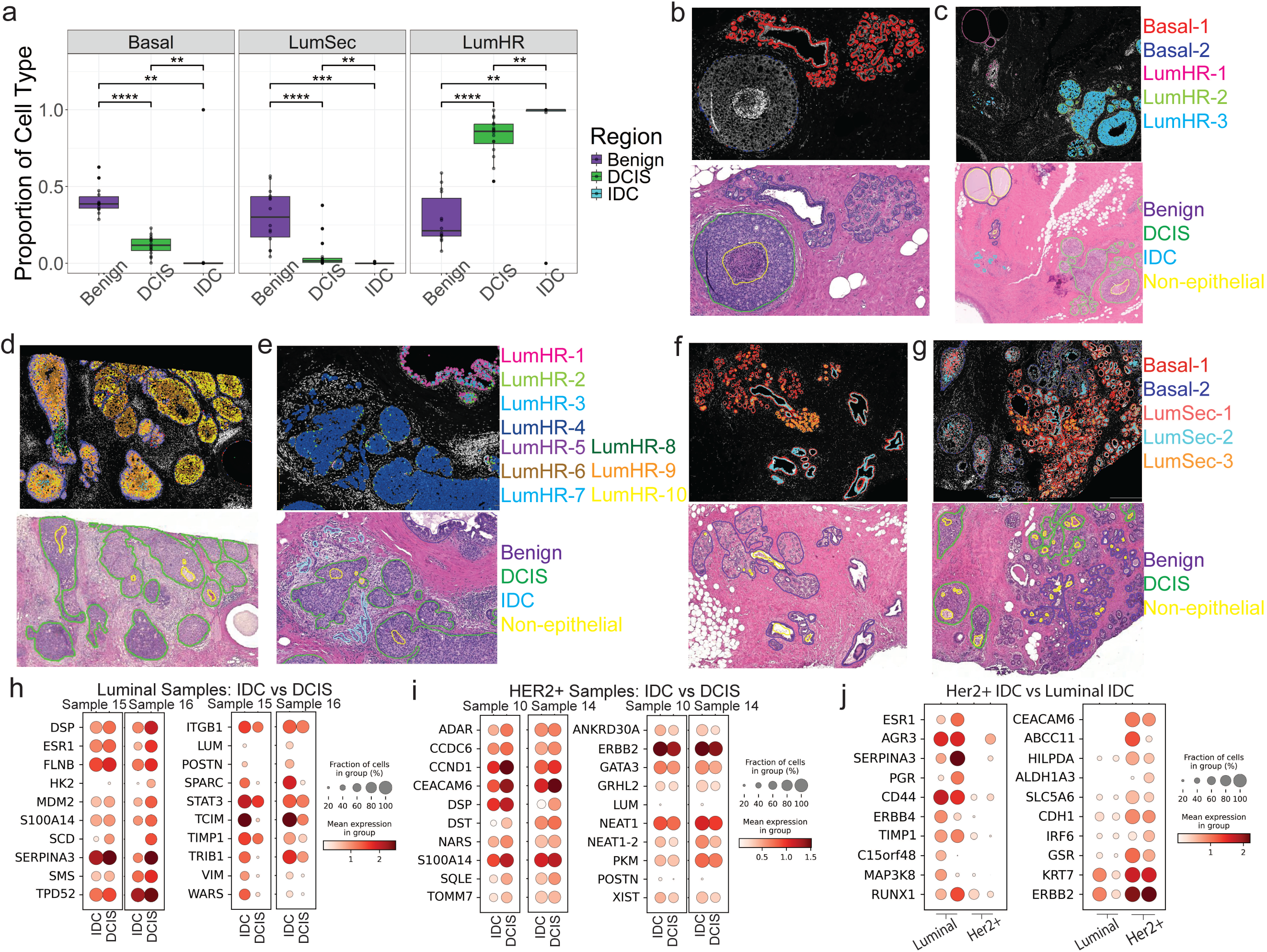
Single-Cell Resolution of DCIS Identifies LumHR Expansion, Basal Dysfunction, and Subtype-Specific Pathways to Invasion. **a,** Box plots showing proportions of Basal, luminal secretory (LumSec) and luminal hormone-responsive (LumHR) cells across benign, DCIS and IDC regions for each Xenium sample. **b,** Spatial organization of Basal-1 and Basal-2 subclusters **c,** Spatial organization of LumHR-1, LumHR-2, and LumHR-3 subclusters **d,** Spatial organization of LumHR-5, LumHR-6, LumHR-7, LumHR-8, LumHR-9, and LumHR-10 **e,** Spatial organization of LumHR-1, LumHR-2, LumHR-3, and LumHR-4 **f,** Spatial organization of Basal-1, LumSec-2, and LumSec-3 **g,** Spatial organization of Basal-1, Basal-2, LumSec-1, LumSec-2, and LumSec-3 **h-j,** Differential gene expression analysis of luminal hormone-responsive (LumHR) subclusters comparing invasive ductal carcinoma (IDC) and ductal carcinoma in situ (DCIS) regions within Luminal and HER2⁺ samples (**h,i**), and between Luminal and HER2⁺ IDC regions (**j**), to identify subtype-specific and shared transcriptional programs associated with progression from DCIS to IDC.

Conversely, Basal-2 showed significantly higher expression of cancer-related genes such as *TIMP1*, *EPCAM*, and *TACSTD2* (*TROP-2*) ^21,22^. EPCAM expression in DCIS basal myoepithelial cells has been linked to high recurrence risk ^10^ (**Figure 2e**). Hallmark pathways ^23^ showed that Basal-2 (DCIS-associated) specific genes mapped to pathways such as, apoptosis (*TIMP1*, *LUM*, *CAV1*, *TIMP2*), angiogenesis (*TIMP1*, *LUM*, *POSTN*), and coagulation (*SPARC*, *TIMP1*, *ITGB3*) (**Figure 2f**). Apoptosis-related gene expression in Basal-2 cells may indicate a constant pressure to evade cell death during the DCIS-to-IDC transition. Although *SPARC*, *TIMP1* and *ITGB3* are canonically linked to coagulation in the basal compartment surrounding DCIS, they likely promote extracellular matrix remodeling, angiogenesis and immune modulation ^24–26^.

### LumSec Heterogeneity in DCIS

The luminal secretory (LumSec) compartment resolved into three subclusters. LumSec-3 was largely confined to benign lobular regions, while LumSec-1 and LumSec-2 were present in both benign and DCIS lesions (**Figure 3f,g**, **& Supplementary Figure 1**). LumSec-3 expressed high levels of mammary progenitor markers *KIT* ^27^ and *PROM1*, along with *PIGR* (which transports antibodies from the bloodstream to milk) and *KLF5* (a PR target gene driving mammary gland remodeling during pregnancy and lactation) ^28^ (**Figure 2e**).

LumSec-1 showed high expression of *ER* and ER target genes (*PGR*, *ANKRD30A*, *AGR3*), consistent with ER-positive, lower-grade tumors ^29^ (**Figure 2e**). In contrast, LumSec-2 expressed *PTN*, *CHI3L1*, and *SCGB2A*, factors linked to immunosuppression via neutrophil recruitment and T cell inhibition ^30^ (**Figure 2e**). LumSec-2 was also enriched for KRAS signaling and epithelial senescence, supporting a more aggressive phenotype (**Figure 2f**). Expansion of LumSec-1 and LumSec-2 within DCIS was observed in only one Luminal sample (sample 9); these subclusters did not contribute substantially to any other DCIS samples (**Supplementary Figure 1**).

### LumHR Subclusters Highlight Distinct Pathways of DCIS Progression and Transition to Invasive Disease

We identified ten LumHR subclusters (LumHR-1 through LumHR-10) (**Figure 2c-f**). LumHR-1 was associated with benign regions and as expected, expressed luminal epithelial markers (*ANKRD30A*, *ESR1*, *PGR*, *AGR3*, *ERBB4*) with low or absent expression of cancer-related genes such as *SQLE*, *ERBB2*, and *NQO1* (**Figure 2e**). LumHR-2 through LumHR-4 were specific to Luminal DCIS (**Figure 3d-e**; **Supplementary Tables 2d-e, & Supplementary Figure 1**), with subcluster-specific genes mapping to apoptosis (largely anti-apoptotic), G2M checkpoint (except LumHR-4), and early and late estrogen response hallmarks (**Figure 2f**).

LumHR-5 was a proliferative subcluster marked by cell cycle genes (*TOP2A*, *MKI67*, *CENPF*, *PCLAF*) and was predominantly localized to the basal regions of DCIS (**Figure 3d**; **& Supplementary Figure 1**). LumHR-6 through LumHR-10 were unique to HER2^+^ samples (**Figure 2c-e**; **& Supplementary Tables 2d,e**), with enriched genes mapping to androgen response, mTORC1 signaling, reactive oxygen species response, MP19 epithelial senescence, and cholesterol homeostasis - pathways whose interplay has been linked to aggressive cancer phenotypes (**Figure 2f**). For example, LumHR-6-10 showed unique high expression of *NQO1 (***Figure 2e**), an enzyme that protects cells from oxidative stress ^31^ and is frequently elevated in tumors. This represents a potential therapeutic vulnerability exploitable by bioactivatable drugs such as deoxynyboquinone (DNQ), which NQO1 converts into a potent cytotoxin. The MP19 epithelial senescence signature in this group is also clinically relevant, as senescence may worsen prognosis by impairing tissue repair and driving chronic inflammation through the senescence-associated secretory phenotype (SASP) ^23,32^ (**Figure 2f**).

### Progressive Enrichment of LumHR Cells Drives the Transition from Benign and DCIS to IDC

We quantified the proportion of Basal, LumSec, and LumHR cells within pathologist-annotated region (benign, DCIS, or IDC). The proportions of Basal and LumSec subclusters decreased significantly from benign tissue to DCIS (q-value < 0.0001) and further to IDC (q-value < 0.01), whereas LumHR cells showed a corresponding increase (q-value < 0.0001 & q-value < 0.01) (**Figure 3a**). These findings indicate that LumHR subclusters represent the predominant epithelial population in DCIS, with further expansion during progression to IDC. The reduction in Basal and LumSec populations suggest a loss of structural and secretory epithelial functions, whereas the expansion of LumHR cells reflect the emergence of a population with enhanced proliferative capacity, survival advantage, and stem-like features. These cells are characterized by activation of hormone signaling and chromatin remodeling programs that may promote cellular plasticity and invasive behavior. Together, the progressive enrichment of LumHR cells from benign tissue through DCIS to IDC positions this population as a central contributor to disease progression and highlights potential therapeutic vulnerabilities associated with this state.

### Subtype-Specific Transcriptional Programs Underlying LumHR-Driven Progression from DCIS to IDC

Given the dominant role of LumHR cells in DCIS to IDC progression, we identified differentially expressed genes (DEGs) between LumHR cells in IDC vs. DCIS across Luminal and HER2⁺ subtypes. In luminal samples, IDC-enriched genes (*ITGB1*, *LUM*, *POSTN*, *SPARC*, *STAT3*, *TCIM*, *TIMP1*, *TRIB1*, *VIM* and *WARS*) reflect ECM remodeling, EMT, and pro-survival signaling (**Figure 3h**, & **Supplementary Table 2f**). HER2⁺ IDC showed enrichment of (*ANKRD30A*, *ERBB2*, *GATA3*, *GRHL2*, *LUM*, *NEAT1_2*, *PKM*, *POSTN*, and *XIST*) indicating luminal identity coupled with metabolic, chromatin, and stress-adaptive programs driving aggressive progression (**Figure 3i**, **& Supplementary Table 2f**).

Comparison of IDC regions in HER2^+^ vs. Luminal subtypes revealed HER2⁺-specific upregulation of *CEACAM6* (a cell adhesion molecule), *ABCC11* (multidrug resistance gene), *HILPDA* (lipid storage and cancer cell survival), *ALDH1A3* (a biomarker of stemness and biomarker of poor prognosis in cancer), *SLC5A6* (sodium-dependant multivitamin transporter), *CDH1* (cadherin-1), *IRF6* (interferon regulatory factor 6), *GSR* (glutathione reductase), *KRT7* (Keratin-7) and *ERBB2* (HER2), consistent with invasion, metabolic flexibility, and therapy resistance ^33^ (**Figure 3j**, **& Supplementary Table 2g**).

In contrast, Luminal IDC retained hormone-driven signaling (*ESR1* (estorgen receptor), *AGR3* (anterior gradient 3), *SERPINA3* (a protease inhibitor), *PGR* (progesterone receptor), *CD44* (cell adhesion), *ERBB4* (HER4), *TIMP1* (tissue inhibitor of metalloproteinase 1), *C15orf48* (chromosome open reading frame 48), *MAP3K8* (mitogen-activated protein kinase kinase kinase 8) and *RUNX1* (transcription factor)), with contributions from ECM remodeling and inflammatory signaling ^34^ (**Figure 3j**, **& Supplementary Table 2g**).

Together, these findings define distinct subtype-specific progression programs, in which HER2⁺ IDC is characterized by metabolically adaptive, therapy-resistant phenotypes, whereas Luminal IDC retains hormonally driven and stromally regulated features.

### Cellular Plasticity and Stemness in LumHR Cells Drive DCIS-to-IDC Progression

Cellular plasticity and stem-like characteristics are thought to drive invasion and metastasis, enabling cancer cells to adapt, survive, and acquire migratory capacity ^35–37^. To test whether cellular stemness underlies DCIS invasive potential, we applied CytoTRACE ^12^ - a method for predicting cellular stemness from scRNA-Seq data from 17 MIND model samples (9 progressors and 8 non-progressors) profiled before transplantation ("Pre") and at nine months ("Post"). scRNA-seq of 68,000 cells recovered the expected epithelial and stromal populations, classified using Kumar et al. markers (**Figure 4a–b**). Among these, LumHR cells emerged as the dominant expanding population and carried the highest stemness scores of any epithelial or stromal subset, in both scRNA-seq and spatial transcriptomics datasets (**Supplementary Figure 2a,b**) immediately implicating them as the cancer-driving compartment.

**Figure 4.**
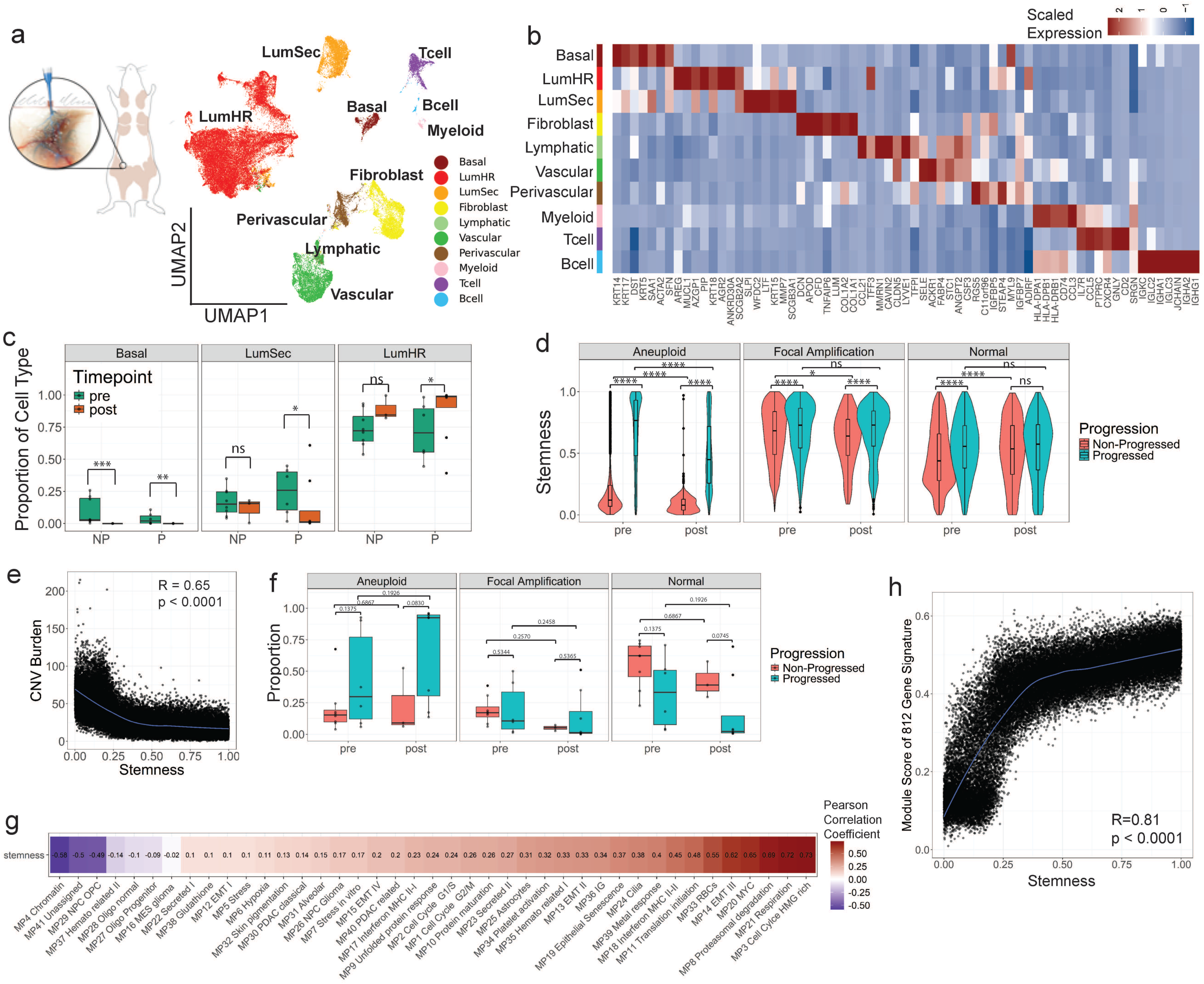
Stemness of luminal hormone-responsive cells predicts invasive progression in the MIND model. **a,** UMAP representation of 68,000 cells profiled by single-cell RNA sequencing from Mouse INtraDuctal (MIND) model samples. Major cell types were annotated by label transfer from the Human Breast Cell Atlas (HBCA) reference dataset published by Kumar *et al*. **b,** Dot plot showing expression of canonical marker genes used to validate transferred HBCA cell-type annotations. **c,** Changes in epithelial cell-type composition across pre- and post-transplantation time points, stratified by progression status. Basal and luminal secretory (LumSec) populations decrease following transplantation, whereas luminal hormone-responsive (LumHR) cells expand significantly, particularly in progressor samples. **d,** CytoTRACE stemness scores of LumHR cells split by CNV classification and across time points and progression status, showing significantly higher stemness in progressor samples both before and after transplantation. **e,** Spearman correlation between CytoTRACE stemness scores and inferred copy number variation (CNV) burden in LumHR cells, revealing a negative association. **f,** Classification of LumHR cells by CNV status (normal, focally amplified or aneuploid) across time points and progression groups. **g,** Correlation between stemness scores and cancer cell meta-programs, showing strong positive associations with chromatin organization (MP3), respiration (MP21), proteasomal degradation (MP8), MYC targets (MP20) and epithelial–mesenchymal transition (MP14). **h,** Spearman correlation between CytoTRACE stemness scores and module score of the 812-gene classifier from Strand *et al*.

Tracking compositional shifts across timepoints reinforced this conclusion (**Figure 4c**). Basal cells were absent post-transplantation regardless of outcome. LumSec cells declined from Pre to Post, with a significant reduction in progressors (q < 0.05). LumHR cells, by contrast, expanded dramatically in post-transplant progressor samples, which were composed almost exclusively of LumHR cells (q < 0.05, mirroring the spatial transcriptomics trend (**Figure 3a**) and identifying LumHR cells as the primary drivers of invasive progression.

To dissect the genomic basis of this expansion, we classified epithelial cells as normal, focally amplified, or aneuploid using InferCNV ^38^ and assessed stemness in LumHR cells across CNV classes (**Figure 4d**). Stemness scores were significantly higher in progressors than non-progressors at both timepoints and across all three CNV classes, with the largest gap in aneuploid cells. Critically, however, stemness scores in LumHR cells were *negatively* correlated with overall CNV burden (**Figure 4e**) and with both copy number gains and losses individually (**Supplementary Figure 2c–d**). Thus, aneuploidy alone is insufficient to drive progression; it is the *combination* of chromosomal instability with stem-like state that fuels the transition to invasion.

CNV classification supported this: post-progressor LumHR cells trended toward higher aneuploid (q = 0.083) and lower normal (q = 0.075) proportions than post-non-progressors (**Figure 4f**).

To define the cellular programs underlying LumHR stemness, we correlated stemness scores with the 41 cancer meta-programs (MPs) defined across 1,163 tumors and 24 cancer types by Gavish et al. ^23^. LumHR stemness was strongly positively correlated with chromatin organization (MP3, Cell Cycle HMG-rich), respiration (MP21), proteasomal degradation (MP8), MYC targets (MP20), and epithelial-mesenchymal transition (MP14) (**Figure 4g**) - a coordinated program coupling chromatin structure, metabolic, and EMT machinery that plausibly equips LumHR cells with the adaptability required for invasion.

Together, these data establish LumHR cells as the dominant, stemness-enriched compartment expanding through DCIS-to-IDC progression, with their invasive potential driven not by genomic instability per se but by the convergence of stem-like state with aneuploidy and a coordinated chromatin-EMT-MYC program.

Notably, stemness was decoupled from proliferation, as correlations with the cell-cycle meta-programs MP1 (G2M) and MP2 (G1/S) were weak or negative, as were correlations with stress-associated programs MP5, MP6 (hypoxia), and MP7 (**Figure 4g**). Thus, the LumHR cells driving invasive progression are defined by a stem-like state rather than active cycling or stress response programs.

To establish clinical relevance, we next asked whether these stemness programs are associated with DCIS recurrence after treatment. Stemness scores correlated significantly with expression of the 812-gene molecular classifier of DCIS recurrence ^11^ (**Figure 4h**), extending our findings beyond untreated progression and indicating that elevated stemness also predicts post-treatment recurrence.

Collectively, these findings establish LumHR cells as principal drivers of DCIS-to-IDC progression and identify elevated stemness, decoupled from proliferation and independent of extensive chromosomal instability, as the defining, clinically actionable feature underlying their invasive potential.

### Stem-Like Cell Niches as Microenvironmental Markers of High-Risk DCIS

We hypothesized that stem-like epithelial cells occupy discrete spatial niches that reinforce stemness through coordinated cell–cell interactions, and that these niches, if detectable within histologically pure DCIS, could serve as microenvironmental markers of invasive risk. To test this, we used spatially constrained clustering to define stemness-enriched niches in DCIS lesions and characterize their cellular composition and signaling architecture.

We first sub-clustered immune and stromal compartments from **Figure 2a**, resolving 14 immune cell types (myeloid and lymphoid subsets; **Supplementary Figure 3a–e, j**) and 9 stromal cell types (vascular, perivascular, lymphatic, and fibroblast subsets; **Supplementary Figure 3f–i, k**). Nearest-neighbor analysis across all samples revealed a consistent microenvironmental shift along the benign-to-IDC axis: Fibroblasts and Vascular-1 cells were closest to benign epithelium, while CAFs-1, Perivascular-2, and M1 macrophages dominated the immediate neighborhood of DCIS and IDC, a pattern preserved across HER2^+^ DCIS, HER2^+^ DCIS/IDC, and Luminal samples (**Supplementary Figures 4–6**).

Using the cross-type pair correlation function (*g*(*r*); **Supplementary Figure 7**) we confirmed and extended these distance findings. Fibroblasts and Vascular-1 cells showed strong clustering with benign epithelium at short ranges (∼50–100 µm) but were dispersed from DCIS and IDC (**Supplementary Figure 7a, g**). CAFs displayed the inverse pattern, clustering with cancer cells - most strongly in IDC - and in some samples actively dispersing from benign epithelium (**Supplementary Figure 7b–c**). Macrophage subsets segregated similarly: M1 macrophages clustered with cancer cells (more so in IDC), while M2 macrophages clustered with benign epithelium at short range, particularly in HER2+ samples (**Supplementary Figure 7j–k**).

Against this rewired microenvironment, spatially constrained clustering identified discrete stem-like niches in HER2^+^ samples, in which individual DCIS lesions or IDC regions formed distinct niches separated from neighboring lesions in the same sample (**Figure 5a,e**). In the first HER2^+^ DCIS sample (**Figure 5a**), niche-8 consisted entirely of LumHR cells and exhibited the highest stemness scores in the sample, exceeding those observed in LumHR cells from other DCIS lesions (niche-2) and benign epithelium (niche-0) (**Figure 5b,c**). Differential gene expression analysis identified CEACAM6 as the defining marker of niche-8 (log2FC = 3.71; 98.4% vs. 39.4% of LumHR cells; q < 0.0001), with concurrent elevation of FOXA1 (**Figure 5d**). A HER2^+^ DCIS/IDC sample displayed a similar architecture: niche-1, an IDC region composed almost entirely of LumHR cells, exhibited the highest stemness scores (**Figure 5e–g**) and was distinguished from the adjacent DCIS lesions (niche-6) by elevated FOXA1. Although CEACAM6 expression was also increased in niche-6, the proportion of CEACAM6-postivive cells was comparable between niches (97.9% vs 96.8%, **Figure 5h**), suggesting that spatial regulation of expression level rather than frequency distinguishes these niches.

**Figure 5.**
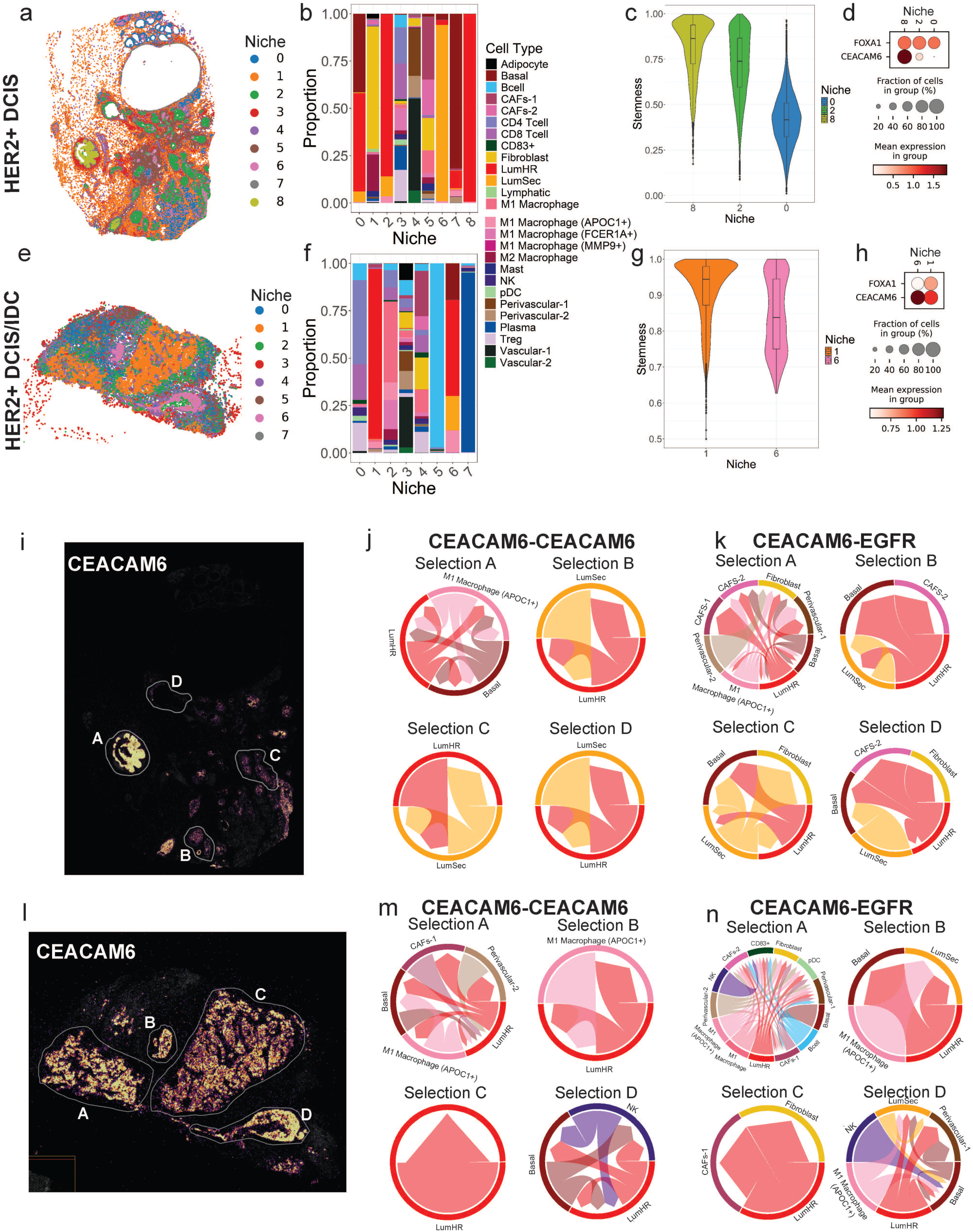
Spatial Mapping of DCIS Identifies Discrete Stem-Like Niches with Rewired Microenvironments and Invasive Potential. **a,** Image of Sample 3 (HER2+ DCIS) with cells colored in by identified niches. **b,** Bar plot showing the proportion of each cell type in each niche. **c,** Violin plot showing the stemness scores of LumHR cells in niche-8, niche-2, and niche-0, the three epithelial niches in Sample 3. **d,** Dot plot showing the expression of FOXA1 and CEACAM6 in LumHR cells in niche-8, niche-2, and niche-0 of Sample 3. **e,** Image of Sample 14 (HER2+ DCIS/IDC) with cells colored in by identified niches. **f,** Bar plot showing the proportion of each cell type in each niche. **g,** Violin plot showing the stemness scores of LumHR cells in niche-1 and niche-6, the two epithelial niches in Sample 14. **h,** Dot plot showing the expression of FOXA1 and CEACAM6 in LumHR cells in niche-1 and niche-6 of Sample 14. **i,** CEACAM6 expression in Sample 3 with selected regions for LR CCI analysis marked by the boundaries of the white lines. Four regions were chosen denoted A, B, C, and D. **j,** Chord plot showing the significantly interacting cell types for the CEACAM6-CEACAM6 LR pair in each selection from **i**. **k,** Chord plot showing the significantly interacting cell types for the CEACAM6-EGFR LR pair in each selection from **i**. **l,** CEACAM6 expression in Sample 14 with selected regions for LR CCI analysis marked by the boundaries of the white lines. Four regions were chosen denoted A, B, C, and D. **m,** Chord plot showing the significantly interacting cell types for the CEACAM6-CEACAM6 LR pair in each selection from **l**. **n,** Chord plot showing the significantly interacting cell types for the CEACAM6-EGFR LR pair in each selection from **l.** *For chord plots the base of the arrow represents the sender cell type and the tip of the arrow represents the receiver cell type. With the sender expressing the first gene denoted in the LR pair, and the receiver expressing the second gene denoted in the LR pair.

To probe how these stem-like niches engage the microenvironment, we performed ligand–receptor (LR) analysis on *CEACAM6*. Two interactions emerged as significant across niches: CEACAM6–CEACAM6 and *CEACAM6*–*EGFR* (**Figure 5j-n**). In both samples, the high-stemness niches harbored substantially more significant LR engagements than lower-stemness regions (**Figure 5i–n**). Notably, *CEACAM6*–*EGFR* ligand–receptor interactions were enriched between LumHR and basal epithelial cells in pure DCIS and multiple stromal populations, including CAFs, perivascular cells, and *APOC1+* macrophages (**Figure 5i,k**; **Selection A**). CAFs and perivascular cells are well-established mediators of tumor progression and immunosuppression, contributing to extracellular matrix remodeling, angiogenesis, and malignant cell invasion ^39,40^. In parallel, APOC1+ macrophages represent a specialized subset of tumor-associated macrophages (TAMs) that predominantly exhibit an M2-like, pro-tumorigenic phenotype^41^. Collectively, these findings indicate that the tumor microenvironment (TME) surrounding high-stemness niches is markedly more pro-tumorigenic and immunosuppressive than that of low-stemness niches. Additional interactions between LumHR and basal cells with NK cells, plasmacytoid dendritic cells, and CD83+ immune cells in DCIS/IDC niches suggest enhanced recruitment of tumor-sensing immune populations to high-stemness regions that have progressed to an invasive state (**Figure 5n**; **Selection A**). Similarly, *CEACAM6*–*CEACAM6* ligand–receptor interactions were enriched within high-stemness niches in both DCIS and DCIS/IDC regions, involving LumHR and basal epithelial cells interacting with *APOC1+* macrophages, perivascular cells, and CAFs (**Figure 5 j,m**; **Selection A**). These findings further support a role for CEACAM6-mediated signaling in establishing and maintaining immunosuppressive, pro-tumorigenic microenvironments specifically within high-stemness niches. *CEACAM6* encodes a GPI-anchored cell adhesion protein associated with poor prognosis and metastatic potential in multiple cancers ^42^. Importantly, therapeutic antibodies targeting CEACAM6 are now in development, offering a translational opportunity to disrupt stemness-driven invasive progression ^43^.

Niches analysis was performed in all samples (Supplementary Figure 8). CEACAM6 was not consistently expressed in all HER2+ samples; however, in the samples that did express it (**Supplementary Figure 8a, d, f, g, h**), it was increased in the niches with the highest stemness.

CEACAM6 was not expressed in Luminal samples. FOXA1 tends to be expressed in LumHR cells niches with the highest stemness.

Together, these findings show that stem-like niches are spatially discrete, transcriptionally distinct, and engaged in active CEACAM6–EGFR signaling with the surrounding microenvironment. Their presence within histologically uniform DCIS lesions positions them as candidate microenvironmental markers of high-risk disease lesions already primed for invasive progression.

### Targeting FOXA1 to Disrupt Stemness and Halt DCIS Progression

To investigate epigenetic regulators of stemness in DCIS, we performed integrated single-nucleus multiome (ATAC-seq and RNA-seq) profiling on epithelial cells from ten patient-derived DCIS and DCIS/IDC samples (**Supplementary Table 1; & Supplementary Figure 9a**). Within each subtype, cells were clustered and ranked by stemness score (**Figure 6a**; **& Supplementary Figure 9c,d**), and the highest-stemness cluster (Cluster 1) was composed predominantly of LumHR cells (**Supplementary Figure 9b**). Transcription factor motif enrichment analysis of open chromatin in Cluster 1 identified strong, consistent enrichment of FOXA1 motifs across all DCIS subtypes (**Figure 6b**), implicating FOXA1 as a direct transcriptional regulator of stemness. Subtype-specific co-enrichment was also observed: GRHL2, AP-2, and AP2 in HER2+ DCIS/IDC, and AP-1 in pure HER2+ DCIS - suggesting that cooperative transcription factor binding shapes subtype-specific stemness programs and malignant progression.

**Figure 6.**
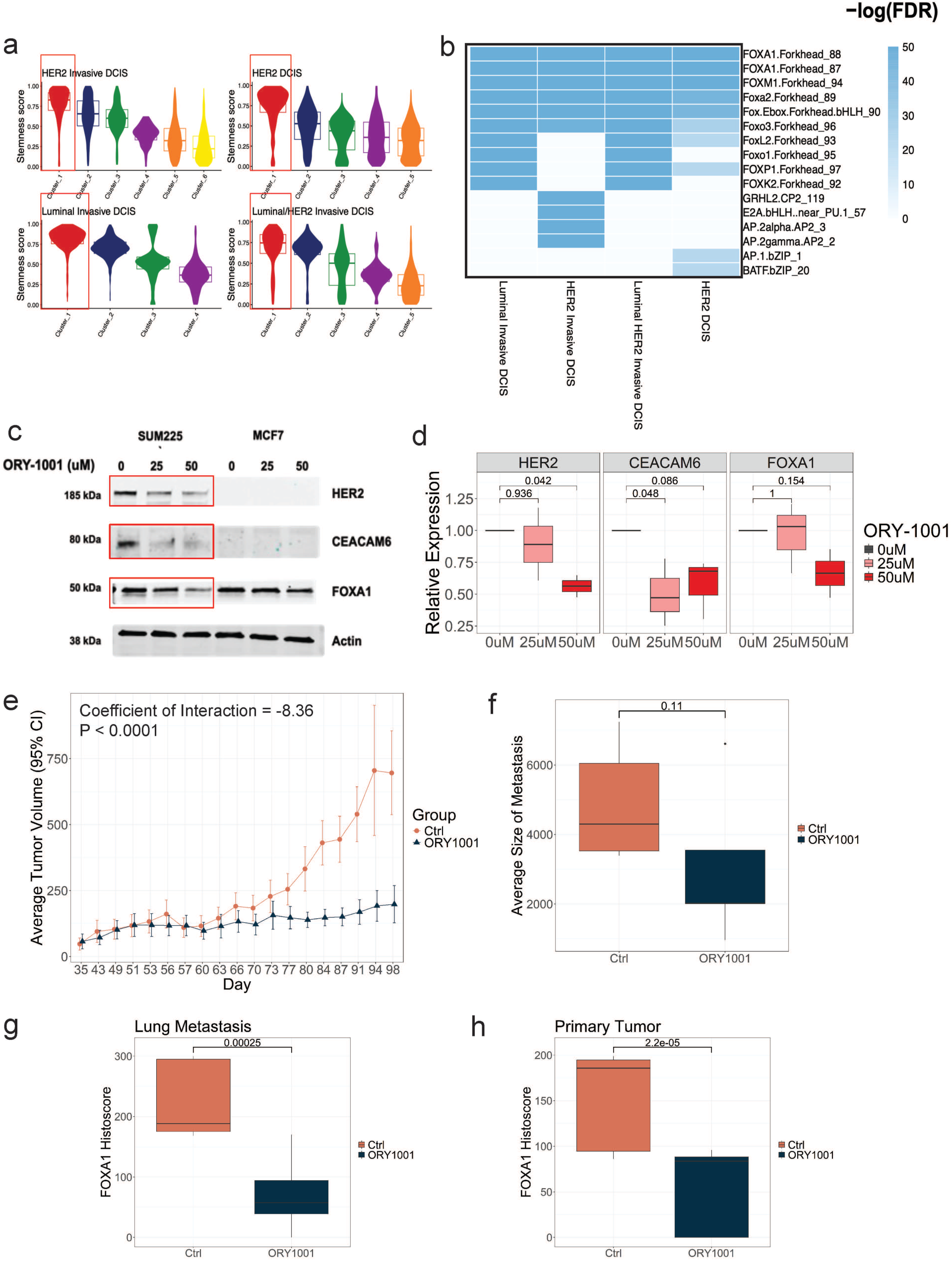
FOXA1-Driven Stemness Programs in DCIS Are Epigenetically Regulated by LSD1 and Therapeutically Targetable. **a,** Violin plot showing the clusters with the highest stemness in snATAC/RNA-seq in each subtype **b,** Top significant binding motifs of stemness-high clusters in each subtype **c,** Representative western blot showing in SUM225, ORY-1001 reduced FOXA1, HER2, and CEACAM6 expression **d,** Boxplot showing change in HER2, CEACAM6, and FOXA1 expression by densitometric analysis for three replicates of the western blot. **e,** Average tumor volume over time in control mice (orange) and ORY-1001 (dark blue) treated mice **f,** Average size of metastasis in control mice (orange) and ORY-1001 (dark blue) treated mice **g-h,** FOXA1 histoscore in lung metastases (**g**) and primary tumor (**h**) of control mice (orange) and ORY-1001 (dark blue) treated mice

To therapeutically target FOXA1, we exploited a recently described regulatory mechanism in which LSD1 (KDM1A) directly demethylates FOXA1 at lysine 270, adjacent to its DNA-binding domain, enhancing FOXA1 chromatin occupancy independently of LSD1’s canonical H3K4 demethylase activity ^44^. Pharmacologic or genetic LSD1 inhibition rapidly displaces FOXA1 from chromatin, reduces accessibility, suppresses FOXA1-dependent transcription, and inhibits tumor growth in prostate cancer models. These effects are abolished by a demethylation-resistant FOXA1 K270R mutant, confirming the specificity of this axis ^44^.

We therefore tested LSD1 inhibition in HER2^+^ DCIS. Treatment of the HER2^+^ DCIS cell line SUM225 with the LSD1 inhibitor ORY-1001 markedly reduced FOXA1 expression along with its downstream targets HER2 and CEACAM6 ^45,46^ (**Figure 6c,d**). FOXA1 was robustly expressed in both primary tumors and lung metastases of the HER2^+^ metastatic xenograft model BCM-3613 (data not shown). We then treated 5 mice with control and 7 mice with ORY-1001. ORY-1001 treatment significantly reduced primary tumor volume over time compared to control (**Figure 6e**). Although metastatic lesion size trended downward, this observation did not reach significance (**Figure 6f**); however, FOXA1 expression was significantly reduced in both lung metastases and primary tumors (**Figure 6g,h**). These results suggest that LSD1 inhibition attenuates metastatic burden, and suppresses FOXA1-driven transcriptional programs, with more complete control of metastatic disease likely requiring combination with cytotoxic or targeted agents. Together, our results establish LSD1 inhibition as a potential strategy to disrupt FOXA1-associated regulatory programs in high-risk DCIS.

## Discussion

In this study, integrated single-cell transcriptomics, chromatin accessibility, and spatial profiling across 43 patient-derived DCIS and DCIS–IDC samples reveal that cellular stemness is a key biological feature of DCIS lesions with invasive potential, with stem-like epithelial cells enriched in lesions that progress to invasion.

### Expansion of stem-like LumHR cells during DCIS progression

Both single-cell and spatial datasets revealed a consistent compositional shift across disease progression: Basal and luminal secretory cells declined from benign tissue through DCIS to IDC, while LumHR cells progressively expanded. LumHR cells also carried the highest stemness scores of any epithelial population, identifying them as both the dominant compartment in DCIS and the principal carriers of invasion-associated stem-like features. Our finding is in line with a recent manuscript which applied single-cell multiomic (WellDR-seq) method to ER+ invasive breast cancers^47^. This study showed that tumors arise from LumHR epithelial cells, which form ancestral subclones that acquire additional genomic alterations and expand into the dominant cancer population, establishing LumHR as the likely cell-of-origin. The study also finds that although sporadic aneuploid cells exist in other epithelial (*i.e.,* LumSec) and stromal compartments, only LumHR-derived clones undergo significant expansion. In relation to our findings, this study provides a mechanistic framework for the progressive enrichment of LumHR cells from benign tissue through DCIS to IDC, demonstrating that this shift is driven by the clonal selection and expansion of LumHR-derived tumor-initiating cells, consistent with their stem-like and invasion-associated properties. In another related study, Qin et al. used single-cell multi-omics to investigate cellular changes associated with DCIS progression to invasive breast cancer^48^. Consistent with our findings, they observed a significant reduction in basal/myoepithelial cells in both ER+ and ER− DCIS compared to normal tissue. LumHR cells were increased in both subtypes, whereas LumSec cells decreased in ER+ DCIS but were significantly increased in ER− DCIS. Notably, HER2 status was not assessed in this cohort. The differences between our findings and those of Qin et al. may reflect differences in cohort composition, as our study did not include triple-negative DCIS and instead focused primarily on HER2^+^ and ER+ cases. Together, these findings support a model in which breast cancer progression is driven by coordinated epithelial compositional shifts characterized by loss of basal and luminal secretory populations and expansion of LumHR cells. The convergence of our results with independent single-cell and multiomic studies suggests that LumHR cells represent the dominant, stem-like, tumor-initiating population, whose clonal expansion underlies disease progression from DCIS to invasive carcinoma.

### Stemness drives invasive behavior independent of proliferation and CNVs

Although experimental and clinical studies highlight dysregulation of stemness and cellular plasticity as key steps in metastatic progression, whether these processes underlie DCIS malignant transition remains unknown. Recent advances provide new opportunities to quantify stemness in cancer. Algorithms such as CytoTRACE predict cellular stemness from single-cell RNA sequencing (scRNA-seq) data^12^. The concept is that less differentiated or more stem-like cells tend to express a broader repertoire of genes (i.e., have a higher number of detectable expressed genes), so CytoTRACE uses the number of expressed genes per cell (“gene counts”) as a predictor of developmental potential^12^. CytoTRACE was validated across ∼150,000 single-cell transcriptomes encompassing 315 phenotypes, 52 lineages, 14 tissue types, 9 sequencing platforms, and 5 species, and outperforms other stemness algorithms^12^. Importantly, genome-wide chromatin accessibility analyses using scATAC-seq in embryonic and hematopoietic stem cells demonstrated a strong concordance between the number of accessible peaks, the number of expressed genes per cell, and CytoTRACE-predicted stemness scores^12^. These findings highlight the key role of the epigenome in regulating epithelial stemness and malignant transition of cancer cells. We hypothesized that CytoTRACE-based stemness score may discriminate DCIS with invasive potential. To test our hypothesis, we used the Mouse INtraDuctal (MIND) model by directly testing whether stemness distinguishes progressing from non-progressing DCIS. Across normal, focally amplified, and aneuploid epithelial cells (InferCNV), stemness scores were higher in progressors than non-progressors, with the largest gap in aneuploid cells. Critically, stemness was negatively correlated with overall CNV burden, indicating that aneuploidy alone is insufficient, and it is the combination of chromosomal instability with a stem-like state that drives progression. Stemness was positively correlated with chromatin organization, metabolism, proteasomal activity, MYC signaling, and EMT meta-programs, but only weakly or negatively with cell cycle and stress programs and significantly correlated with a published DCIS recurrence gene signature. Together, these findings establish stemness, rather than CNV burden, as a key driver of progression and recurrence risk. Consistent with our study, scRNA-seq was used to identify stemness-associated malignant epithelial cell populations that drive cancer progression in esophageal squamous cell carcinoma (ESCC) ^49^, lung adenocarcinoma (LUAD) ^50^, and colorectal cancer (CRC) ^51^. All three papers showed that tumors harbor subpopulations of cells with stem–like properties—characterized by high proliferative capacity, self-renewal, and dedifferentiation—which are linked to worse clinical outcomes. In ESCC, a distinct proliferative stem-like cluster (marked by genes such as MKI67 and UBE2C) was used to build a stemness-associated scoring model (SASM) that predicts survival and immune infiltration. Similarly, in LUAD, a transcriptome-based stemness index (mRNAsi) and machine learning models revealed that high-stemness tumors exhibit poor prognosis, increased malignancy, and immunosuppressive microenvironments. In CRC, a novel single-cell stemness signature (SCS_sig) demonstrated that stemness exists along a continuous spectrum rather than discrete states, reflecting high cellular plasticity and heterogeneity. Collectively, these studies, together with our own, underscore the strong translational potential of integrating single-cell transcriptomic approaches to characterize tumor stemness, enabling more precise prognostic stratification and the identification of clinically actionable therapeutic targets.

### Stem-like cells form spatial niches in high-risk DCIS

Beyond cell-intrinsic features, we identified discrete spatial niches enriched for stem-like LumHR cells within DCIS lesions, marked by structured interactions with immune and lymphatic cells including CEACAM6–EGFR signaling. While prior work has documented progression-associated changes in immune composition and stromal signaling without linking them to specific epithelial states ^10^, our findings connect these observations: stem-like epithelial cells occupy defined microenvironments that likely reinforce plasticity and invasion. Recent studies increasingly support the idea that cancer stem-like cells are not randomly distributed but instead reside within spatially organized microenvironmental niches shaped by specific cell–cell interactions, as demonstrated by spatial transcriptomics and single-cell analyses across multiple tumor types. For example, work in glioblastoma and other cancers has shown that distinct tumor cell states are maintained by microenvironmental cues ^52^, while spatial mapping approaches have revealed structured tumor–stroma and immune interactions within defined tissue regions ^53^. However, most prior work has examined either cell states or microenvironmental signaling independently, or inferred interactions without fully resolving their spatial coupling. Our findings extend this framework by directly linking stem-like epithelial states to discrete spatial niches and specific ligand–receptor interactions (*e.g.,* CEACAM6–EGFR), providing stronger evidence that microenvironmental context actively reinforces tumor cell plasticity and progression.

### FOXA1 as an epigenetic regulator of stemness in DCIS

Integrated scRNA/ATAC sequencing identified FOXA1 as a candidate regulator of DCIS stemness. FOXA1 expression and motif accessibility were highest in stemness-enriched epithelial clusters, and stemness scores correlated strongly with FOXA1 levels. FOXA1 is an established pioneer transcription factor that shapes enhancer landscapes and luminal identity in breast cancer ^54^; our data extend this role to maintenance of stem-like transcriptional programs underlying invasive potential.

### From lineage gatekeeper to plasticity driver: contextual control of FOXA1 function

How FOXA1 can simultaneously support lineage specification and stemness remains an important mechanistic question. Using synthetic ChIP-ISO, Xu et al. showed that FOXA1 occupancy is primarily governed by motif strength and co-factor cooperation, rather than chromatin context ^55^. FOXA1 binds high-affinity consensus motifs independently; however, its engagement at the more prevalent suboptimal motifs within active enhancers requires cooperation with co-factors, most notably AP-1 and CEBPB, through weak, polymorphic protein–protein interactions that are tolerant of variable spacing. Importantly, modulation of AP-1 levels redistributes FOXA1 binding genome-wide, indicating that co-factor availability, rather than chromatin environment alone, dictates FOXA1 occupancy landscapes. Functionally, this mechanism enables FOXA1 to access weak enhancers and promote transcriptional plasticity rather than fixed lineage commitment. In cancer, this context-dependent activity can be co-opte to sustain progenitor-like states, facilitate lineage reprogramming, and support adaptive responses to therapy.

Eyunni et al. ^56^ reinforced this contextual model in prostate cancer, showing that distinct FOXA1 mutation classes drive divergent trajectories. Class 1 mutations initiate androgen-dependent adenocarcinomas via coordinated AR and PI3K/mTORC1 activation, while C-terminal truncating Class 2 mutations promote progression by establishing an AP-1/KLF5-centered neo-cistrome that specifies a progenitor-like luminal state with increased TEAD engagement, paralleling regulatory programs in stem-like, castration-resistant disease. Together, these studies frame FOXA1 as a context-dependent regulator: strong motifs support stable, lineage-defining programs, while weak motifs require co-factor cooperation and enable the dynamic regulation underlying plasticity and stemness. Whether FOXA1 drives stemness in DCIS predominantly through high-affinity motifs or co-factor–dependent engagement at suboptimal sites remains to be determined.

### Therapeutic targeting of stemness through LSD1 inhibition

To test whether FOXA1-driven stemness is therapeutically targetable, we inhibited LSD1, which directly demethylates FOXA1 to enable its pioneer function ^44^. LSD1 inhibition reduced FOXA1, HER2, and CEACAM6 expression in vitro and significantly suppressed tumor growth in vivo. LSD1 may also suppress FOXA1-dependent transcription by altering H3K4 methylation at enhancers, consistent with FOXA1’s preferential recruitment to distal enhancers marked by H3K4me2 enrichment and H3K9me2 exclusion ^54^. Additional contributions are likely, as LSD1 also promotes EMT through cooperation with SNAIL family transcription factors ^57^; these mechanisms warrant further investigation.

LSD1 inhibitors are already in clinical development, primarily in hematologic malignancies. Early trials of iadademstat (ORY-1001) demonstrate manageable toxicity and biological activity ^58,59^. These data support the feasibility of repurposing LSD1 inhibitors for early interception in high-risk DCIS.

### Limitations

This study has several limitations. First, although the cohort is large for a multimodal single-cell analysis of DCIS, the number of samples within individual molecular subtypes remains limited. Larger cohorts with long-term clinical follow-up will be required to validate stemness as a predictive biomarker. Second, invasive potential was assessed using the MIND model rather than patient outcomes; while highly informative, this model cannot fully capture systemic influences present in humans. Third, our functional validation focused on HER2^+^ disease, and whether FOXA1-LSD1–driven stemness is equally relevant in luminal DCIS remains to be determined.

## Methods

### Tissue acquisition

Ductal carcinoma in situ (DCIS) and/or DCIS–invasive ductal carcinoma (DCIS/IDC) tissues were obtained from patients undergoing surgical resection at the University of Kansas Medical Center under protocols approved by the institutional Human Subjects Committee (HSC; IRB# 11513), with written informed consent obtained from all participants.

### Tissue processing for single-cell analysis

Upon receipt, each tissue was weighed and then transferred to a Teflon block, finely minced with scalpels, and transferred to a 50-ml conical tube containing freshly prepared, filter-sterilized digestion medium (10 ml per 1 g of tissue contained 5 mg collagenase [Roche Applied Science, Indianapolis, IN], 0.24 mg hyaluronidase [2140 units/mg; Sigma-Aldrich St. Louis, MO], 200 mg BSA, 100 μl antibiotic-antimycotic, and 10 ml DMEM/F12). Following a 16-h incubation (50 rpm at 37°C), the specimens were removed, briefly shaken by hand, and centrifuged at 200 × g for 1 min. Prewarmed trypsin-EDTA (1 ml; Stem Cell Technologies, Vancouver, BC) was added to the resulting pellet and gently pipetted up and down for 1 min. Hank’s balanced salt solution (HBSS) with 2% fetal bovine serum (FBS) (HF)was added, and specimens were centrifuged at 400 × g for 5 min. The supernatant was removed, and 1 ml prewarmed 5 mg/ml Dispase (Stem Cell Technologies) and 100 μl of 1 mg/ml DNase I (Stem Cell Technologies) were added to the pellet. To resuspend the pellet, the specimen was pipetted up and down for 1 min. An additional 10 ml cold HF was added to the cell suspension, which was filtered through a 40-μm cell strainer. The cell suspension was centrifuged at 400 × g for 5 min, the resulting cell pellet was resuspended in 200 μl phosphate-buffered saline (PBS), and cells were counted. Cells were then frozen in 93% FBS and 7% DMSO and stored in liquid nitrogen until intraductal injection and/or single cell analysis.

### Animals and MIND surgeries

Recipient mice were 8- to 10-week-old virgin female NOD-SCID IL2Rgammanull (NSG) mice, which were purchased from Jackson Laboratories. Animal experiments were conducted following protocols approved by the University of Kansas School of Medicine Animal Care and Use and Human Subjects Committee.

For MIND surgeries, a 30-gauge Hamilton syringe with a 50-μl capacity and a blunt-ended 1/2-inch needle was used to deliver the cells as previously described ^13^. Two microliters of PBS (with 0.04% trypan blue) containing ∼35,000 cells were injected. After 6-12 months, mice were sacrificed, and mammary tissues were either digested and processed for magnetic sorting and/or fixed and processed for embedding.

### Immunofluorescence (IF)

IF was performed as previously described ^60^. Nuclei were counterstained with hoechst. Negative controls were carried out using secondary antibodies without primary antibodies. Imaging was performed on a laser-scanning confocal microscope (Model 510; Carl Zeiss MicroImaging, Inc, Thornwood, NY, USA). The acquisition software used was Pascal (Carl Zeiss MicroImaging, Inc). Procedures for the evaluation of MIND model invasiveness has been published ^14^.

### Mouse mammary gland processing and magnetic sorting

To obtain “Post” transplant cells, mammary glands from MIND xenografts were excised (at a median of 9 months post-intraductal injection) and digested overnight as described above. Single mammary epithelial cells were then magnetically labeled with mouse MHCI/II antibodies (listed in **List of Antibodies**) and with MACS Anti-Biotin MicroBeads UltraPure (Miltenyl Biotec #130-105-637) and negatively sorted for human DCIS cells using Miltenyl LD columns (Miltenyl Biotec #130-042-901) following the manufacturer’s protocol. A sample of sorted cells was then analyzed with fluorescence activated cell sorting (FACS) to examine the purity, and the rest of the cells were cryopreserved.

### List of Antibodies

|  | Company | Catalog # | Dilution |
| --- | --- | --- | --- |
| **IF staining** |  |  |  |
| Primary antibodies |  |  |  |
| Cytokeratin 5 | Vector | VPC400 | 1:25 |
| Cytokeratin 19 | Thermoscientific | MS198 | 1:50 |
| SMA | Thermoscientific | PA5-18292 | 1:50 |
| Secondary antibodies |  |  |  |
| Alexa Fluor 488 anti-rabbit | Invitrogen | A11008 | 1:200 |
| Alexa Fluor 594 anti-mouse | Invitrogen | A21203 | 1:200 |
| Alexa Fluor 488 anti-mouse | Invitrogen | A11012 | 1:200 |
| **Magnetic sorting** |  |  |  |
| Mouse MHC I | BD Pharmingen | 553546 | 1:100 |
| Mouse MHC II | BD Pharmingen | 553564 | 1:100 |
| **Flow cytometry** |  |  |  |
| Human EpCAM | Abcam | ab37305 | 1:20 |

### Single cell multiome ATAC + gene expression sample preparation and sequencing

Single cells were obtained from the surgical specimens following an overnight digestion using Fresh or cryopreserved specimens as described. Viable nuclei were isolated from single cell suspensions using optimized lysis conditions. Nuclei were counted and assessed for integrity prior to downstream processing.

Single-cell multiome ATAC and gene expression libraries were prepared using a commercially available microfluidic platform (*e.g.,* Chromium iX, 10x Genomics) following the manufacturer’s instructions. Briefly, isolated nuclei were loaded to enable partitioning into nanoliter-scale droplets, where simultaneous transposition of accessible chromatin and reverse transcription of mRNA were performed with unique cellular barcodes. Following barcoding, ATAC fragments and cDNA were recovered and amplified separately to generate sequencing-ready libraries.

Libraries were quantified using fluorometric assays and assessed for fragment size distribution by capillary electrophoresis. Indexed libraries were pooled and sequenced on an Illumina platform (Novaseq 6000) using paired-end reads to a depth consistent with recommended guidelines for both gene expression and chromatin accessibility assays.

### Tissue preparation for Xenium Samples

Ductal carcinoma in situ (DCIS) and/or DCIS–invasive ductal carcinoma (DCIS/IDC) tissues were obtained from patients undergoing surgical resection at the University of Kansas Medical Center under protocols approved by the institutional Human Subjects Committee (HSC; IRB# 11513), with written informed consent obtained from all participants. Formalin-fixed paraffin-embedded (FFPE) tissue sections (5 µm) were mounted on Xenium-compatible glass slides following the manufacturer’s protocol (10x Genomics, Xenium In Situ User Guide). Slides were baked, deparaffinized (for FFPE), and subjected to antigen retrieval to optimize probe accessibility. Tissue morphology was preserved through fixation and mild permeabilization steps.

### Gene panel design

A 380 targeted gene panel was used to detect gene expression. This panel was made up of the 280 Xenium Human Breast Panel and 100 add-on genes (**Supplemental Table 2b**).

### Probe hybridization and in situ chemistry

Transcript detection was performed using the Xenium Human Breast Panel with 100 add-on genes (10x Genomics, Supplementary Table 2b), which employs a multiplexed probe-based hybridization strategy for direct mRNA detection within intact tissue architecture. After deparaffinization and permeabilization, a custom probe panel along with control probes was added to the slides at a final concentration of 10 nM, followed by overnight hybridization at 50 °C. Stringent post-hybridization washes were then performed to remove unbound probes. Subsequently, probe ligation and annealing of the rolling circle amplification primer were carried out at 37 °C for 2 hours. The circularized probes were enzymatically amplified for 2 hours at 37 °C, generating multiple copies of the gene-specific barcodes corresponding to each RNA-binding event. After amplification, samples were washed and chemically quenched to reduce background fluorescence. Finally, nuclei were stained with DAPI, and the sections were loaded onto the Xenium Analyzer for imaging and transcript detection. Sections were incubated with reagents and fluorescently labelled probes for RNA detection, which were automatically cycled in, imaged, and removed by the Xenium Analyzer over 15 imaging rounds. Z-stack images were acquired across the full tissue thickness at 0.75-μm step intervals. Sequential rounds of fluorescent oligonucleotide ligation, imaging, and signal cleavage enabled subcellular-resolution transcript quantification across thousands of RNA targets simultaneously.

### Imaging and data acquisition

Automated high-resolution imaging was conducted using the Xenium Analyzer. Fluorescence images were captured across multiple spectral channels to detect unique probe barcodes. Raw image data were processed using Xenium Onboard Analysis (XOA) software, which performs image alignment, spot calling, and base calling to generate decoded transcript maps. Z-stack images acquired by the Xenium Analyzer in each cycle and fluorescence channel were processed and stitched using the DAPI channel as a reference to generate a single composite image representing one region of interest (ROI). This image was used to construct a spatial map of transcript distribution across the tissue section. Every punctum, representing an observed photon event, was detected by the Xenium Onboard Analysis (XOA) pipeline across all imaging cycles and channels to capture potential mRNA signals. Emitted light from localized puncta at the same spatial coordinates across cycles was registered and fitted to a Gaussian distribution. Using the fluorescence intensity values from all four color channels across 15 Xenium cycles, an optical barcode unique to each gene was generated. A Phred-scaled quality score (Q score) was then assigned to each decoded transcript, and transcripts with Q ≥ 20 were retained for downstream spatial gene plots.

Cell segmentation was performed using DAPI images, defining boundaries within a 15-μm radius from each nucleus or until contact with adjacent cells. The on-instrument pipeline generated output files including the feature-cell matrix, decoded transcripts, and a CSV file of cell boundaries (containing differentially enriched genes per cluster). All subsequent data integration and downstream analyses were performed off-instrument and visualized using Xenium Explorer.

### Post-Xenium histology

Following Xenium image acquisition, hematoxylin and eosin (H&E) staining was performed on the same tissue microarray sections previously subjected to Xenium analysis. Bright-field images of the sections were captured before and after H&E staining at 20× magnification using a Keyence microscope (Itasca, IL). The resulting images were stitched to generate high-resolution composite images of the tissue sections. Xenium Explorer was then used to align the post-Xenium bright-field images with the corresponding Xenium fluorescence images.

### Single cell RNA (scRNA) data processing, integration, and clustering

Raw sequencing reads were aligned to GRCh38 (human) reference using Cell Ranger (v.7.1.0) software with default parameters. Downstream analysis was performed on filtered features counts generated by Cell Ranger. Low-quality cells were removed using the following thresholds: number of genes per ≤ 300 and ≥ 10000, log_10_(Genes/UMI) < 0.8, or percent mitochondrial reads per cell > 10%. A gene was removed if it was not detected in at least 10 cells. Samples were integrated using scVI in the scvi-tools Python package with the following parameters: n_latent=10, n_layer=1, gene_likelihood=’zinb’. Integrated data were then clustered based on the scVI representation using the Leiden algorithm and a resolution of 1.0.

### scRNA Cell Type Labeling

To annotate cell types, we downloaded scRNA-seq data reported in Kumar et al., 2023 ^61^ using the CELLxGENE database. Then, the Seurat *TransferData* function was used to transfer cell type labels from their scRNA-seq to our scRNA-seq data.

### CytoTRACE

To predict the differentiation state of each cell, we used CytoTRACE ^12^, which outputs a stemness score ranging from 0 to 1. To have comparable stemness scores across all samples, we used the *iCytoTRACE* function in R. For the scRNA-seq data, CytoTRACE was first run using all cells in the dataset, and then only for epithelial (Basal, LumSec, LumHR) cells to compute the stemness score. This algorithm was also used for the Xenium and snATAC/RNA-seq data, independently.

### Copy number variation (CNV) analysis

Copy number variation (CNV) analysis was performed using inferCNV ^62^ on single-cell RNA-sequencing data to infer large-scale chromosomal alterations from gene expression profiles. Briefly, genes were ordered according to chromosomal position, and expression intensities in epithelial/tumor cells were normalized relative to a reference population of non-malignant cells. Smoothed expression patterns across genomic intervals were then used to infer gains and losses across chromosomes. Cells were classified based on the magnitude and pattern of inferred CNV signal: cells showing broad chromosomal gains and/or losses were annotated as **aneuploid**, cells showing discrete high-level regional amplifications were classified as **focally amplified**, and cells lacking substantial large-scale CNV deviations relative to the reference population were classified as **normal**.

### Module scores of meta-programs and 812-gene signature

Genes defining the 41 meta-programs were downloaded from Gavish et al ^23^ and the 812 genes defining the DCIS recurrence classifier were downloaded from Strand et al ^11^. The *AddModuleScore* function in the Seurat package was used to calculate an expression score for each of the meta-programs and the 812 gene classifier. The stemness scores of LumHR cells were then correlated to the computed module scores. Pearson correlation was used for correlation between stemness and meta-program, Spearman correlation was used for correlation between stemness and the 812 gene signature.

### Xenium data processing, integration, and clustering

Raw data and standard output files from the XOA pipeline were used for downstream analysis. Cells were initially segmented using the DAPI morphology image and assigning transcripts to the closest nucleus within a maximum distance of 15 µm. But, based on current recommendations and newest version of the Xenium suite of tools from 10x Genomics, we re-segmented cells with Xenium-Ranger (v2.0.1) using a nuclear expansion distance of 5µm. Only cells with 10 or more counts were kept for downstream analysis. Integration of all 16 Xenium data sets was performed across using scVI in the scvi-tools Python package with the following parameters: n_latent=10, n_layer=1, gene_likelihood=’zinb’. Integrated data were then clustered based on the scVI representation using the Leiden algorithm and a resolution of 1.0.

### Labeling of major cell types in Xenium data

To label cell types in the Xenium data we used a two-fold approach 1) transferring labels from a reference scRNA and snRNA data set and 2) looking at cluster specific expression and comparing to known cell type markers. The results from both these methods were then compared to create a consensus cell type labeling for all clusters identified in the Xenium data. More specifically, for the first method, we downloaded scRNA-seq and snRNA-seq data reported in Kumar et al., 2023^61^ using the CELLxGENE database. Then, we used Seurat to transfer cell type labels from the scRNA-seq and the snRNA-seq to the Xenium data. For the second method, we looked at the reported cell type markers of the scRNA-seq and snRNA-seq data that were measured in the Xenium data and corrected any clusters that might have been mislabeled based on the label transfer alone. This two-fold approach was used since the Xenium panel only consisted of 380 genes, which can affect the accuracy of the transfer data method.

### Sub-clustering of epithelial, immune, and stromal cell populations

Epithelial cells (Basal, LumSec, LumHR), immune cells (Myeloid-1, Myeloid-2, Mast, Tcell, Plasma, Bcell), and stromal cells (Adipocyte, Vascular, Lymphatic, Perivascular, CAFs, Fibroblast) were separately extracted from the full dataset and underwent their own integration and clustering to identify more specific subtype specific cell populations. Integration and clustering were performed in the same way as before. Marker genes of each epithelial, immune, or stromal cell population were found using the *FindAllMarkers* function in Seurat with the Wilcoxon rank sum test. A gene was defined as a marker gene of the population if the adjusted p-value < 0.05 and proportion of expressing cells > 0.5. Visualizations of epithelial cell populations on the Xenium image were created in Xenium Explorer v.3.

### Enrichment analysis of epithelial cell populations

To test functional enrichment of marker genes of the epithelial cell populations we used overrepresentation analysis. The input gene sets were the marker genes for each population as defined above. We tested for enrichment in the Human Molecular Signatures Database hallmark gene sets and previously described gene-expression programs in cancer cells which are termed meta-programs (MP) ^23^. The enricher function in R was used and a p-value < 0.05 was considered a statistically significant overlap.

### Xenium differential expression of DCIS and IDC regions

Using the pathologist annotated H&E images, we extracted cells from the annotated regions and labeled them as benign, DCIS, or IDC accordingly. The cell extraction of those regions were done in Xenium Explorer using the free-hand selection tool. Then, differential expression analyses were conducted on LumHR labeled cells in these regions. Testing was performed using the *FindMarkers* function in Seurat with the Wilcoxon rank sum test. For IDC versus DCIS comparisons, the analysis was performed within sample individually. A gene was differentially expressed if the adjusted p-value < 0.05 and proportion of expressing cells > 0.1. Results were then compared across samples within subtype to find genes that are commonly differentially expressed between IDC and DCIS. For IDC versus IDC comparisons, the test was done grouping HER2^+^ samples together and Luminal samples together. A gene was differentially expressed if the adjusted p-value < 0.05 and proportion of expressing cells > 0.1 in all samples of that subtype.

### Colocalization analysis of epithelial cell populations

Colocalization analysis was performed to understand cell-cell interactions of the epithelial sub-clusters in the spatial data. First, Delaunay triangulation simplices were constructed using the centroid x- and y-coordinates of each epithelial cell using the geometry::*delaunayn* function in R. Then, the simplices were converted into an edge list by extracting all edges of connecting cell types. Each edge was annotated with the epithelial subtype label of the two connecting cells, and the number of interacting cell types was quantified by counting the number of edges between each pair of cell types. Next, the edge list was converted into a matrix, with the rows and columns corresponding to epithelial cell types, and the entries in the matrix representing the total number of edges observed between the corresponding pair of cell types. The matrix was normalized by dividing each value by the row sum, and the matrix was made symmetric by taking the max value. This normalized value for a pair of cell types represents the colocalization score, or the relative frequency of an interaction. This process was done independently for each sample. To visualize how the epithelial sub-clusters interact with each other in each sample, we used this colocalization score to create a graph, using the igraph package in R. The nodes represent the cell types, with their size reflective of the abundance of the cell type in the sample. Edge thickness is reflective of the colocalization score. A threshold of 0.1 was used for edge filtering.

### Spatial nearest-neighbor analysis

To calculate the distances of immune and stromal cell populations from benign epithelium, DCIS lesions, and IDC regions, a nearest-neighbor analysis was used. Reference populations were defined as cells classified as either 1) benign epithelium, 2) DCIS lesions, or 3) IDC. Target populations were defined as all other stromal and immune cell populations. Then, for each reference population cell, the distance to the nearest target population cell was calculated for all target populations (*e.g.,* DCIS cell to nearest Fibroblast cell, DCIS cell to nearest T cell, etc.). Nearest neighbor search was conducted using the *nn2* function in the R package RANN. Xenium spatial coordinates are reported in µm so the Euclidean distances reported between spatial coordinates can be directly interpreted as distances in µm.

### Spatial cross-type pair correlation function analysis

The cross-type pair correlation function (*g*(*r*)) measures the association between two types of points (*a* and *b*) at a specified distance (*r*). For each sample independently, two-dimensional multitype point pattern objects were constructed using the cell centroid coordinates using the *ppp* function in the spatstat packaged in R. Points were marked by their cell type with epithelial cells as either benign epithelium, DCIS, or IDC, and all other immune and stromal cell types as their specific cell type label. The *pcfcross* function from spatstat was used to compute *g*(*r*) using a range of distances (*r*) ranging from 0 to 500 µm by increments of 5 µm. Edge correction was implemented with the “translate” method. We evaluated the function for all stromal and immune subsets (*b*) relative to the different areas of IDC, DCIS, or benign epithelium (*a*). For example, to evaluate the spatial clustering of Fibroblasts and DCIS cells at distances of 100 µm, *g_a,b_*(*r*) = *g_DCIS,Fibroblast_*(100). A *g*(*r*) = 1 indicates independence of the two cell types, meaning the cell types are randomly distributed; *g*(*r*) > 1 indicates there is more clustering of the two cell types than expected by random chance; *g*(*r*) < 1 indicates the two cell types are more dispersed from each other than expected by random chance. We also summarized the spatial correlation per group (HER2^+^ DCIS, HER2^+^ DCIS/IDC, Luminal DCIS, and Luminal DCIS/IDC) by computing a weighted average of *g*(*r*) at each distance evaluated for each set of samples. The weights were the number of pairs evaluated (i.e., the number of IDC, DCIS, or benign cells multiplied by the number of target cells).

### Spatially constrained clustering to identify niches

Spatially constrained clustering was performed using a method similar to the *scc* function in the Python dynamo package, which integrates both gene expression profiles and spatial locations for the identification of tissue domains. For each sample, we constructed a 𝑘 nearest-neighbor graph (𝐺*_expression_*) based on gene expression profiles, with 𝑘 = 15. Next, a graph (𝐺*_spatial_*) was created from the spatial coordinates using Delaunay triangulation. Scanpy’s *neighbors* function and Squidpy’s *spatial_neighbors* function were used to create each graph, respectively. Then, we constructed a joint graph (𝐺*_joint_*.) by adding the two graphs together, weighted by a parameter α:

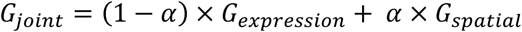

We used an α of 0.25 to give more weight to the gene expression graph. Then clustering was performed on 𝐺*_joint_*. using the Leiden algorithm. We assessed clustering (i.e., niches) at resolutions from 0.2 to 1.0 by increments of 0.1.

### Ligand-receptor-mediated cell-cell communication analysis

Ligand–receptor–mediated cell–cell communication was analyzed using the SpaCCI (Spatially Aware Cell–Cell Interaction analysis) package in R ^63^. A ligand-receptor (LR) database was put together by pulling LR pairs from CellChat, CellPhoneDB, and OmniPath. After pulling LR pairs with available gene expression in the Xenium data, there were only 100 LR pairs we could evaluate with the data. Xenium samples were partitioned into different areas to evaluate the differences in interactions between niches with differing stemness scores. For each partitioned area, for each sample, SpaCCI was run. Significant interactions were determined by an adjusted p-value < 0.05.

### Single cell multiome ATAC + gene expression (scATAC/RNA) data processing

Raw sequencing reads were aligned to GrCh38 reference using Cell Ranger ARC software with the default parameters. Downstream analysis on the gene expression data (scRNA) was performed on filtered features counts generated by the pipeline. Low-quality cells were removed using the following thresholds: number of genes per ≤ 250, log_10_(Genes/UMI) < 0.8, or percent mitochondrial reads per cell > 15%. scRNA data were used to compute CytoTRACE scores and annotate cell types. Cell types were annotated in the same as the other scRNAseq data as described above. Single-cell ATAC (scATAC) data were processed using ArchR ^64^ (v1.0.3). Arrow files were generated with the *createArrowFiles* function using a minimum transcription start site (TSS) enrichment score of 5 and a minimum of 1000 fragments per cell. Tile matrices and gene score matrices were generated during Arrow file construction. Doublet scores were computed using *addDoubletScores*, and an ArchR project was created from the resulting Arrow files. Cells identified as putative doublets were subsequently filtered out using *filterDoublets*. To match cells across modalities, ArchR cell barcodes were reformatted and aligned to the corresponding scRNA-seq barcodes, and only cells present in both modalities were retained. Cell type labels and stemness scores derived from the matched scRNA-seq analysis were added to the ArchR project using *addCellColData*. The projected stemness score was further rescaled to the unit interval for visualization and comparison. Dimensionality reduction was performed using iterative latent semantic indexing (LSI) based on the tile matrix, followed by graph-based clustering with *addClusters*. UMAP embeddings were generated from the LSI representation and used to visualize cluster structure, transferred cell type labels, and projected stemness scores.

### Transcription factor motif enrichment analysis

For peak calling and motif analysis, pseudo-bulk group coverages were generated by cluster using *addGroupCoverages*, and reproducible peak sets were identified using MACS2 through the *addReproduciblePeakSet* function. A peak accessibility matrix was then constructed using *addPeakMatrix*, and transcription factor binding motif annotations were added using HOMER motif sets via *addMotifAnnotations*. Within each sample subtype, we performed differential accessibility and motif enrichment analyses by comparing the cluster with the highest stemness score (Cluster 1) against all remaining clusters. Specifically, differential accessibility analysis was carried out using *getMarkerFeatures* with Wilcoxon rank-sum testing and bias correction for TSS enrichment and log-transformed fragment counts. Motif enrichment analysis was subsequently performed using *peakAnnoEnrichment*, restricting attention to peaks with FDR <= 0.05 and Log2FC >= 0.5.

### Statistical analysis

Differences in stemness scores and differences in proportions of different cell types were tested with a Wilcoxon rank sum test. A p-value < 0.05 was considered significantly different. In cases of multiple testing, p-values were corrected with the Benjamini-Hochberg procedure and adjusted p-values (q-value) < 0.05 was considered significantly different.

Change in tumor volume over time between controls and treatment with ORY-1001 was tested with a linear mixed-effects model. The model was fitted with random intercepts for mouse and fixed effects for time, treatment group, and their interaction on tumor volume. We reported the p-value for the interaction between time and treatment group, which would indicate whether the change in tumor volume over time differed among control and ORY-1001 treated mice. Differences in average size of metastasis and FOXA1 histoscores were tested with a Wilcoxon rank sum test. A p-value < 0.05 was considered significantly different.

## Acknowledgements

This work was supported by Cancer Research UK and by KWF Kankerbestrijding (ref. C38317/A24043; FB); KU Cancer Center Support Grant (P30 CA168524) & The Kansas Institute for Precision Medicine – COBRE (P20 GM130423) (AKG); Pilot Grant: NCATS Frontiers-CTSA grant from NCATS awarded to the University of Kansas for Frontiers, ref# UL1TR002366 (FB), Dr. Ralph and Marian Falk Medical Research Trust (FB), BC230628P1 (FB), R01CA271588 (FB); TBEL CDMC grant U24CA274212 (ZL); STRIDE (MDACC) funding to METI and NIH RO1 1R01CA226269 (KR).

We would like to acknowledge support from the University of Kansas Medical Center’s Biospecimen Repository Core Facility staff, including Maura Kluthe, Alex Webster, Eric Johnson, and Lauren DiMartino for helping to identify, collect, and process the human specimens. This study was supported in part by the KU Cancer Center’s Support Grant (P30 CA168524) and the Kansas Institute for Precision Medicine (P20 GM130423)

## Author Contributions

Conceptualization and scientific direction: FB, KR, ES, ZL, SAK, AT. Data Work and Analysis: ES, VK, EA, YD, PY, ZL, RH, JL, MM, RM, TF, NN, DCK. Technical and Resource Contributions: ES, VK, EA, YD, PY, ZL, RH, JL, MM, RM, TF, JLW, KL, CB, AA, AH, OW, JP, MH, CS, EJJ, LJK, CF, MTL, AKG, DCK. Writing and Communication: All Authors. Oversight and Support: FB, KR, AT, ZL, MM, RM, TF, SAK, CF, AKG.

## References

1 Leonard, G. D. & Swain, S. M. Ductal carcinoma in situ, complexities and challenges. J Natl Cancer Inst 96, 906–920 (2004).

2 Collins, L. C. et al. Outcome of patients with ductal carcinoma in situ untreated after diagnostic biopsy: results from the Nurses’ Health Study. Cancer 103, 1778–1784, doi:10.1002/cncr.20979 (2005).

3 Page, D. L., Dupont, W. D., Rogers, L. W. & Landenberger, M. Intraductal carcinoma of the breast: follow-up after biopsy only. Cancer 49, 751–758 (1982).

4 Sanders, M. E., Schuyler, P. A., Dupont, W. D. & Page, D. L. The natural history of low-grade ductal carcinoma in situ of the breast in women treated by biopsy only revealed over 30 years of long-term follow-up. Cancer 103, 2481–2484, doi:10.1002/cncr.21069 (2005).

5 Maxwell, A. J. et al. Risk factors for the development of invasive cancer in unresected ductal carcinoma in situ. Eur J Surg Oncol 44, 429–435, doi:10.1016/j.ejso.2017.12.007 (2018).

6 Maxwell, A. J. et al. Unresected screen-detected ductal carcinoma in situ: Outcomes of 311 women in the Forget-Me-Not 2 study. Breast 61, 145–155, doi:10.1016/j.breast.2022.01.001 (2022).

7 Early Breast Cancer Trialists’ Collaborative, G. et al. Overview of the randomized trials of radiotherapy in ductal carcinoma in situ of the breast. J Natl Cancer Inst Monogr 2010, 162–177, doi:10.1093/jncimonographs/lgq039 (2010).

8 Narod, S. A., Iqbal, J., Giannakeas, V., Sopik, V. & Sun, P. Breast Cancer Mortality After a Diagnosis of Ductal Carcinoma In Situ. JAMA Oncol 1, 888–896, doi:10.1001/jamaoncol.2015.2510 (2015).

9 Narod, S. A. & Sopik, V. Is invasion a necessary step for metastases in breast cancer? Breast Cancer Res Treat 169, 9–23, doi:10.1007/s10549-017-4644-3 (2018).

10 Risom, T. et al. Transition to invasive breast cancer is associated with progressive changes in the structure and composition of tumor stroma. Cell 185, 299–310 e218, doi:10.1016/j.cell.2021.12.023 (2022).

11 Strand, S. H. et al. Molecular classification and biomarkers of clinical outcome in breast ductal carcinoma in situ: Analysis of TBCRC 038 and RAHBT cohorts. Cancer Cell 40, 1521–1536 e1527, doi:10.1016/j.ccell.2022.10.021 (2022).

12 Gulati, G. S. et al. Single-cell transcriptional diversity is a hallmark of developmental potential. Science 367, 405–411, doi:10.1126/science.aax0249 (2020).

13 Behbod, F. et al. An intraductal human-in-mouse transplantation model mimics the subtypes of ductal carcinoma in situ. Breast cancer research : BCR 11, R66, doi:10.1186/bcr2358 (2009).

14 Hong, Y. et al. Mouse-INtraDuctal (MIND): an in vivo model for studying the underlying mechanisms of DCIS malignancy. J Pathol 256, 186–201, doi:10.1002/path.5820 (2022).

15 Sflomos, G. et al. A Preclinical Model for ERalpha-Positive Breast Cancer Points to the Epithelial Microenvironment as Determinant of Luminal Phenotype and Hormone Response. Cancer Cell 29, 407–422, doi:10.1016/j.ccell.2016.02.002 (2016).

16 Molinaro, A. M., Sison, J. D., Ljung, B. M., Tlsty, T. D. & Kerlikowske, K. Risk prediction for local versus regional/metastatic tumors after initial ductal carcinoma in situ diagnosis treated by lumpectomy. Breast Cancer Res Treat 157, 351–361, doi:10.1007/s10549-016-3814-z (2016).

17 Hutten, S. J. et al. A living biobank of patient-derived ductal carcinoma in situ mouse-intraductal xenografts identifies risk factors for invasive progression. Cancer Cell 41, 986–1002 e1009, doi:10.1016/j.ccell.2023.04.002 (2023).

18 Kumar, T. et al. A spatially resolved single-cell genomic atlas of the adult human breast. Nature 620, 181–191, doi:10.1038/s41586-023-06252-9 (2023).

19 Soloff, M. S. Oxytocin receptors and mammary myoepithelial cells. J Dairy Sci 65, 326–337, doi:10.3168/jds.S0022-0302(82)82194-2 (1982).

20 Jeong, J. Y., Jeoung, N. H., Park, K. G. & Lee, I. K. Transcriptional regulation of pyruvate dehydrogenase kinase. Diabetes Metab J 36, 328–335, doi:10.4093/dmj.2012.36.5.328 (2012).

21 Shou, Y. et al. TIMP1 Indicates Poor Prognosis of Renal Cell Carcinoma and Accelerates Tumorigenesis via EMT Signaling Pathway. Front Genet 13, 648134, doi:10.3389/fgene.2022.648134 (2022).

22 Morgenstern-Kaplan, D. et al. Genomic, immunologic, and prognostic associations of TROP2 (TACSTD2) expression in solid tumors. Oncologist 29, e1480–e1491, doi:10.1093/oncolo/oyae168 (2024).

23 Gavish, A. et al. Hallmarks of transcriptional intratumour heterogeneity across a thousand tumours. Nature 618, 598–606, doi:10.1038/s41586-023-06130-4 (2023).

24 Arnold, S. A. & Brekken, R. A. SPARC: a matricellular regulator of tumorigenesis. J Cell Commun Signal 3, 255–273, doi:10.1007/s12079-009-0072-4 (2009).

25 Cheng, G. et al. Higher levels of TIMP-1 expression are associated with a poor prognosis in triple-negative breast cancer. Mol Cancer 15, 30, doi:10.1186/s12943-016-0515-5 (2016).

26 Zhu, C. et al. ITGB3/CD61: a hub modulator and target in the tumor microenvironment. Am J Transl Res 11, 7195–7208 (2019).

27 Lim, E. et al. Aberrant luminal progenitors as the candidate target population for basal tumor development in BRCA1 mutation carriers. Nature medicine 15, 907–913, doi:10.1038/nm.2000 (2009).

28 Liu, R. et al. Krupple-like factor 5 is essential for mammary gland development and tumorigenesis. J Pathol 246, 497–507, doi:10.1002/path.5153 (2018).

29 Xu, Q. et al. Anterior Gradient 3 Promotes Breast Cancer Development and Chemotherapy Response. Cancer Res Treat 52, 218–245, doi:10.4143/crt.2019.217 (2020).

30 Taifour, T. et al. The tumor-derived cytokine Chi3l1 induces neutrophil extracellular traps that promote T cell exclusion in triple-negative breast cancer. Immunity 56, 2755–2772 e2758, doi:10.1016/j.immuni.2023.11.002 (2023).

31 Ross, D. & Siegel, D. Functions of NQO1 in Cellular Protection and CoQ(10) Metabolism and its Potential Role as a Redox Sensitive Molecular Switch. Front Physiol 8, 595, doi:10.3389/fphys.2017.00595 (2017).

32 Puram, S. V. et al. Single-Cell Transcriptomic Analysis of Primary and Metastatic Tumor Ecosystems in Head and Neck Cancer. Cell 171, 1611–1624 e1624, doi:10.1016/j.cell.2017.10.044 (2017).

33 Carvalho, E., Canberk, S., Schmitt, F. & Vale, N. Molecular Subtypes and Mechanisms of Breast Cancer: Precision Medicine Approaches for Targeted Therapies. Cancers 17, doi:10.3390/cancers17071102 (2025).

34 Bertucci, F., Finetti, P., Goncalves, A. & Birnbaum, D. The therapeutic response of ER+/HER2- breast cancers differs according to the molecular Basal or Luminal subtype. NPJ Breast Cancer 6, 8, doi:10.1038/s41523-020-0151-5 (2020).

35 Jolly, M. et al. Implications of the hybrid epithelial/mesenchymal phenotype in metastasis. Frontiers in Oncology 5, doi:10.3389/fonc.2015.00155 (2015).

36 Garg, M. Epithelial, mesenchymal and hybrid epithelial/mesenchymal phenotypes and their clinical relevance in cancer metastasis. Expert Rev Mol Med 19, e3, doi:10.1017/erm.2017.6 (2017).

37 Yu, M. et al. Circulating breast tumor cells exhibit dynamic changes in epithelial and mesenchymal composition. Science 339, 580–584, doi:10.1126/science.1228522339/6119/580 [pii] (2013).

38 Tirosh, I. et al. Single-cell RNA-seq supports a developmental hierarchy in human oligodendroglioma. Nature 539, 309–313, doi:10.1038/nature20123 (2016).

39 Glabman, R. A., Choyke, P. L. & Sato, N. Cancer-Associated Fibroblasts: Tumorigenicity and Targeting for Cancer Therapy. Cancers 14, doi:10.3390/cancers14163906 (2022).

40 Bahri, M., Anstee, J. E., Opzoomer, J. W. & Arnold, J. N. Perivascular tumor-associated macrophages and their role in cancer progression. Essays Biochem 67, 919–928, doi:10.1042/EBC20220242 (2023).

41 Yang, S., Du, J., Wang, W., Zhou, D. & Xi, X. APOC1 is a prognostic biomarker associated with M2 macrophages in ovarian cancer. BMC Cancer 24, 364, doi:10.1186/s12885-024-12105-z (2024).

42 Chiang, W. F. et al. Carcinoembryonic antigen-related cell adhesion molecule 6 (CEACAM6) promotes EGF receptor signaling of oral squamous cell carcinoma metastasis via the complex N-glycosylation. Oncogene 37, 116–127, doi:10.1038/onc.2017.303 (2018).

43 Pinkert, J. et al. T cell-mediated elimination of cancer cells by blocking CEACAM6-CEACAM1 interaction. Oncoimmunology 11, 2008110, doi:10.1080/2162402X.2021.2008110 (2022).

44 Gao, S. et al. Chromatin binding of FOXA1 is promoted by LSD1-mediated demethylation in prostate cancer. Nat Genet 52, 1011–1017, doi:10.1038/s41588-020-0681-7 (2020).

45 Zhao, M. et al. FOXA1, induced by RC48, regulates HER2 transcription to enhance the tumorigenic capacity of lung cancer through PI3K/AKT pathway. J Cancer 15, 5863–5875, doi:10.7150/jca.100210 (2024).

46 Wang, W. et al. CircDIAPH1 Promotes Liver Metastasis and Development of Colorectal Cancer by Initiation of CEACAM6 Expression. Mol Carcinog 64, 897–910, doi:10.1002/mc.23896 (2025).

47 Wang, K. et al. Coalescing single-cell genomes and transcriptomes to decode breast cancer progression. Cell 188, 6355–6369 e6316, doi:10.1016/j.cell.2025.08.012 (2025).

48 Qin, X. et al. Single-Cell Expression Analysis of Ductal Carcinoma In Situ Identifies Complex Genotypic-Phenotypic Relationships Altering Epithelial Composition. Cancer Res 85, 2302–2319, doi:10.1158/0008-5472.CAN-24-3023 (2025).

49 Ye, W., Su, W., Lei, C., Huang, C. & Du, M. Single-cell analysis identifies a stemness-associated tumor cell subpopulation and develops a prognostic scoring model in esophageal squamous cell carcinoma. Translational oncology 64, 102653, doi:10.1016/j.tranon.2025.102653 (2026).

50 Lin, G. et al. scRNA-seq revealed high stemness epithelial malignant cell clusters and prognostic models of lung adenocarcinoma. Scientific reports 14, 3709, doi:10.1038/s41598-024-54135-4 (2024).

51 Lin, K. et al. Identification of Colorectal Cancer Cell Stemness from Single-Cell RNA Sequencing. Mol Cancer Res 22, 337–346, doi:10.1158/1541-7786.MCR-23-0468 (2024).

52 Neftel, C. et al. An Integrative Model of Cellular States, Plasticity, and Genetics for Glioblastoma. Cell 178, 835–849 e821, doi:10.1016/j.cell.2019.06.024 (2019).

53 Moncada, R. et al. Author Correction: Integrating microarray-based spatial transcriptomics and single-cell RNA-seq reveals tissue architecture in pancreatic ductal adenocarcinomas. Nat Biotechnol 38, 1476, doi:10.1038/s41587-020-00776-5 (2020).

54 Lupien, M. et al. FoxA1 translates epigenetic signatures into enhancer-driven lineage-specific transcription. Cell 132, 958–970, doi:10.1016/j.cell.2008.01.018 (2008).

55 Xu, C. et al. Systematic dissection of sequence features affecting binding specificity of a pioneer factor reveals binding synergy between FOXA1 and AP-1. Mol Cell 84, 2838–2855 e2810, doi:10.1016/j.molcel.2024.06.022 (2024).

56 Eyunni, S. et al. Divergent FOXA1 mutations drive prostate tumorigenesis and therapy-resistant cellular plasticity. Science 389, eadv2367, doi:10.1126/science.adv2367 (2025).

57 Ambrosio, S., Sacca, C. D. & Majello, B. Epigenetic regulation of epithelial to mesenchymal transition by the Lysine-specific demethylase LSD1/KDM1A. Biochim Biophys Acta Gene Regul Mech 1860, 905–910, doi:10.1016/j.bbagrm.2017.07.001 (2017).

58 Salamero, O. et al. Iadademstat in combination with azacitidine in patients with newly diagnosed acute myeloid leukaemia (ALICE): an open-label, phase 2a dose-finding study. Lancet Haematol 11, e487–e498, doi:10.1016/S2352-3026(24)00132-7 (2024).

59 Salamero, O. et al. First-in-Human Phase I Study of Iadademstat (ORY-1001): A First-in-Class Lysine-Specific Histone Demethylase 1A Inhibitor, in Relapsed or Refractory Acute Myeloid Leukemia. J Clin Oncol 38, 4260–4273, doi:10.1200/JCO.19.03250 (2020).

60 Valdez, K. E. et al. Human primary ductal carcinoma in situ (DCIS) subtype-specific pathology is preserved in a mouse intraductal (MIND) xenograft model. J Pathol 225, 565–573, doi:10.1002/path.2969 (2011).

61 Kumar, T. et al. A spatially resolved single cell genomic atlas of the adult human breast. bioRxiv, doi:10.1101/2023.04.22.537946 (2023).

62 Timothy Tickle, e. a. InferCNV of the Trinity CTAT Project<{https://github.com/broadinstitute/inferCNV}> (2019).

63 Bernard, V. et al. A spatial atlas of chemoradiation therapy in pancreatic cancer identifies cellular and microenvironmental determinants of persister populations. bioRxiv, doi:10.1101/2025.06.20.660757 (2025).

64 Granja, J. M., et al. ArchR: An integrative and scalable software package for single-cell chromatin accessibility analysis. BioRxiv (2020).

