## Supplementary Table Legends for "Cellular stemness identifies high-risk ductal carcinoma *in situ* and offers a therapeutic interception opportunity"

**Supplementary Table 1 | Clinicopathologic features and multi-omic profiling of DCIS samples.**

Summary of all DCIS samples analyzed in this study, including sequencing modality (scRNA-seq, scATAC/RNA-seq, or spatial transcriptomics) and timing relative to intraductal (MIND) transplantation (Pre or Post). Progression status indicates whether lesions progressed to invasive carcinoma in vivo or remained non-invasive. Molecular subtype was assigned based on ER, PR, and HER2 status. ER and PR positivity were defined as ≥1% nuclear staining by immunohistochemistry. HER2 status was determined by immunohistochemistry (0–3+) and confirmed by FISH where indicated; HER2 positivity was defined as IHC 3+ or FISH amplification. Tumor and nuclear grade, Ki67 proliferation index, p53 status, architectural patterns (comedo, cribriform, papillary, micropapillary, and solid), and presence of metastasis are indicated where available. Check marks denote presence of a feature; dashes denote absence; blank entries indicate data not available.

**Supplementary Table 2 | Single-cell and spatial transcriptomic annotation and differential gene expression analyses.**

**a,** Overview of the 16 Xenium samples, including molecular subtype (Luminal or $HER2^{+}$), disease stage (DCIS or DCIS/IDC), and metastasis status.

**b,** List of the 380 genes included in the Xenium gene panel.

**c,** Marker genes used for validation of Human Breast Cell Atlas (HBCA)–based cell-type annotations transferred from Kumar *et al.*

**d,** Marker genes defining epithelial subclusters identified by unbiased subclustering of epithelial cells.

**e,** Association of epithelial subclusters with molecular subtype (Luminal versus $HER2^{+}$) and pathologist-annotated regions (benign, DCIS or IDC).

**f,** Differentially expressed genes (DGEs) identified in luminal hormone-responsive (LumHR) subclusters comparing invasive ductal carcinoma (IDC) and ductal carcinoma in situ (DCIS) regions within Luminal and $HER2^{+}$ samples.

**g,** DGEs comparing IDC regions between $HER2^{+}$ and Luminal samples, highlighting subtype-specific transcriptional programs associated with invasive progression.
