## Supplementary Figures and Legends for "Cellular stemness identifies high-risk ductal carcinoma *in situ* and offers a therapeutic interception opportunity"

Supplementary Figure 1

### Xenium Sample 1: HER2+ DCIS (Metastasis to Lymph Nodes and Bone)

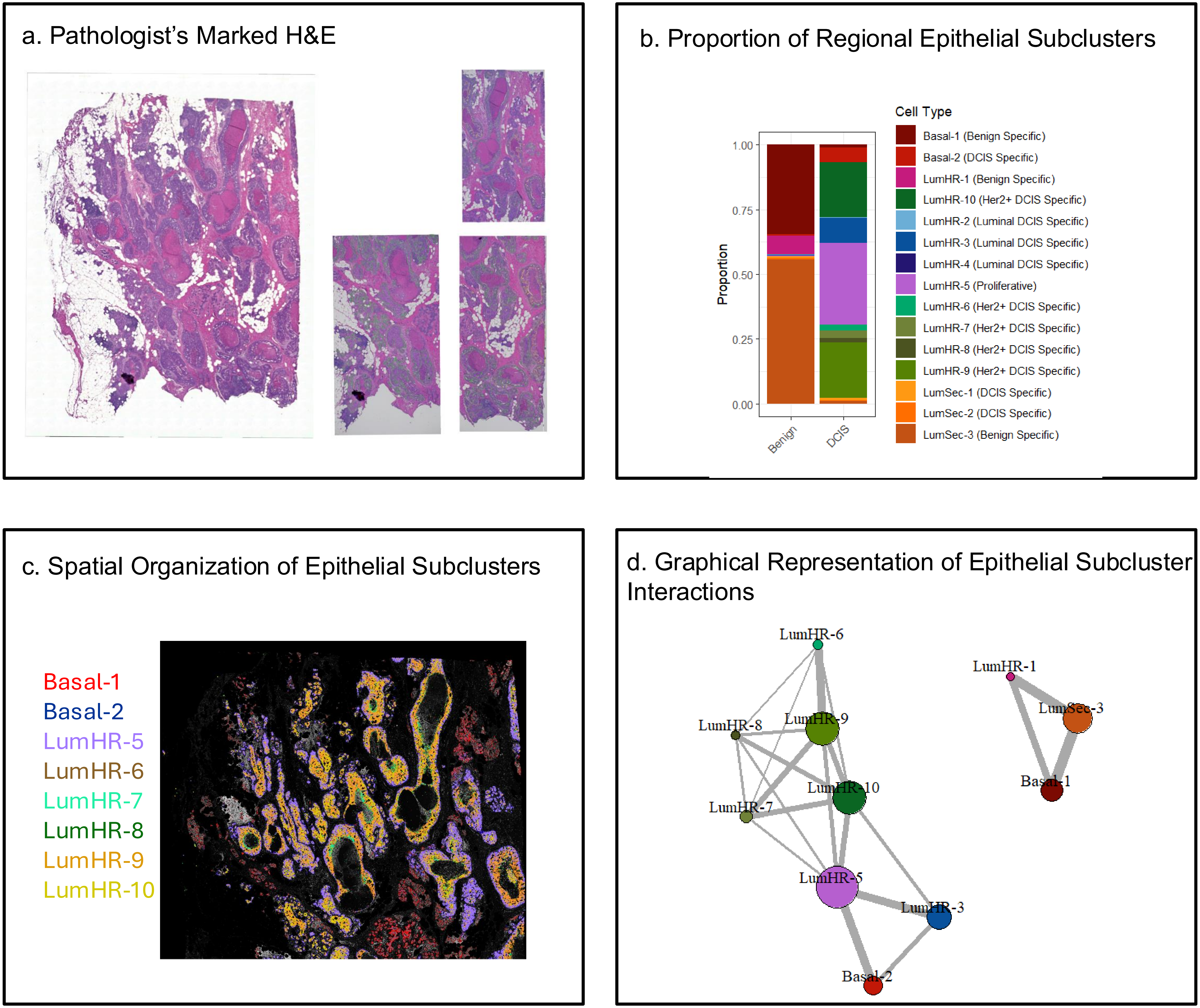

Supplementary Figure 1

### Xenium Sample 2: HER2+ DCIS

a. Pathologist's Marked H&E

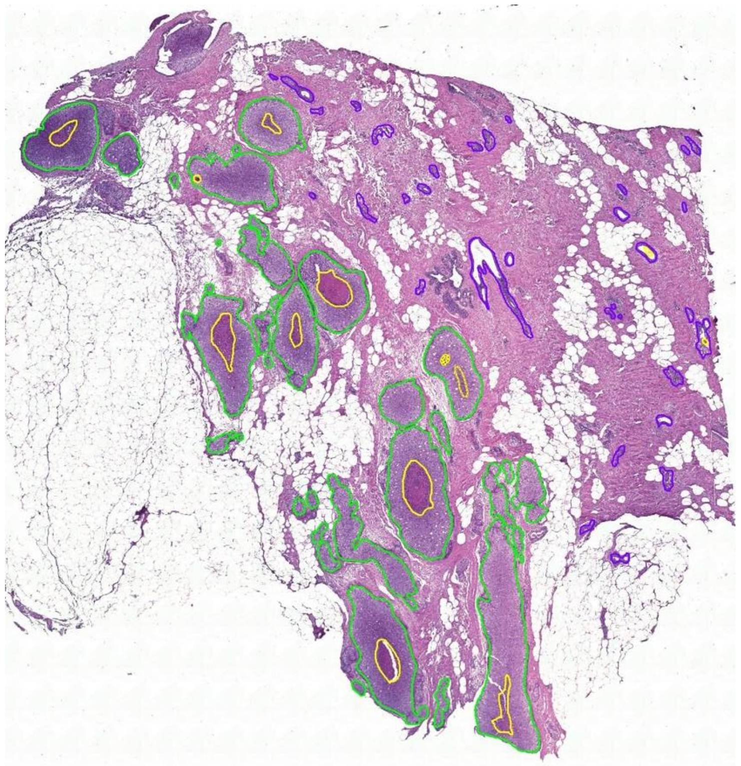

b. Proportion of Regional Epithelial Subclusters

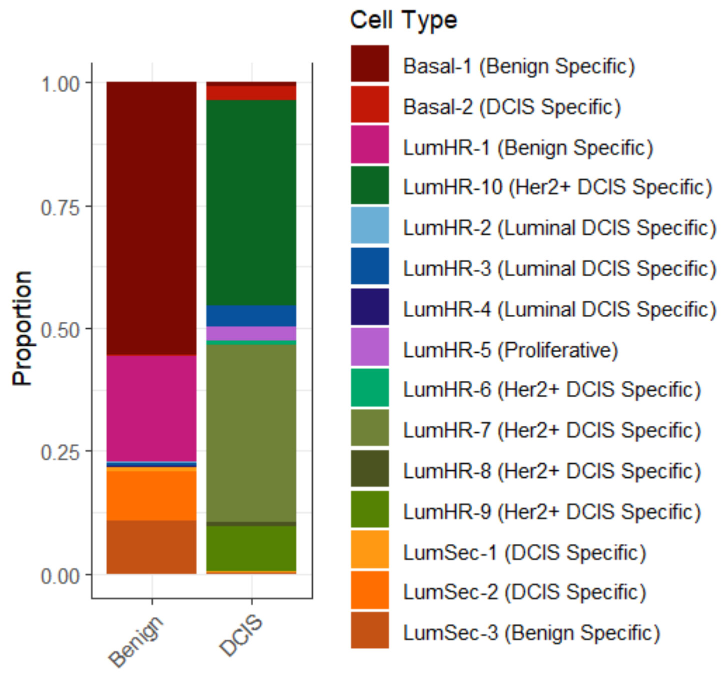


c. Spatial Organization of Epithelial Subclusters

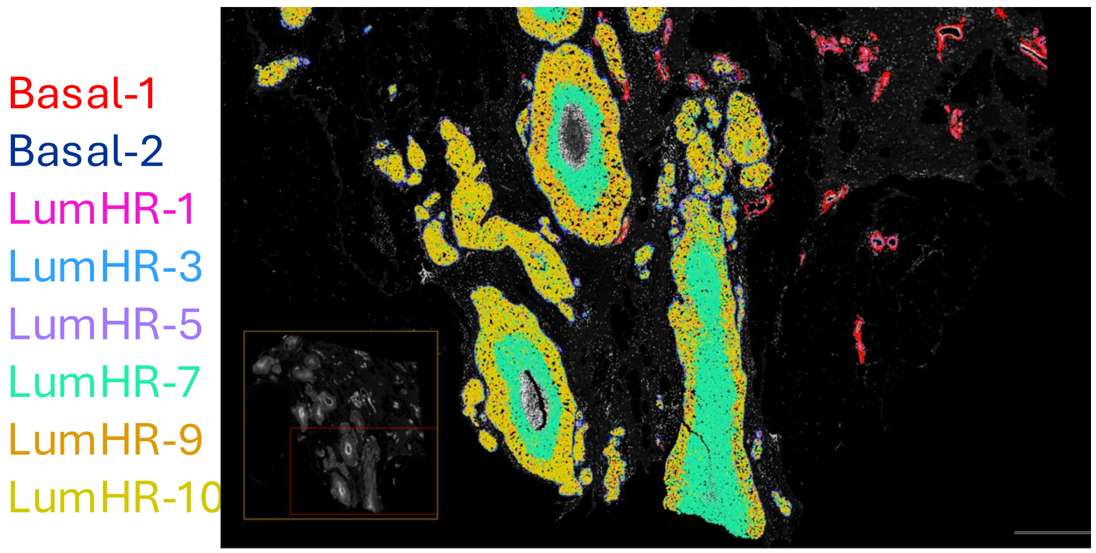


d. Graphical Representation of Epithelial Subcluster Interactions

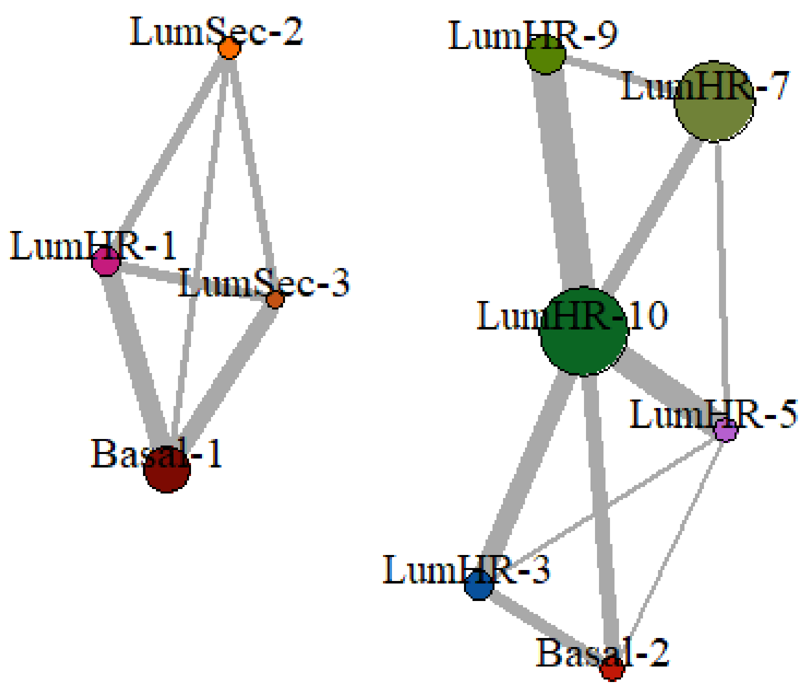

Supplementary Figure 1

### Xenium Sample 3: DCIS HER2+

#### a. Pathologist's Marked H&E

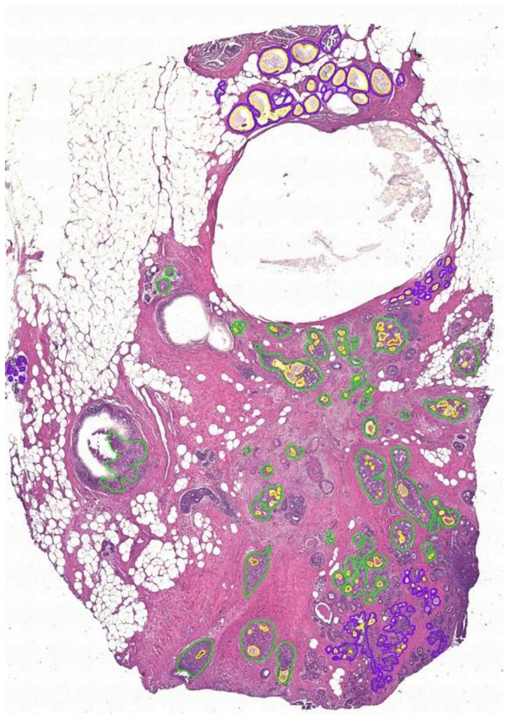

#### b. Proportion of Regional Epithelial Subclusters

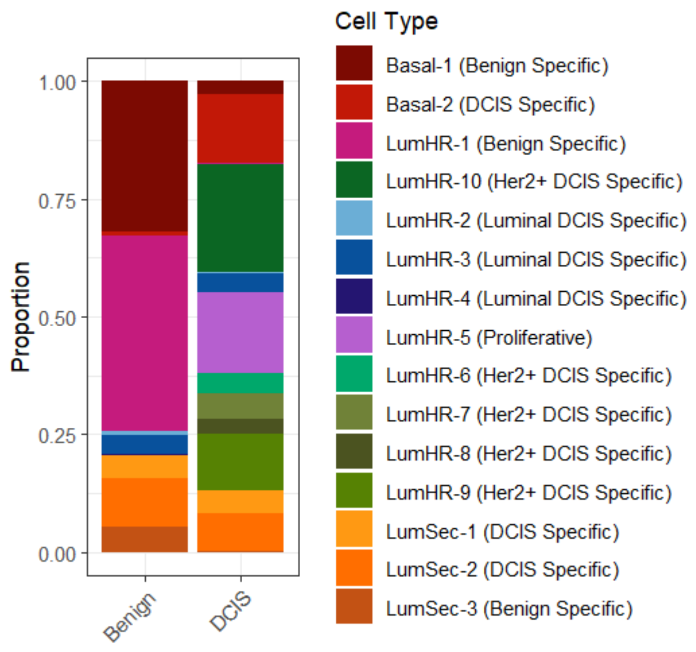


#### c. Spatial Organization of Epithelial Subclusters

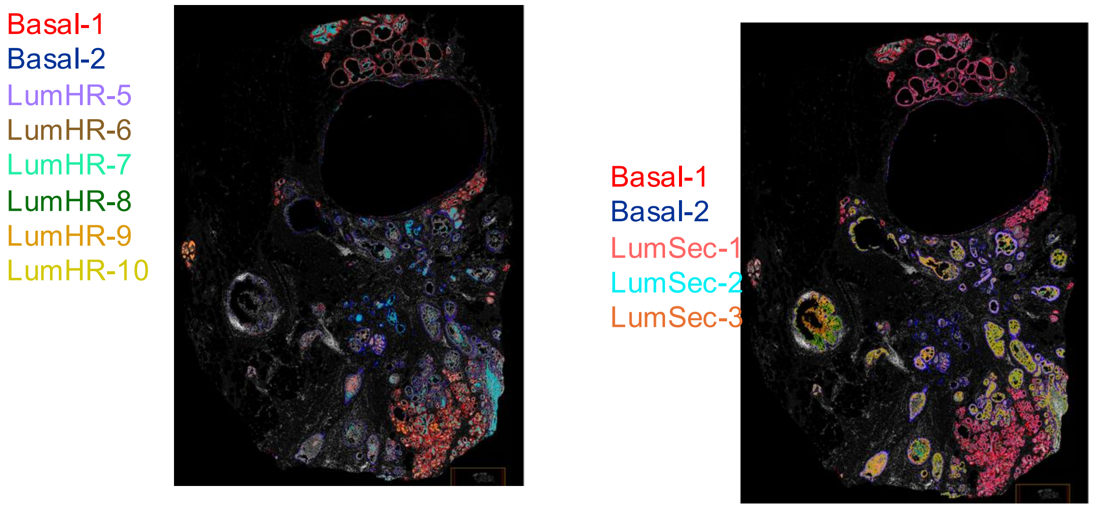


#### d. Graphical Representation of Epithelial Subcluster Interactions

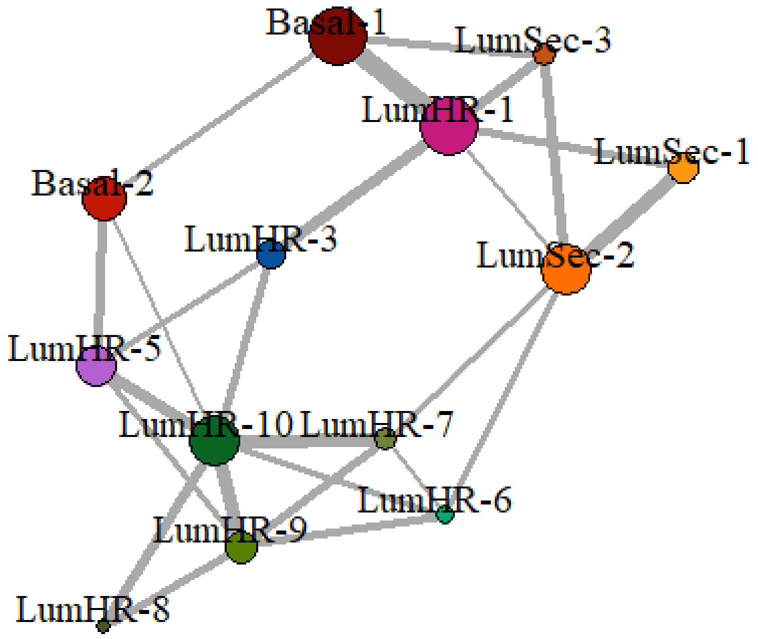

Supplementary Figure 1

### Xenium Sample 4: DCIS HER2+

a. Pathologist's Marked H&E

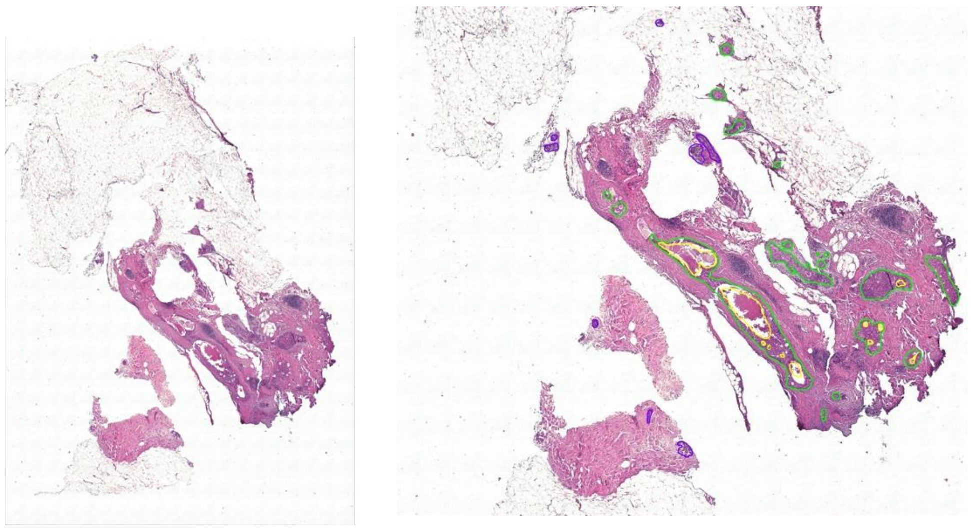

b. Proportion of Regional Epithelial Subclusters

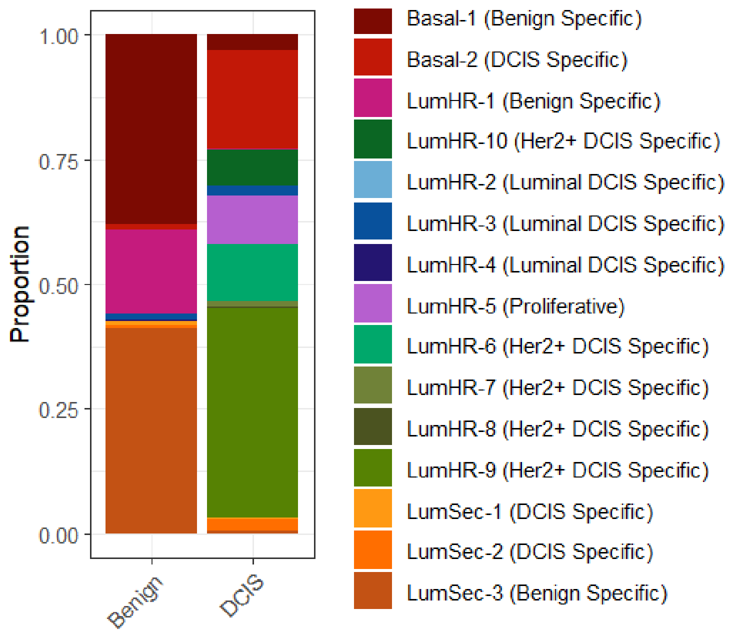


c. Spatial Organization of Epithelial Subclusters

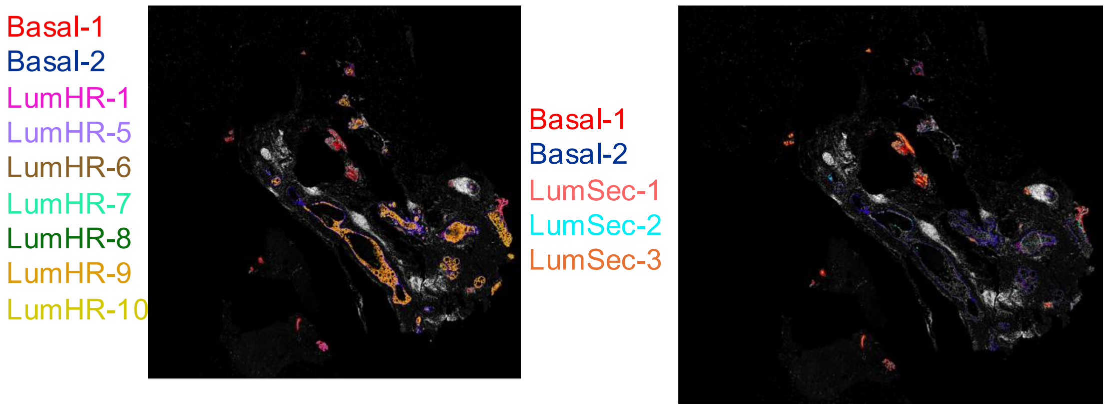


d. Graphical Representation of Epithelial Subcluster Interactions

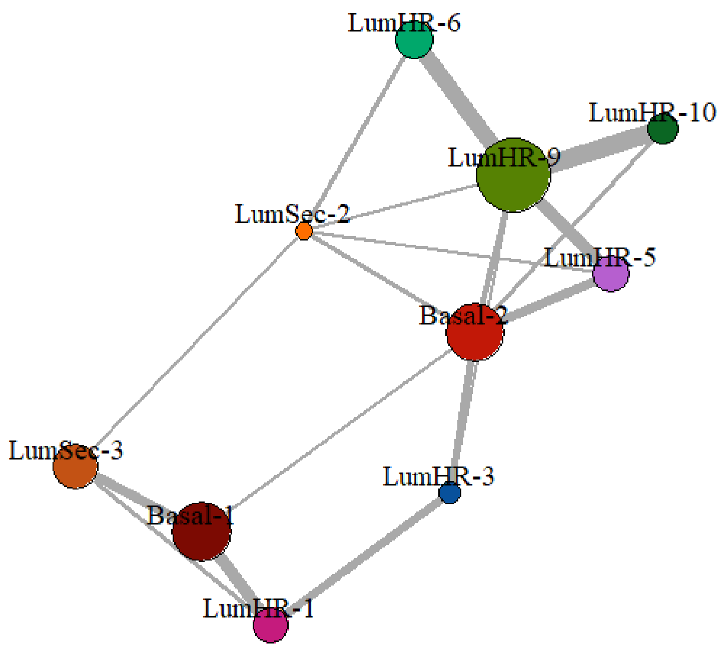

Supplementary Figure 1

### Xenium Sample 5: DCIS HER2+

a. Pathologist's Marked H&E

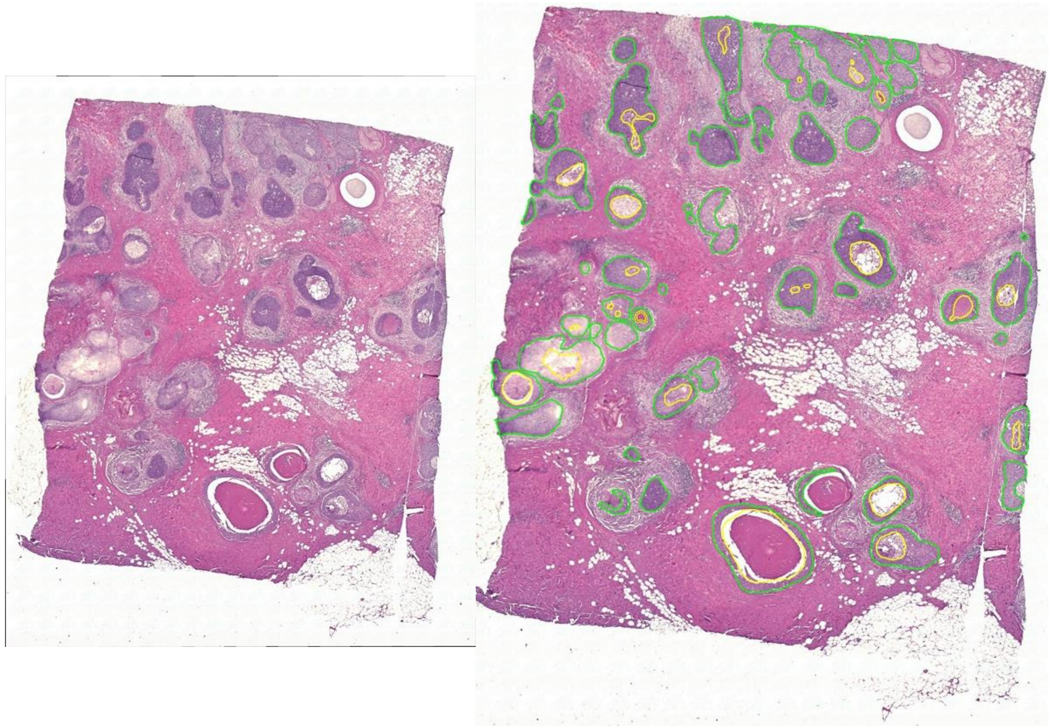

b. Proportion of Regional Epithelial Subclusters

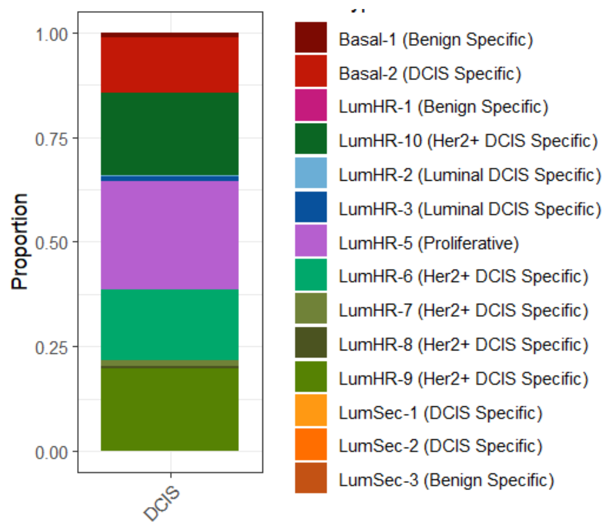


c. Spatial Organization of Epithelial Subclusters

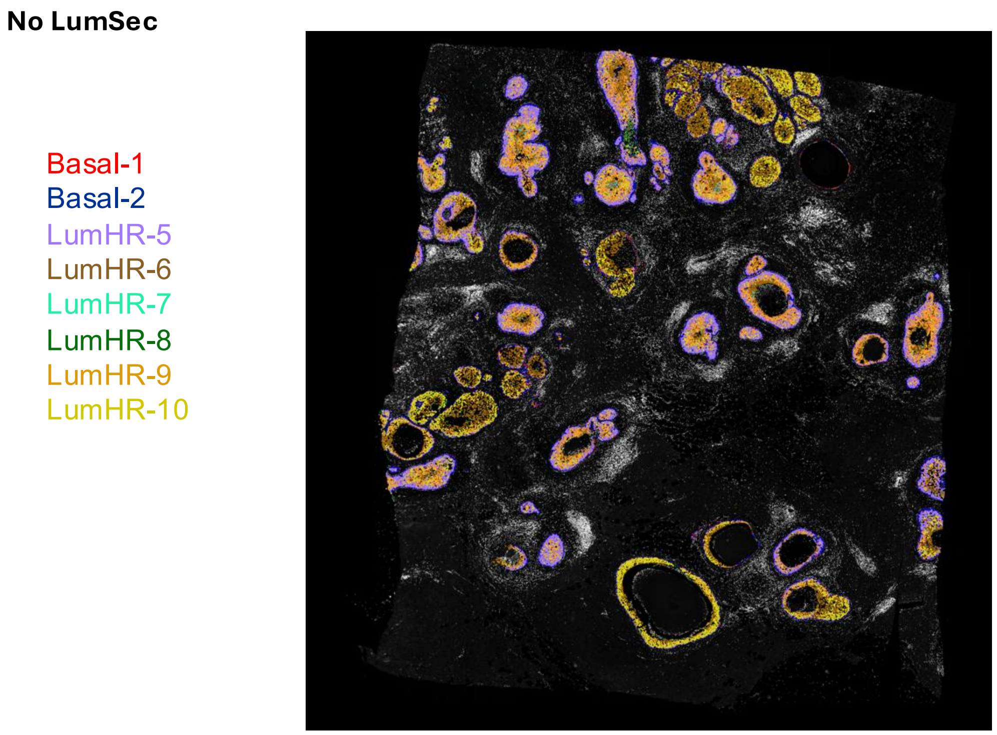


d. Graphical Representation of Epithelial Subcluster Interactions

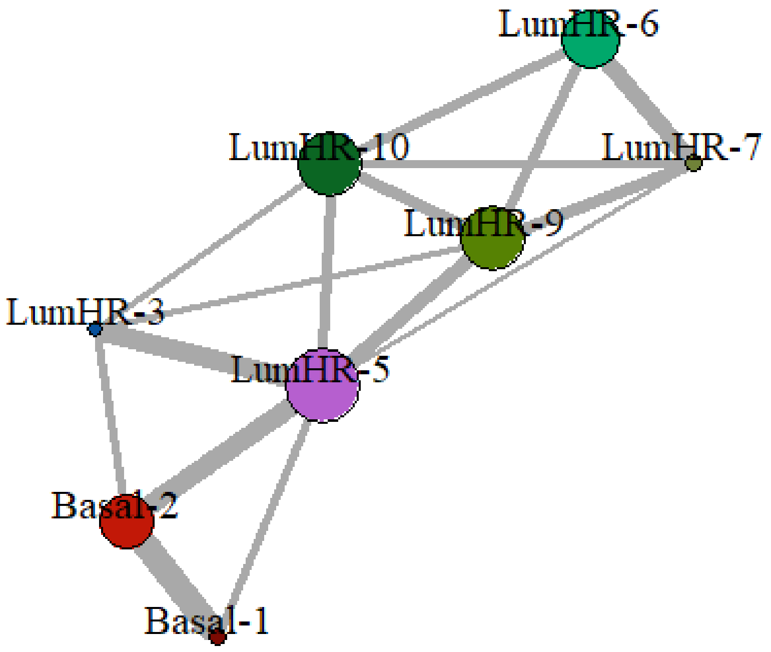

Supplementary Figure 1

### Xenium Sample 6: DCIS HER2+

a. Pathologist’s Marked H&E

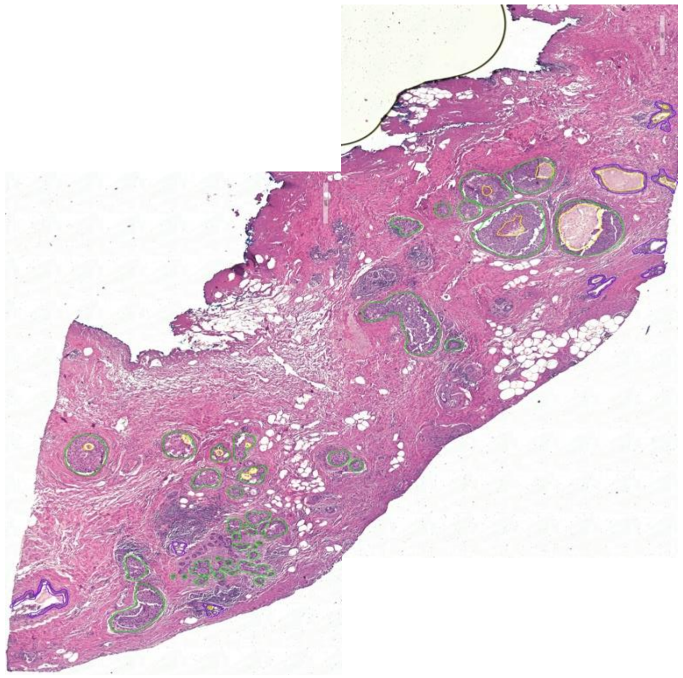

b. Proportion of Regional Epithelial Subclusters

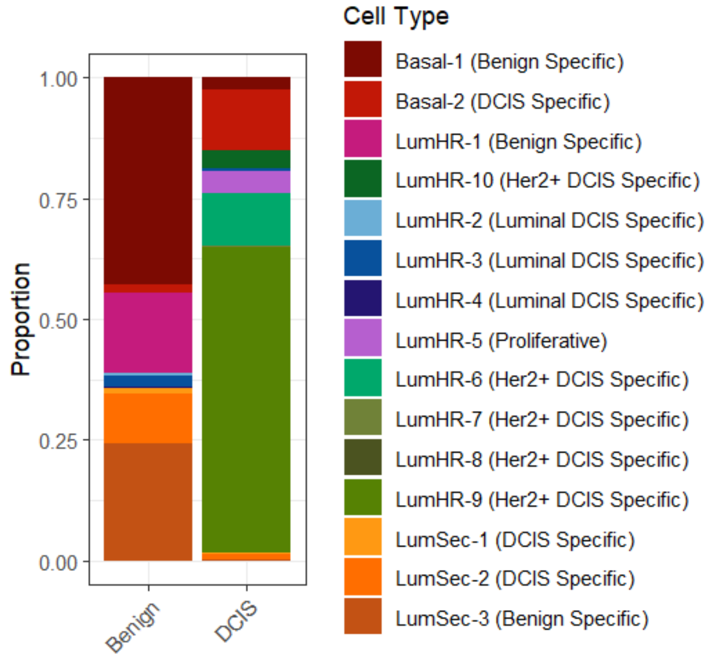


c. Spatial Organization of Epithelial Subclusters

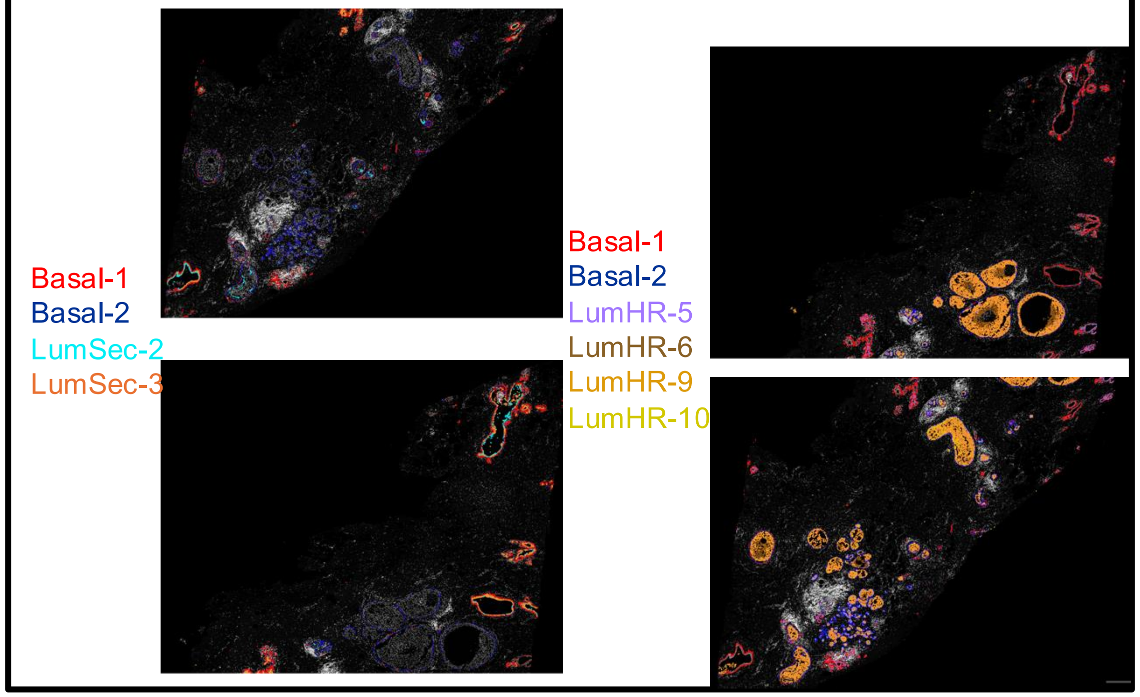


d. Graphical Representation of Epithelial Subcluster Interactions

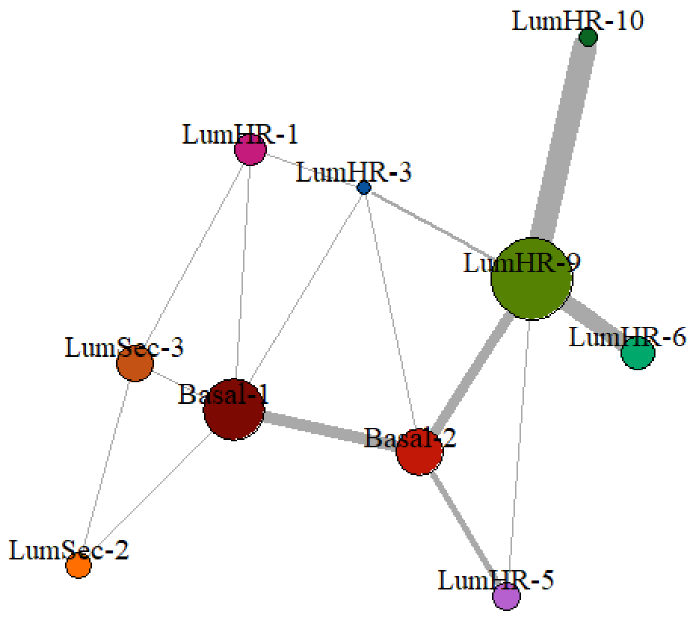

Supplementary Figure 1

### Xenium Sample 7: DCIS Luminal

a. Pathologist's Marked H&E

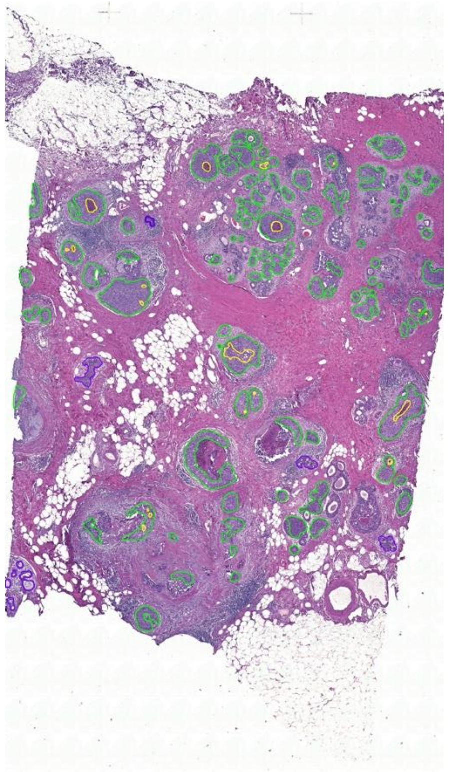

b. Proportion of Regional Epithelial Subclusters

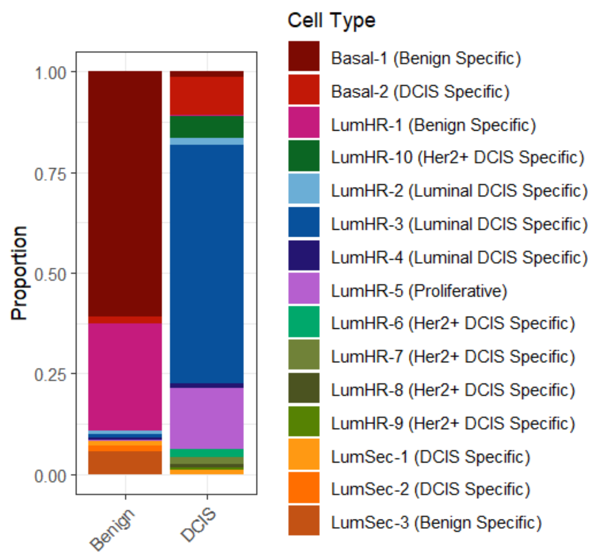


c. Spatial Organization of Epithelial Subclusters

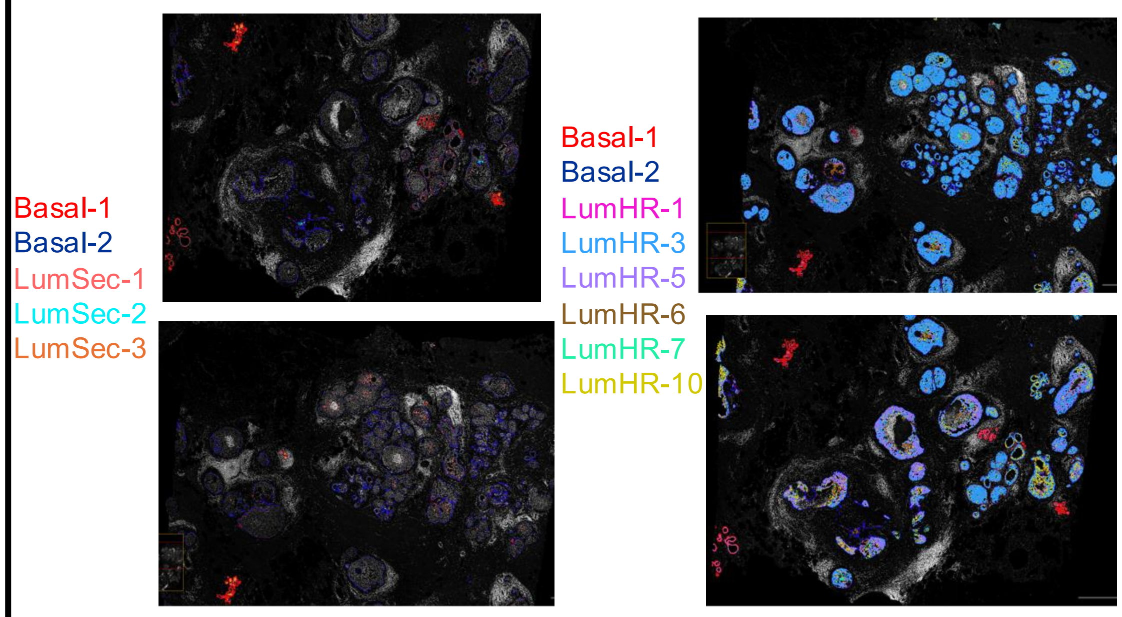


d. Graphical Representation of Epithelial Subcluster Interactions

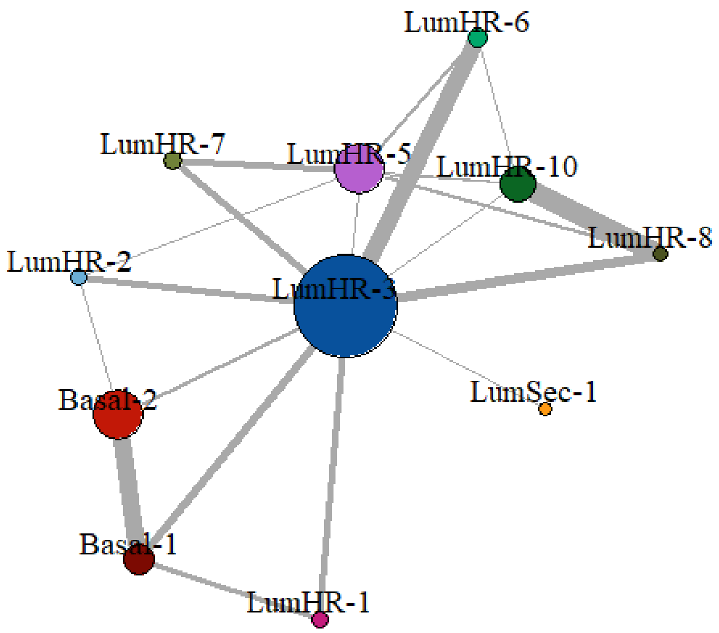

Supplementary Figure 1

### Xenium Sample 8: DCIS Luminal

#### a. Pathologist's Marked H&E

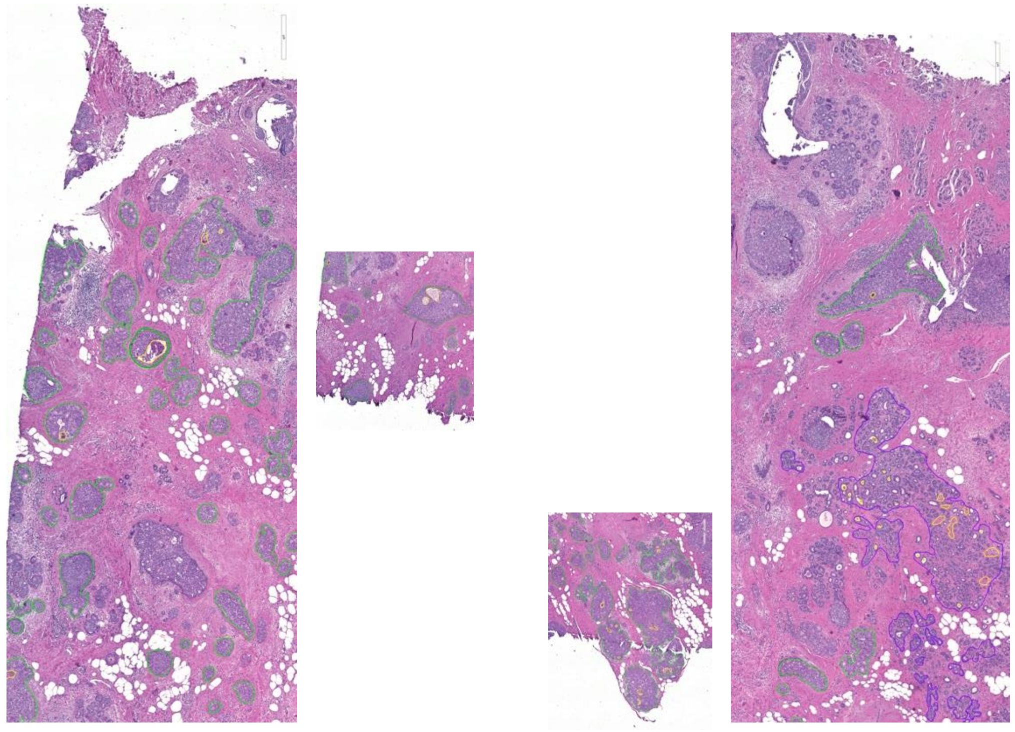

#### b. Proportion of Regional Epithelial Subclusters

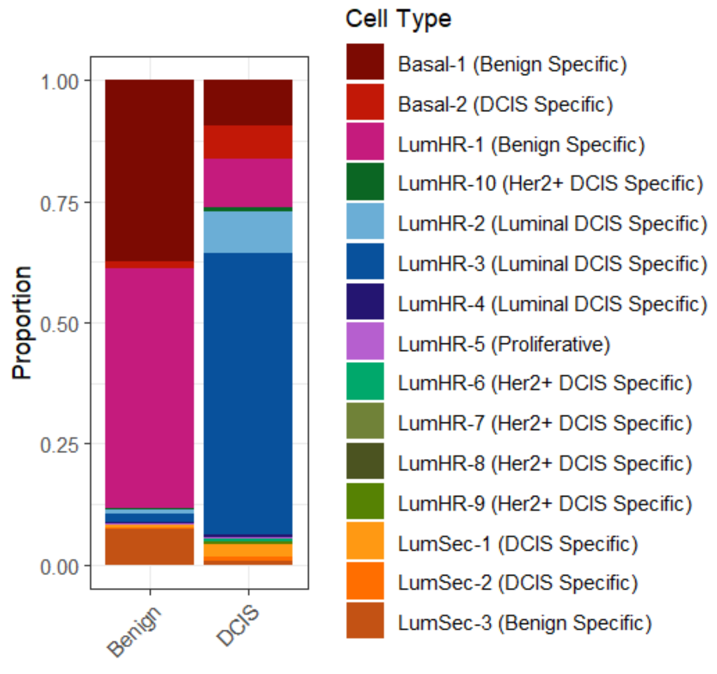


#### c. Spatial Organization of Epithelial Subclusters

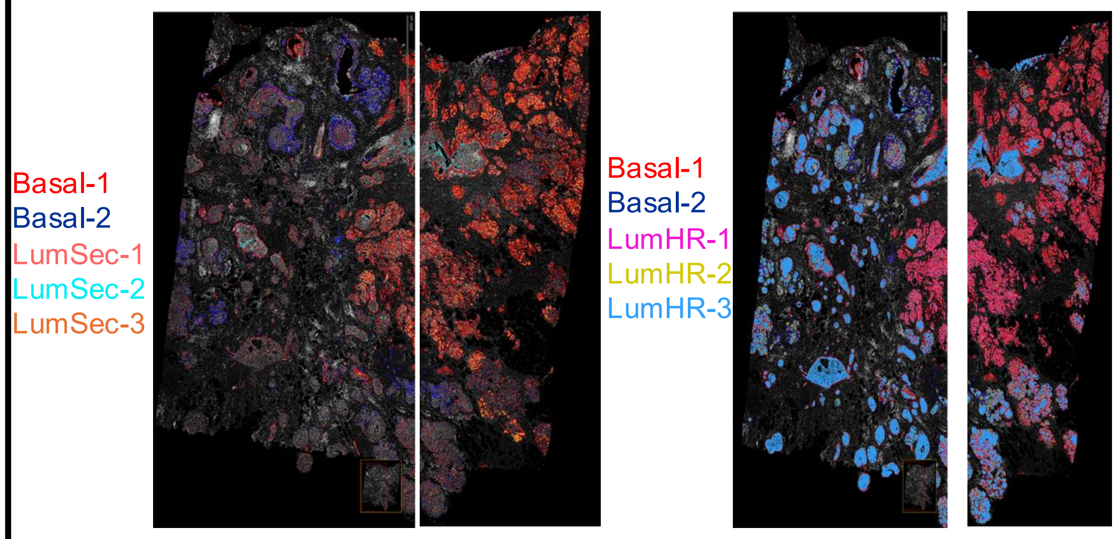


#### d. Graphical Representation of Epithelial Subcluster Interactions

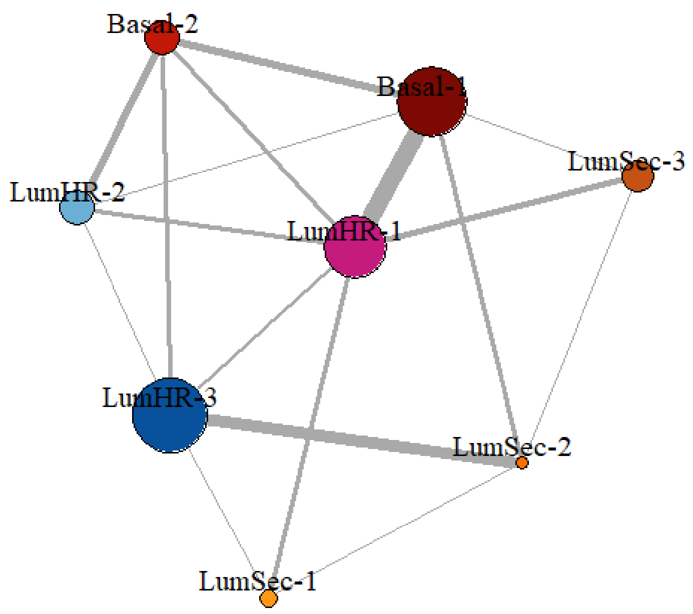

Supplementary Figure 1

### Xenium Sample 9: DCIS Luminal

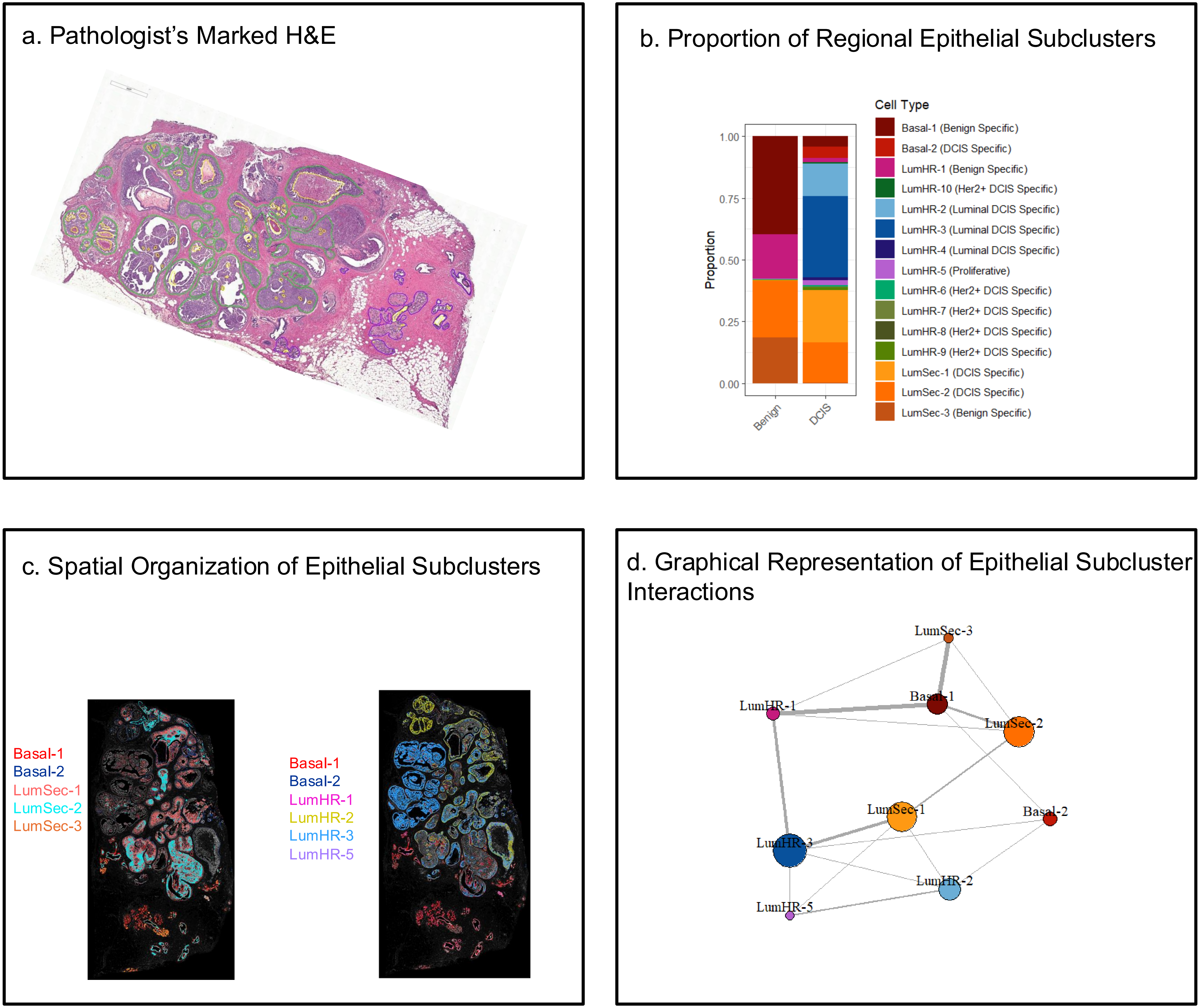

Supplementary Figure 1

### Xenium Sample 10: DCIS/IDC HER2+

#### a. Pathologist’s Marked H&E

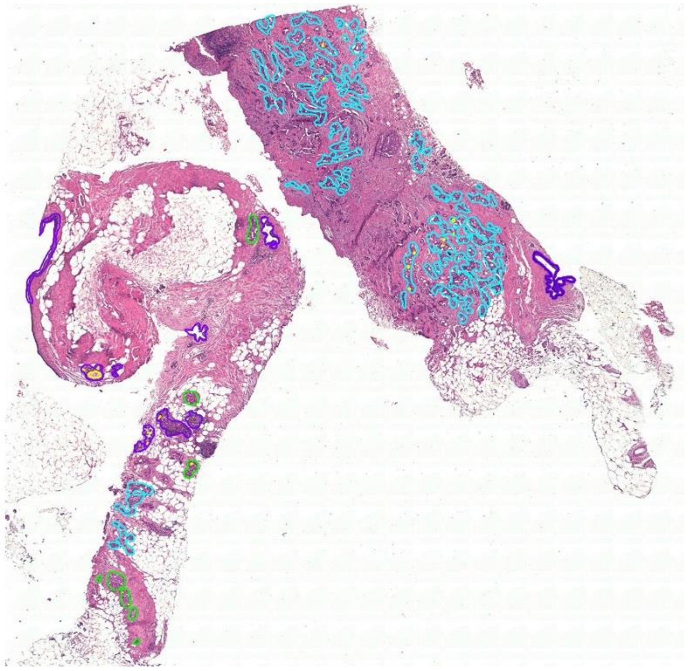

#### b. Proportion of Regional Epithelial Subclusters

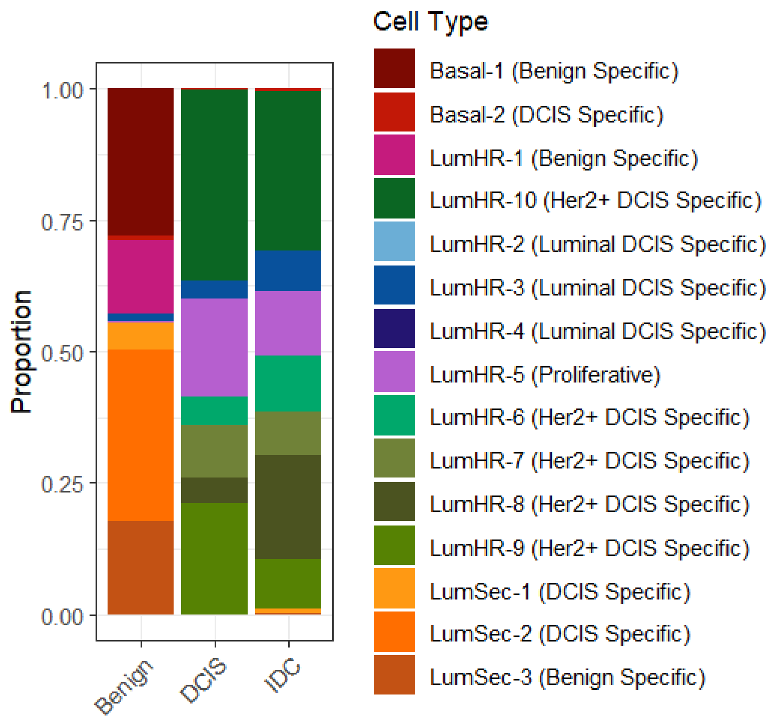


#### c. Spatial Organization of Epithelial Subclusters

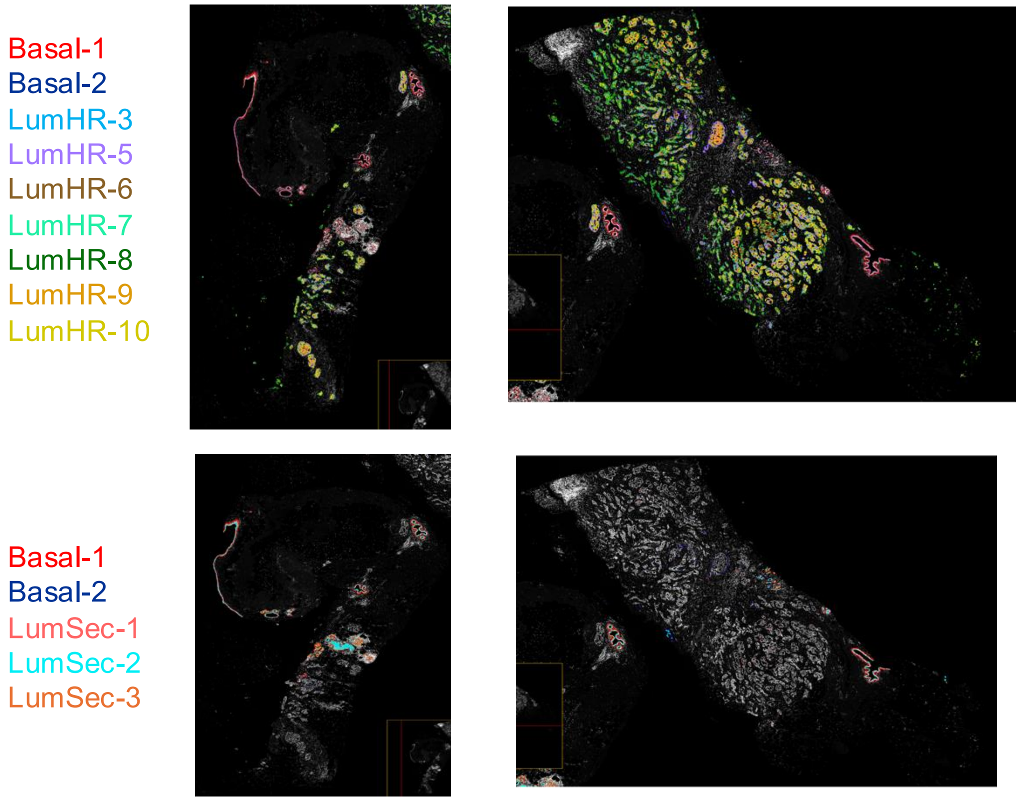


#### d. Graphical Representation of Epithelial Subcluster Interactions

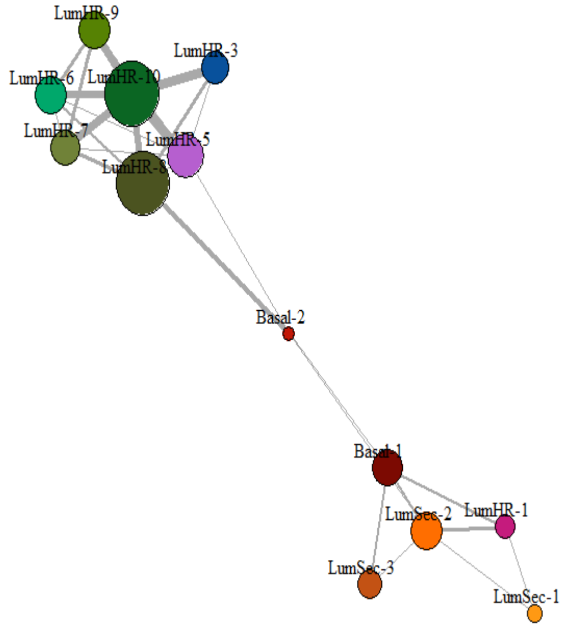

Supplementary Figure 1

### Xenium Sample 11: DCIS/IDC HER2+ DCIS (Lymph Node Metastasis)

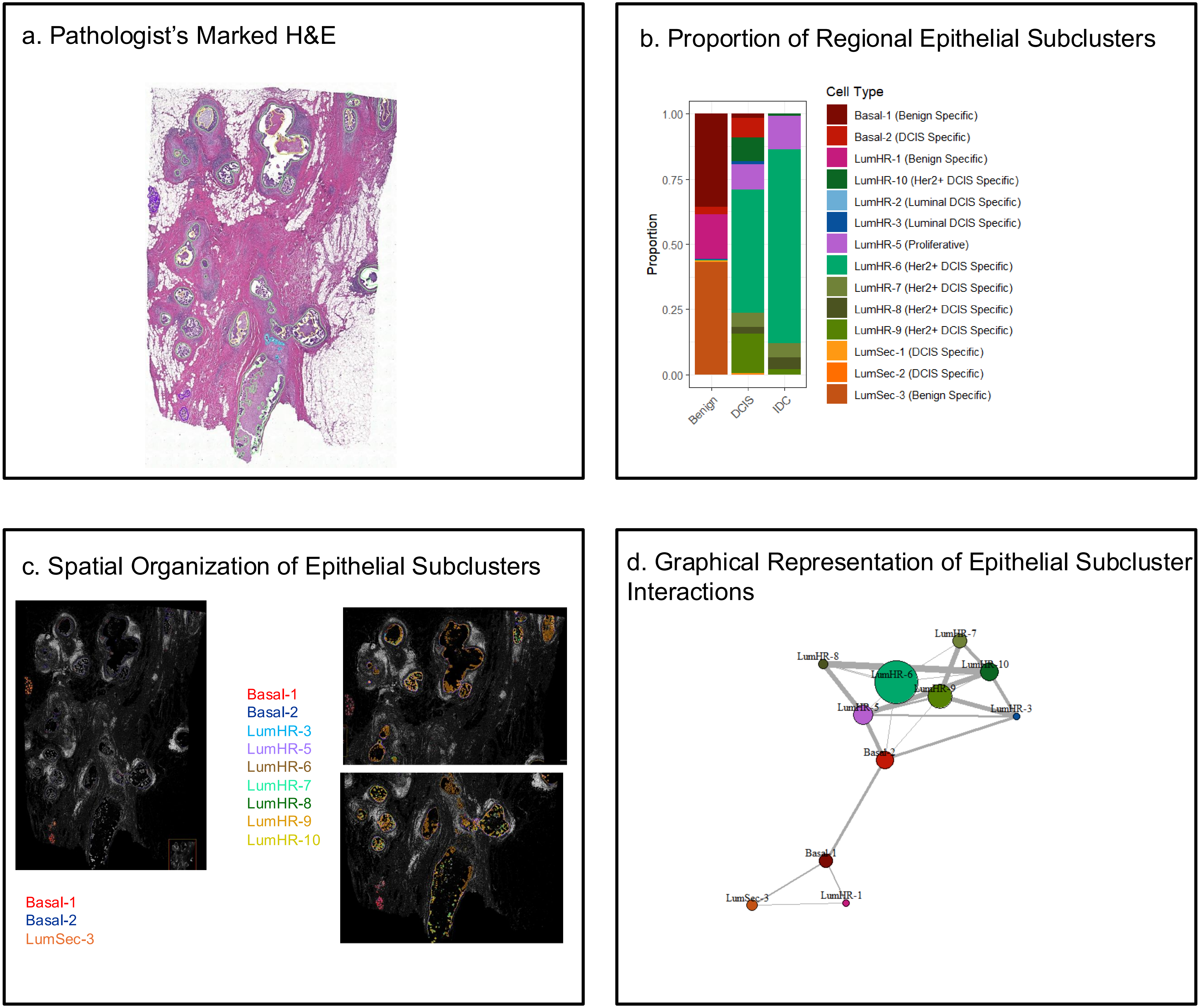

Supplementary Figure 1

### Xenium Sample 12: DCIS/IDC HER2+ (Lymph Node Metastasis)

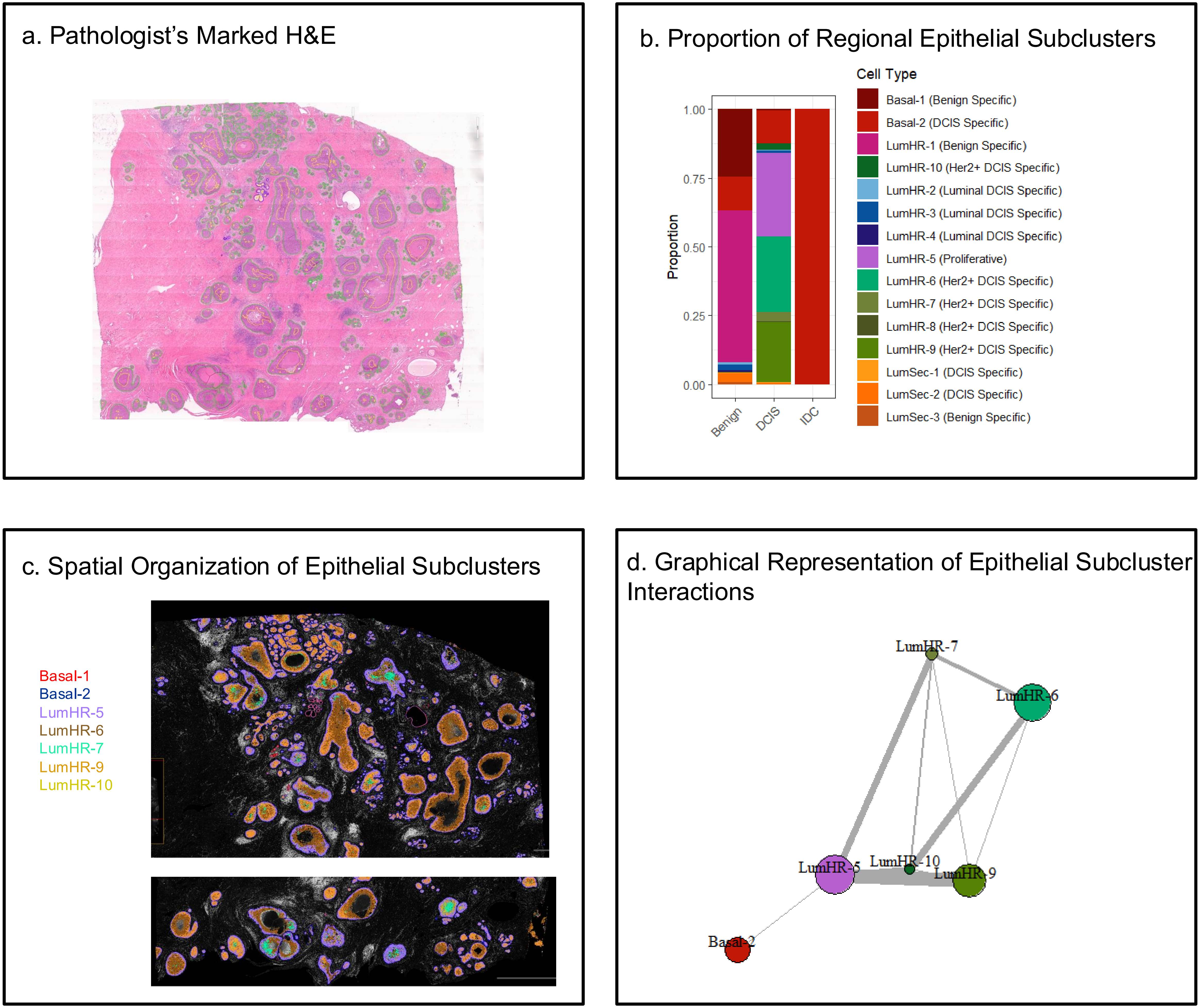

Supplementary Figure 1

### Xenium Sample 13: DCIS/IDC HER2+ (Lymph Node Metastasis)

#### a. Pathologist’s Marked H&E

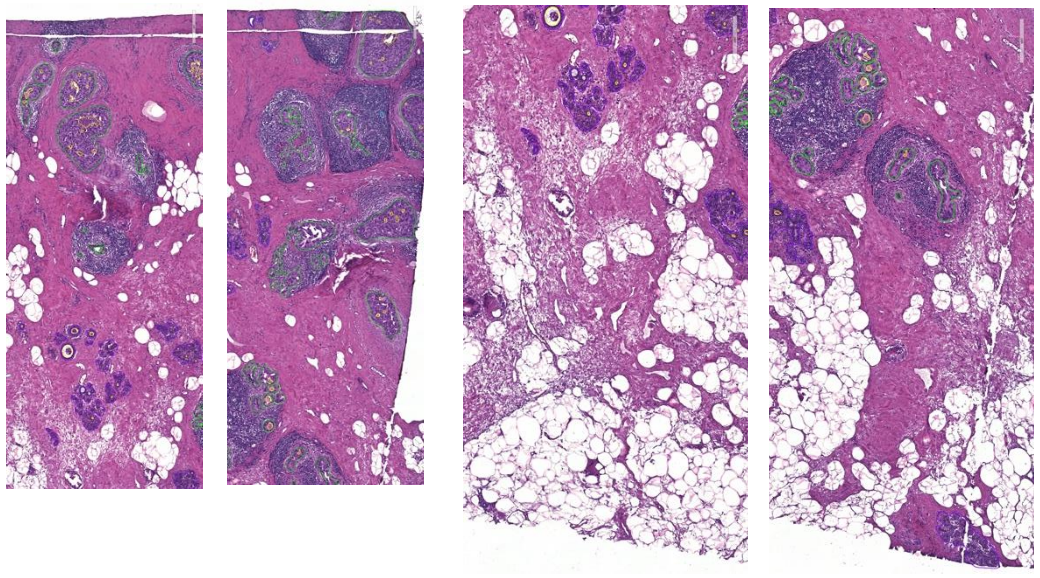

#### b. Proportion of Regional Epithelial Subclusters

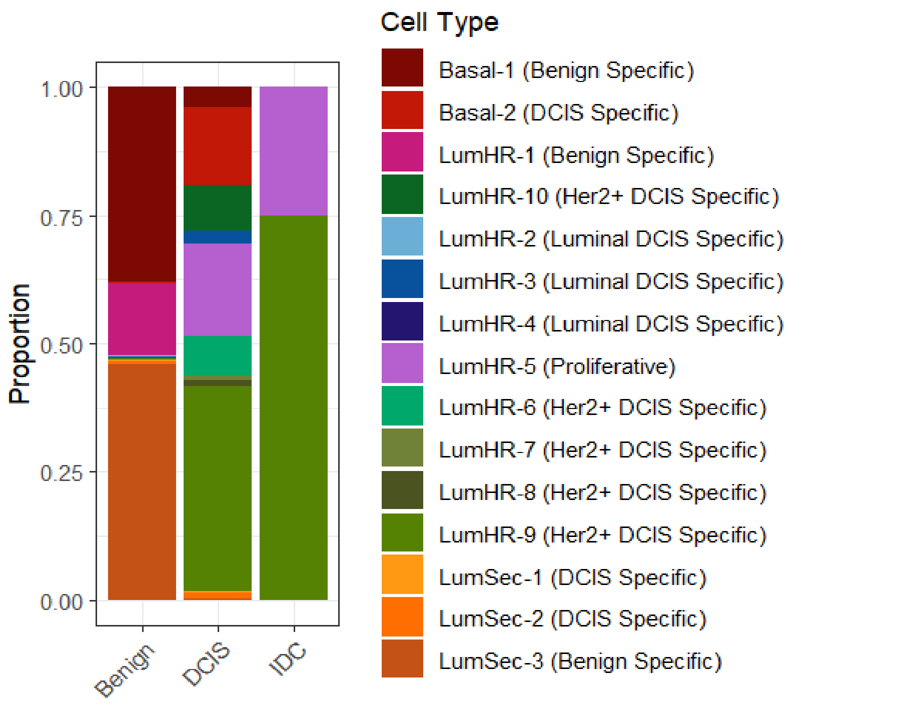


#### c. Spatial Organization of Epithelial Subclusters

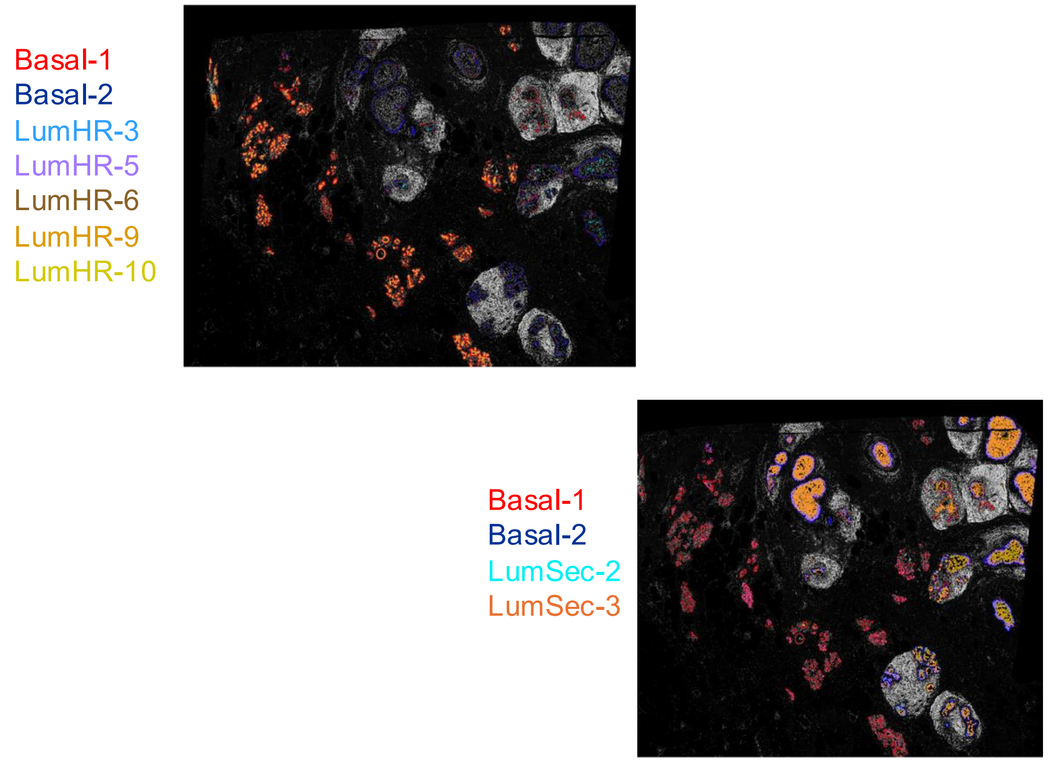


#### d. Graphical Representation of Epithelial Subcluster Interactions

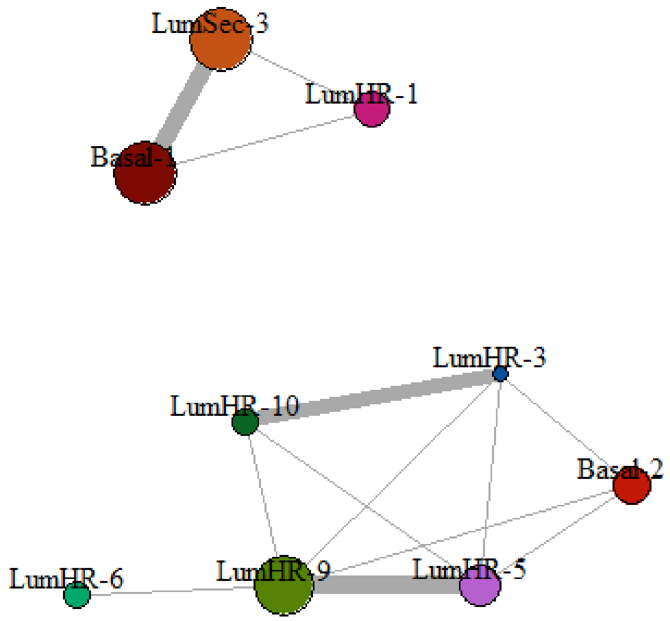

Supplementary Figure 1

### Xenium Sample 14: DCIS/IDC HER2+ DCIS

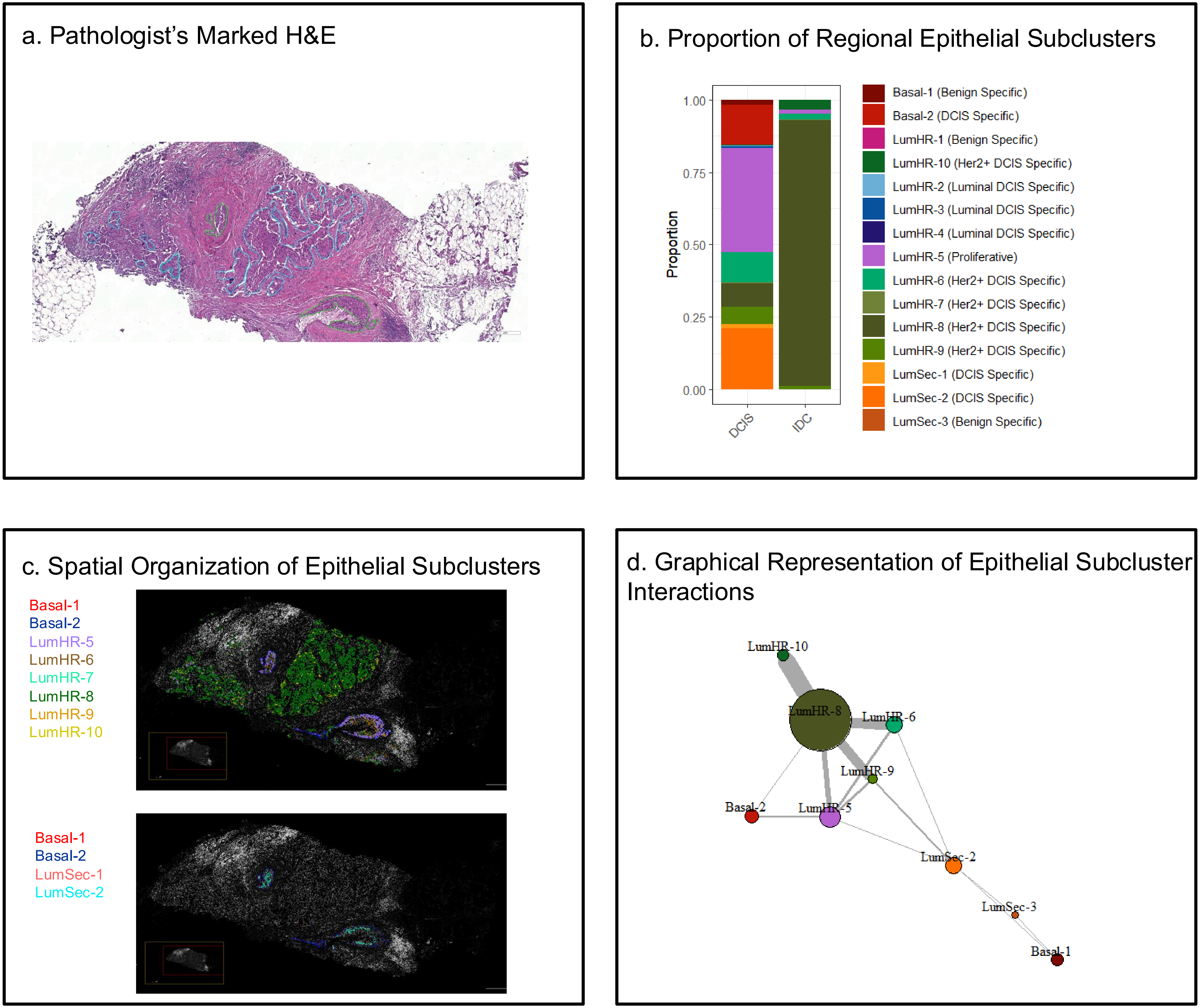

Supplementary Figure 1

### Xenium Sample 15: Luminal DCIS/IDC (Lymph Node Metastasis)

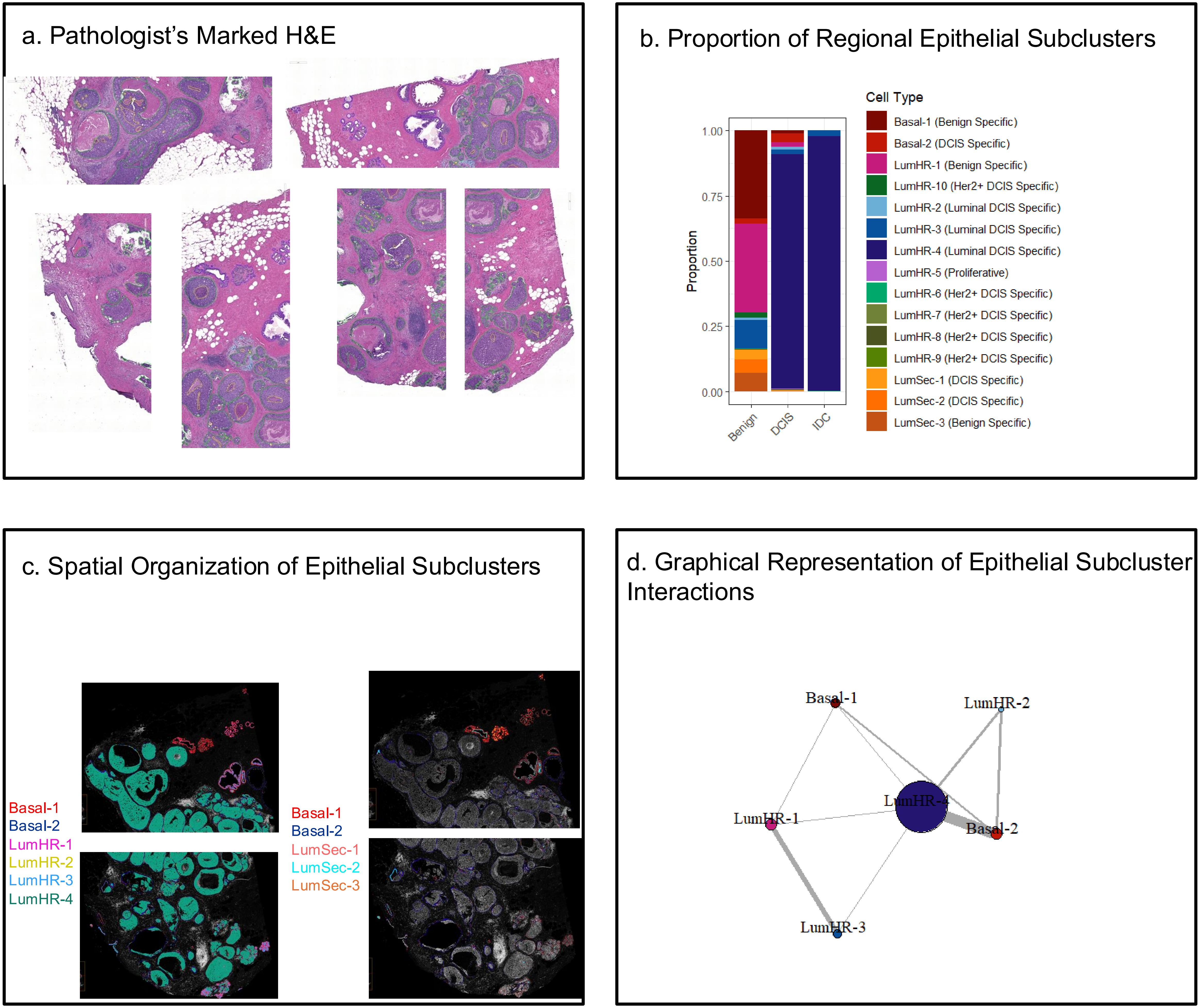

Supplementary Figure 1

### Xenium Sample 16: Luminal DCIS/IDC

a. Pathologist's Marked H&E

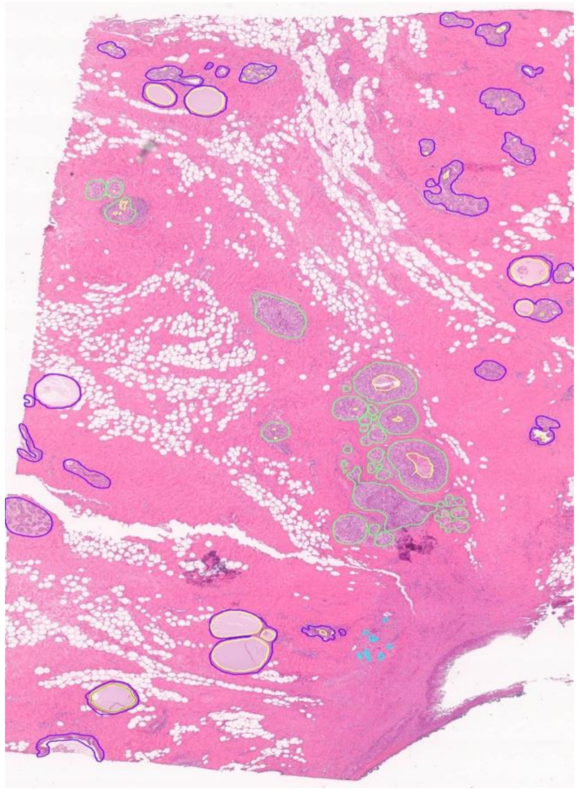

b. Proportion of Regional Epithelial Subclusters

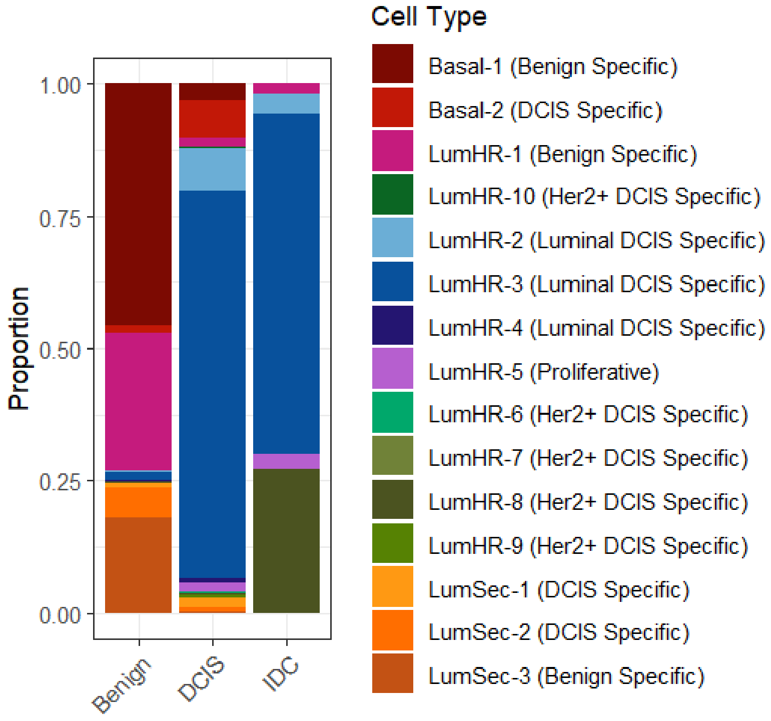


c. Spatial Organization of Epithelial Subclusters

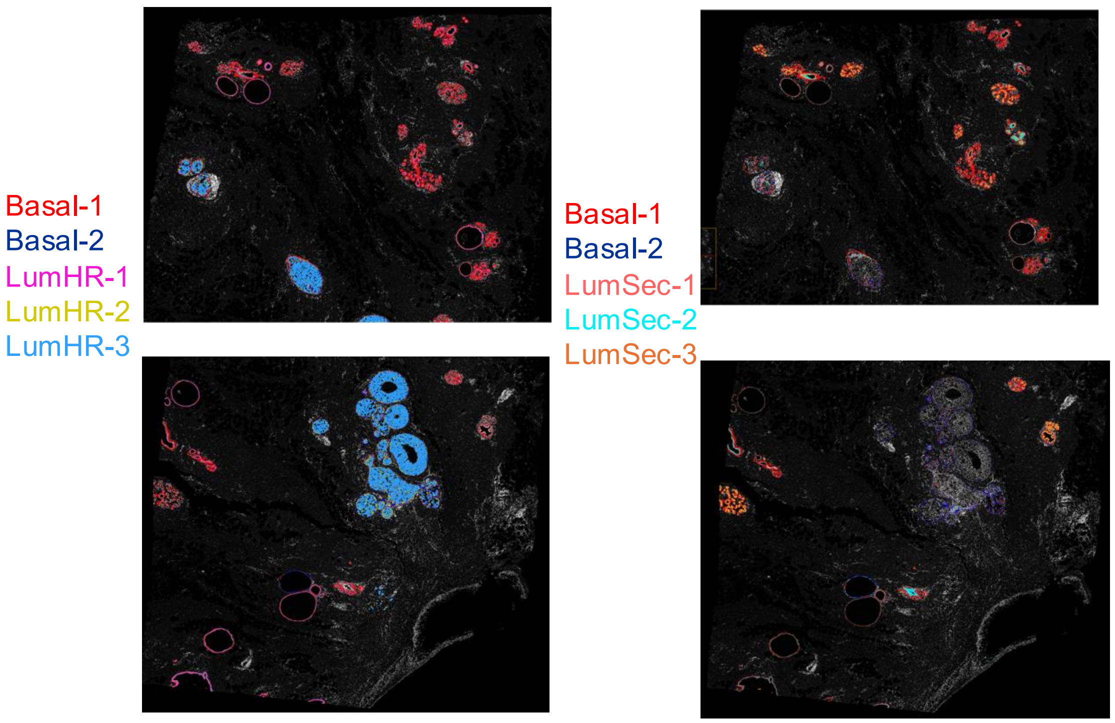


d. Graphical Representation of Epithelial Subcluster Interactions

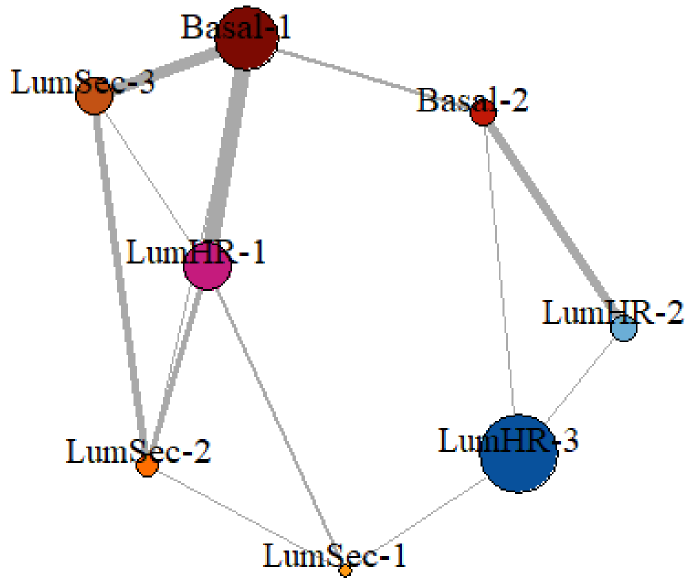

**Supplementary Figure 1 | Spatial and cellular organization of epithelial subclusters across benign, DCIS and IDC regions in the 16 Xenium Samples.**

**a,** Haematoxylin and eosin (H&E)–stained sections from all 16 Xenium samples annotated by a breast pathologist, indicating benign epithelium, ductal carcinoma in situ (DCIS) and invasive ductal carcinoma (IDC) regions.

**b,** Proportional distribution of epithelial subclusters within each pathologist-annotated region (benign, DCIS and IDC) across all samples.

**c,** Spatial localization of epithelial subclusters overlaid on tissue sections, illustrating region-specific and subtype-specific epithelial organization.

**d,** Networks showing cell-cell interactions among epithelial subclusters, highlighting co-occurrence and spatial proximity patterns across samples. Size of the node is proportional to the abundance of the cell type in that sample. Width of the edge (lines) between nodes is proportional to the amount of interactions between two cell types.

### Supplementary Figure 2

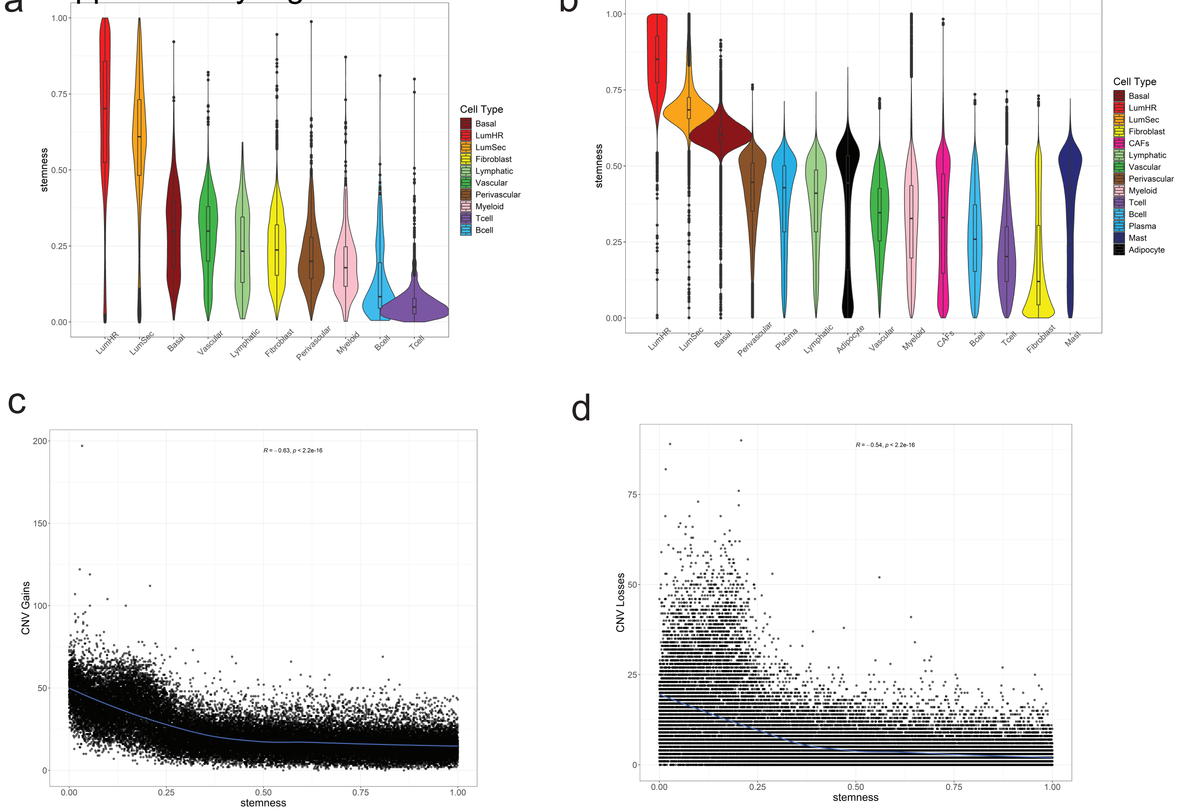

**Supplementary Figure 2 | Stemness, copy number variation and meta-program associations in MIND model epithelial cells.**
**a,** Distribution of CytoTRACE stemness scores across major epithelial and stromal cell populations identified by single-cell RNA sequencing in the MIND model, showing highest stemness in luminal hormone-responsive (LumHR) cells.
**b,** Comparison of CytoTRACE stemness scores among epithelial subtypes in Xenium samples, confirming enrichment of stem-like features in LumHR cells relative to basal and luminal secretory (LumSec) populations.
**c,d,** Spearman correlations between CytoTRACE stemness scores and inferred copy number gains (**c**) or copy number losses (**d**) in LumHR cells, demonstrating a negative association between stemness and genomic instability.

### Supplementary Figure 3

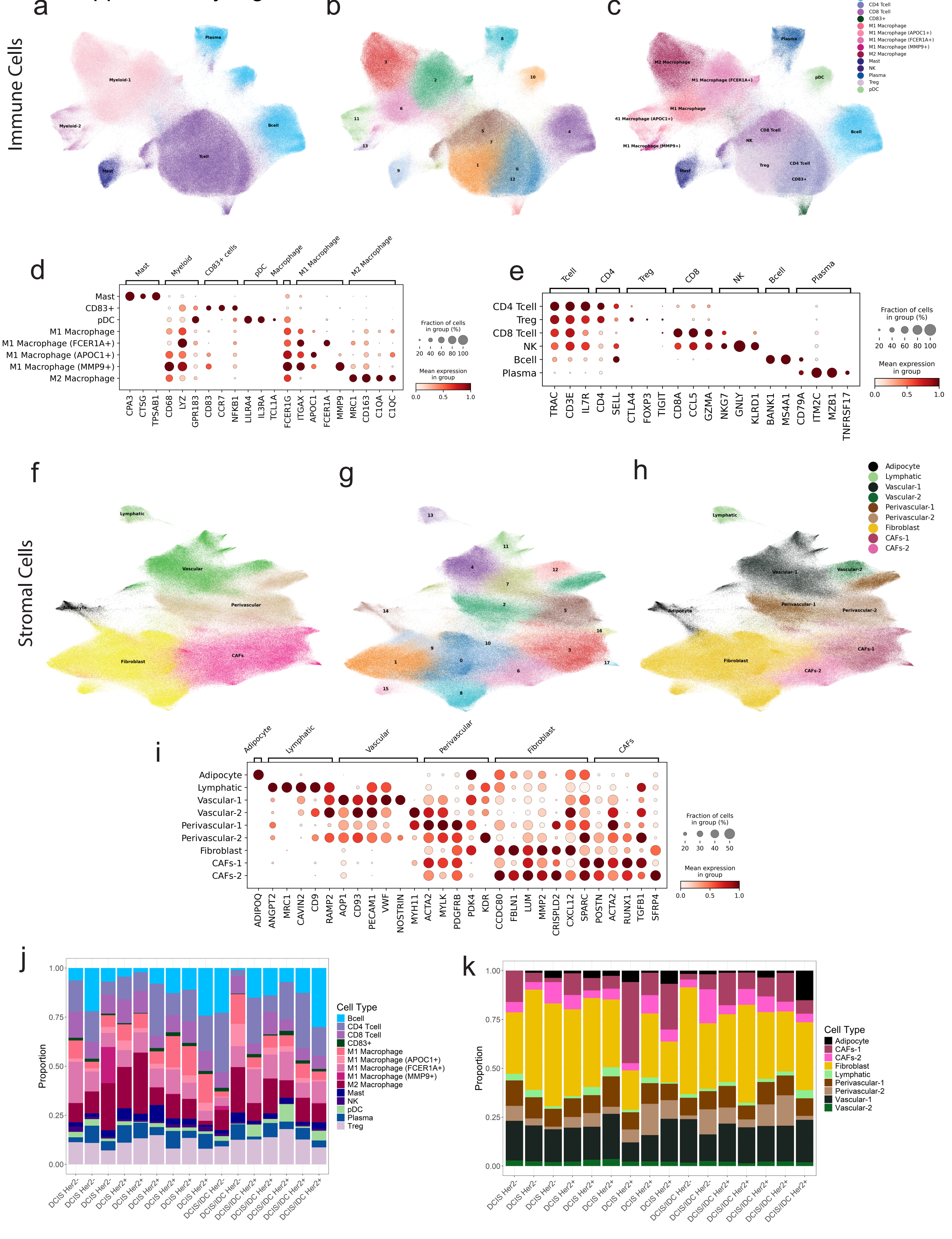

**Supplementary Figure 3 | Identification of immune and stromal populations in Xenium samples**

**a,** UMAP of integrated immune cells, labeled by their identity in Figure 2a.
**b,** UMAP of re-clustered immune cells
**c,** UMAP of re-labeled immune cells
**d,** Dot plot showing marker genes to identify myeloid populations
**e,** Dot plot showing marker genes to identify lymphocyte populations
**f,** UMAP of integrated stromal cells, labeled by their identity in Figure 2a.
**g,** UMAP of re-clustered stromal cells
**h,** UMAP of re-labeled stromal cells
**i,** Dot plot showing marker genes to identify stromal cells
**j,** Bar plot of the proportion of immune cells across all 16 samples
**k,** Bar plot of the proportion of stromal cells across all 16 samples

#### Supplementary Figure 4

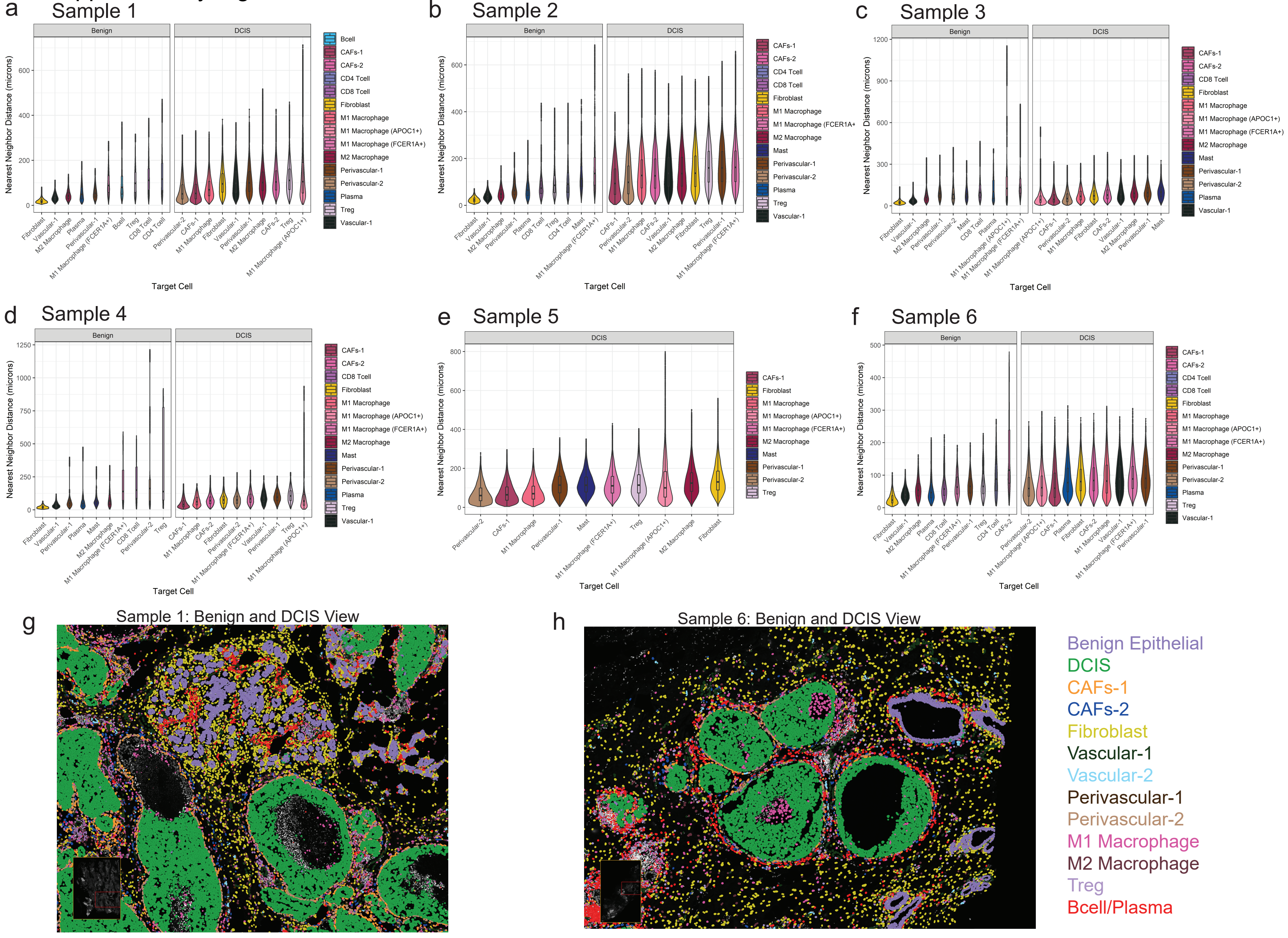

**Supplementary Figure 4 | Tumor microenvironment nearest-neighbors in HER2+ DCIS samples.**

**a-f,** Violin plots showing the top 10 cells with the smallest average distance to benign epithelium or DCIS lesions for each individual sample. Each panel represents an individual sample. Each sub-panel represents the distance (y-axis) between either benign epithelium or DCIS cells to specific target cells (x-axis). *Sample 5 did not have any benign epithelium.

**g,** View of DCIS lesions and benign epithelium and surrounding cells from sample 1. This is a zoomed in area from the entire sample to show the localization of the nearest cell types to epithelial cells.

**h,** View of DCIS lesions and benign epithelium and surrounding cells from sample 6. This is a zoomed in area from the entire sample to show the localization of the nearest cell types to epithelial cells.

### Supplementary Figure 5

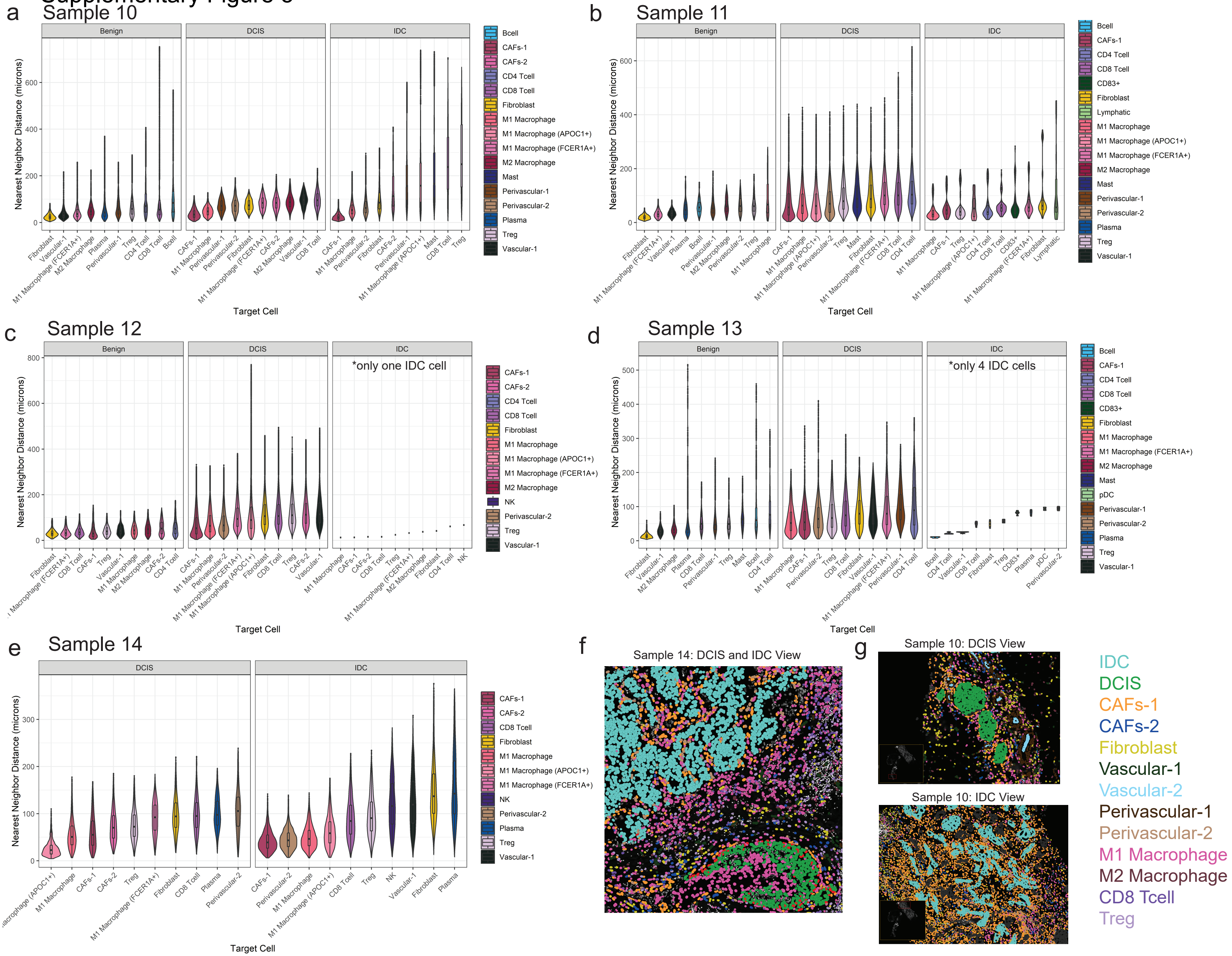

**Supplementary Figure 5 | Tumor microenvironment nearest-neighbors in HER2+ DCIS/IDC samples.**

**a-e,** Violin plots showing the top 10 cells with the smallest average distance to benign epithelium, DCIS lesions, or IDC for each individual sample. Each panel represents an individual sample. Each sub-panel represents the distance (y-axis) between either benign epithelium, DCIS cells, or IDC cells to specific target cells (x-axis). *Sample 12 only has one IDC cell. Sample 13 only has 4 IDC cells. Sample 14 did not have any benign epithelium.

**f,** View of DCIS lesion and IDC region and surrounding cells from sample 14. This is a zoomed in area of the entire sample to show the localization of the nearest cell types to the cancer cells.

**g,** View of DCIS lesions (top) and IDC region (bottom) and surrounding cells from sample 10. These are zoomed in areas from the entire sample to show the localization of the nearest cell types to the cancer cells.

Supplementary Figure 6

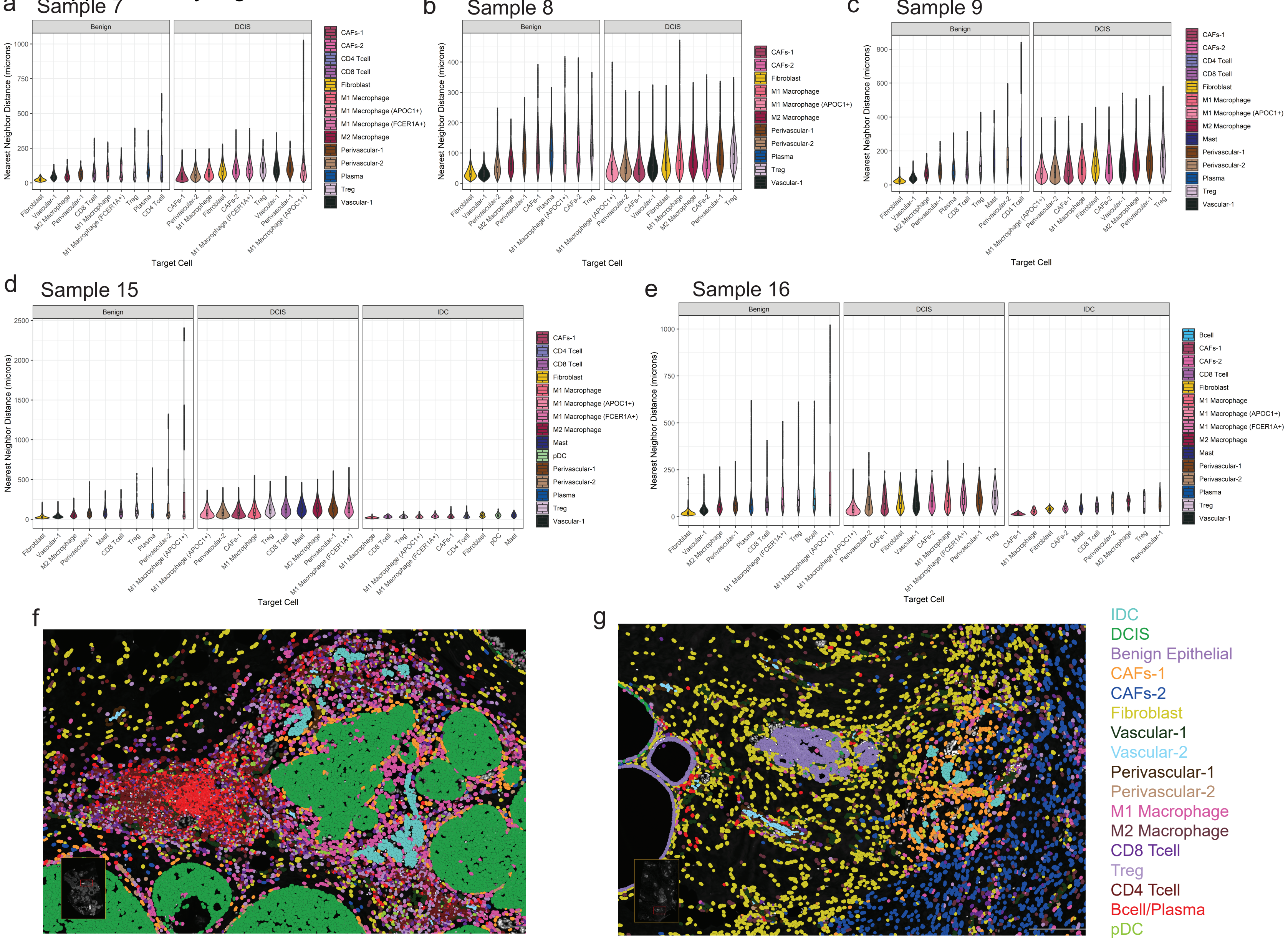

**Supplementary Figure 6 | Tumor microenvironment nearest-neighbors in Luminal DCIS (a-c) and DCIS/IDC (d-e) samples.**

**a-e,** Violin plots showing the top 10 cells with the smallest average distance to benign epithelium, DCIS lesions, or IDC for each individual sample. Each panel represents an individual sample. Each sub-panel represents the distance (y-axis) between either benign epithelium, DCIS cells, or IDC cells to specific target cells (x-axis).

**f,** View of IDC cells and DCIS lesions and surrounding cells from sample 15. This is a zoomed in area of the entire sample to show the localization of the nearest cel types to the cancer cells.

**g,** View of IDC cells and benign epithelium and surrounding cells from sample 16. This is a zoomed in area of the entire sample to show the localization of the nearest cel types to the cancer cells.

### Supplementary Figure 7

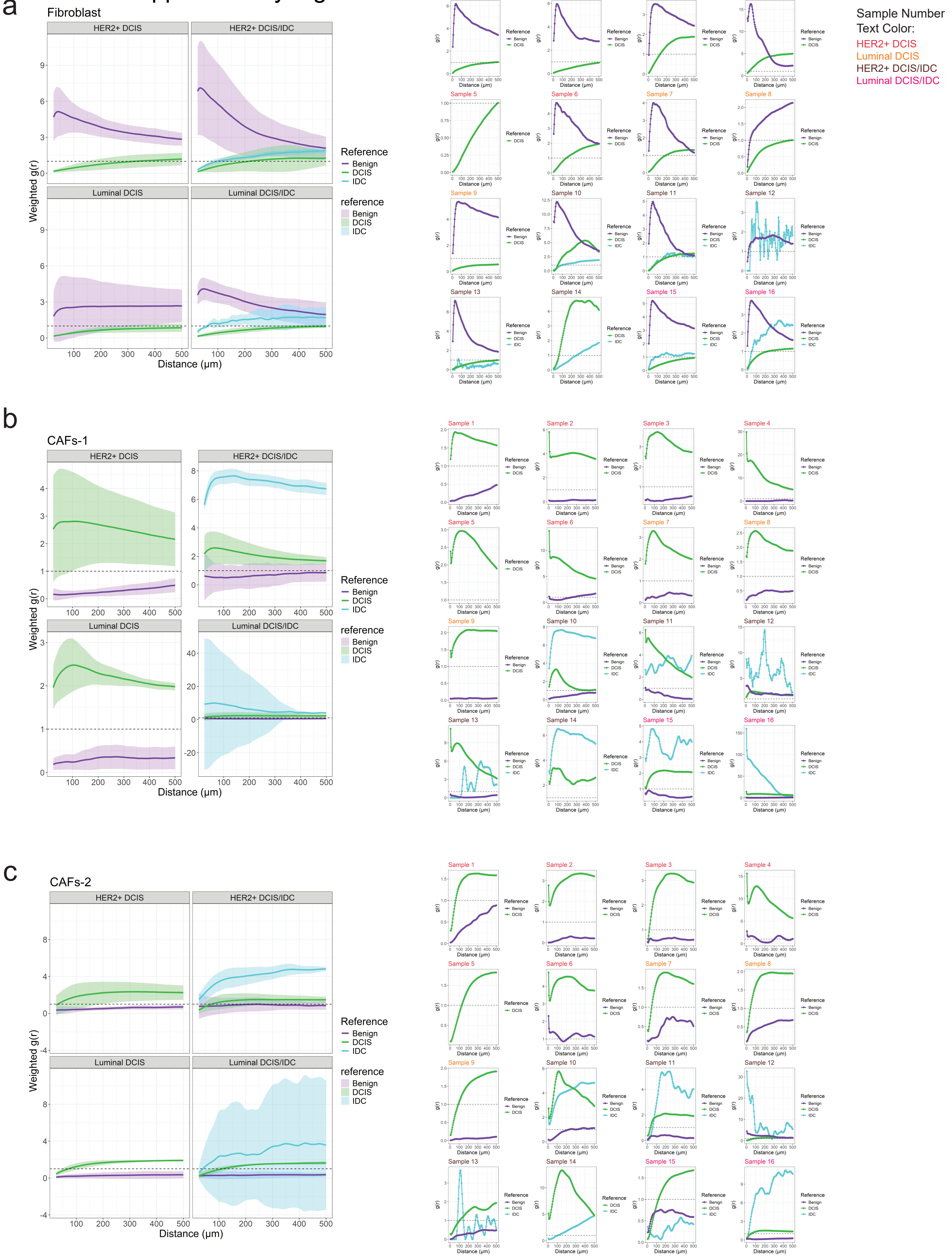

### Supplementary Figure 7

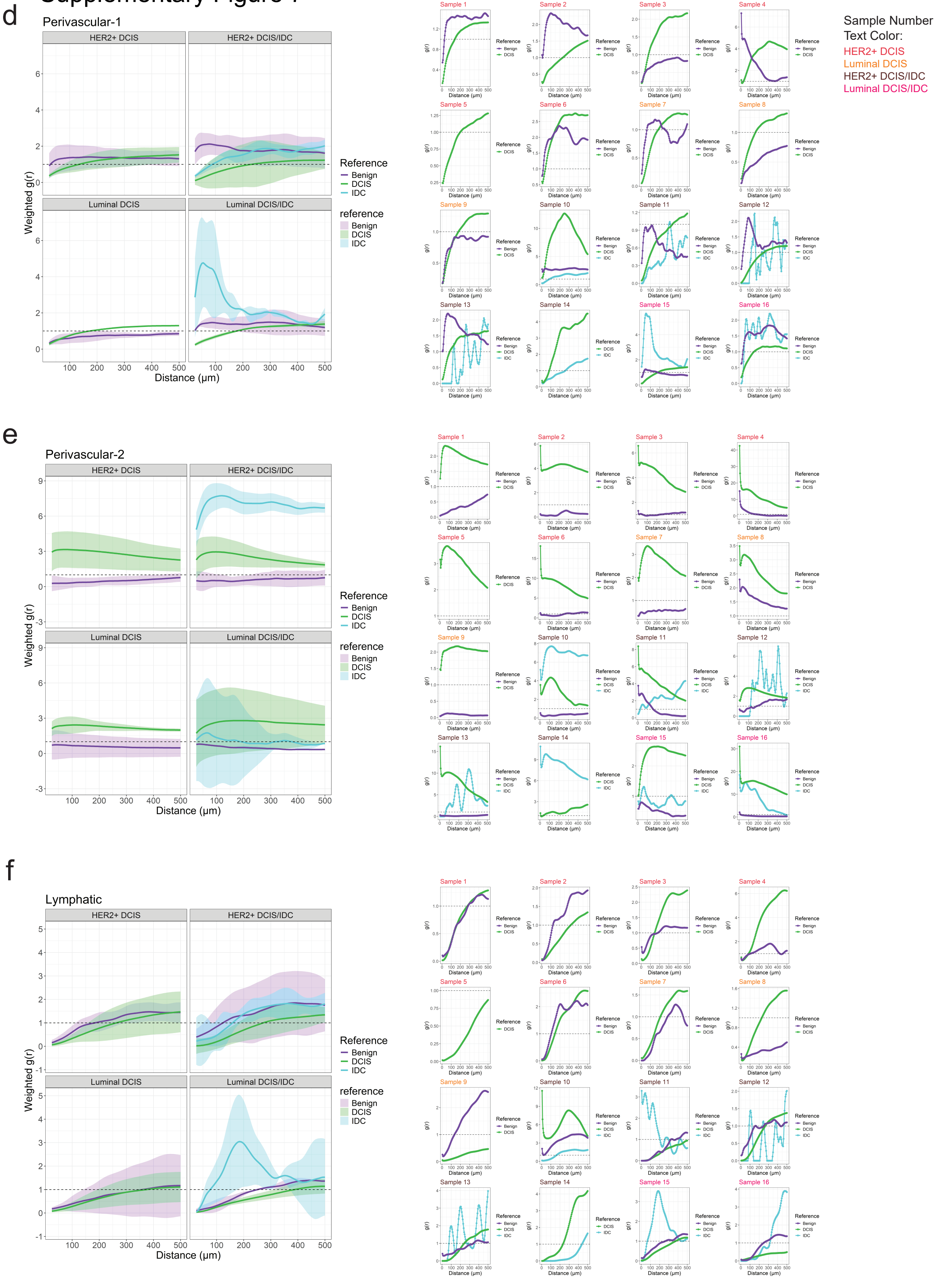

### Supplementary Figure 7

g

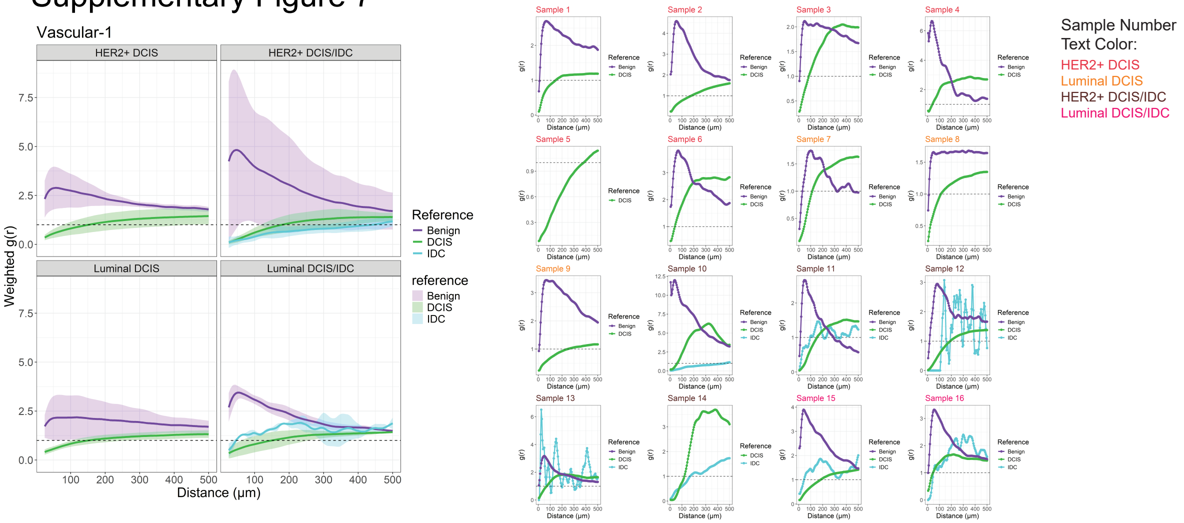


h

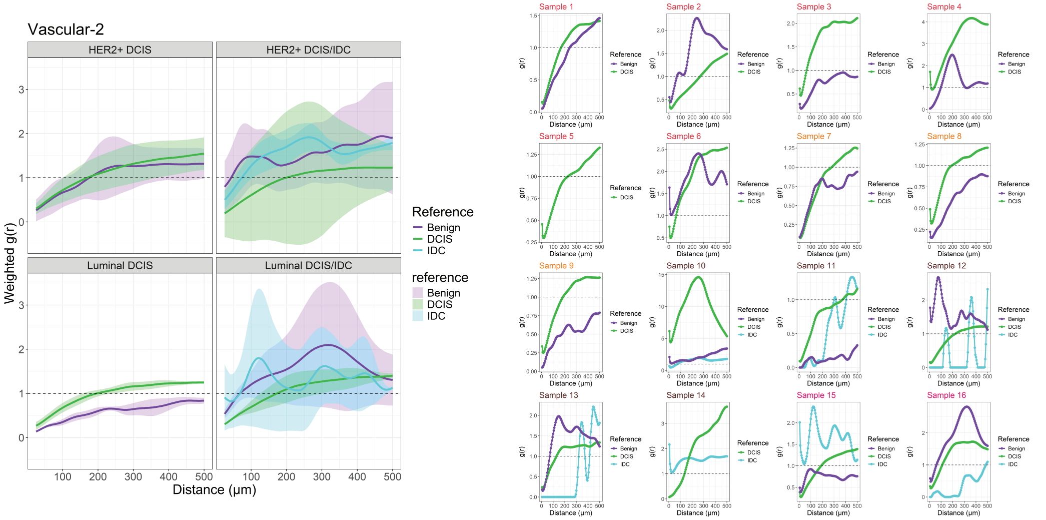


i

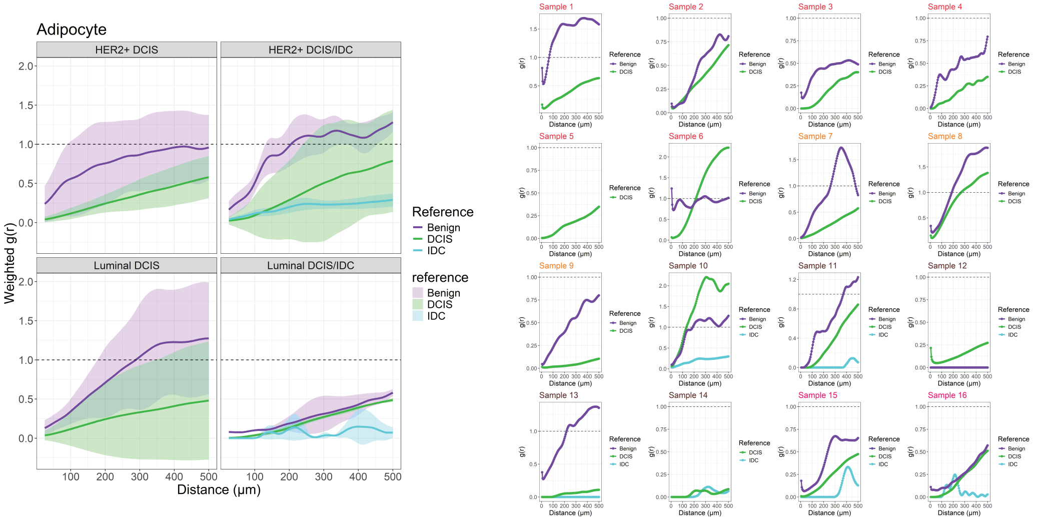

### Supplementary Figure 7

j

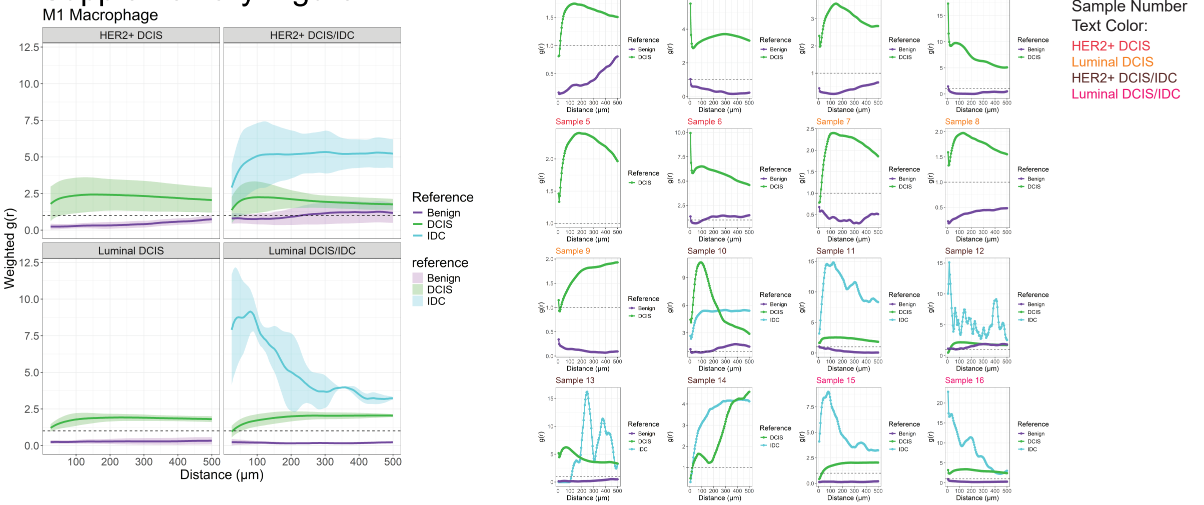


k

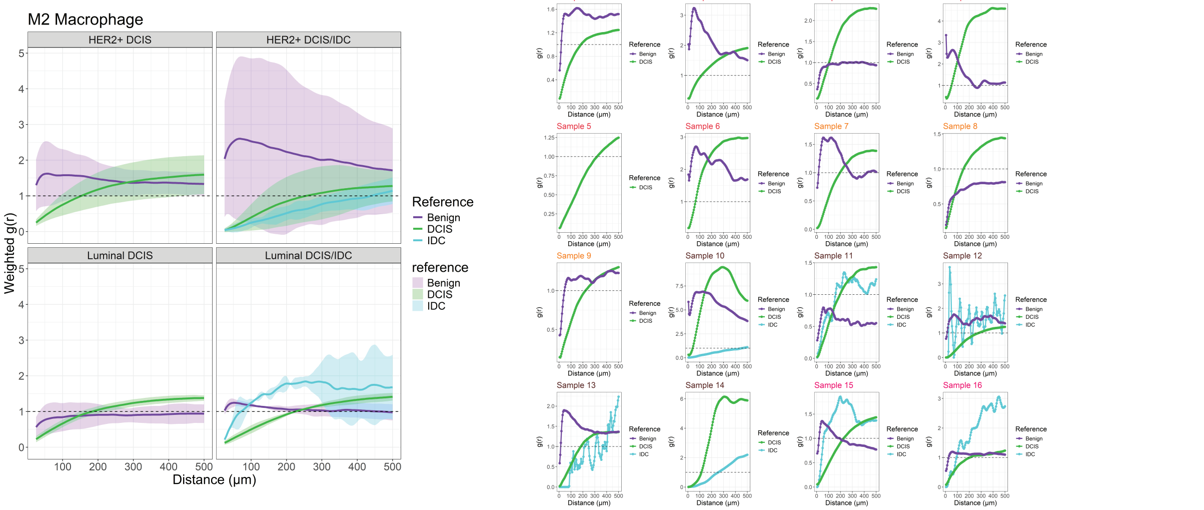


l

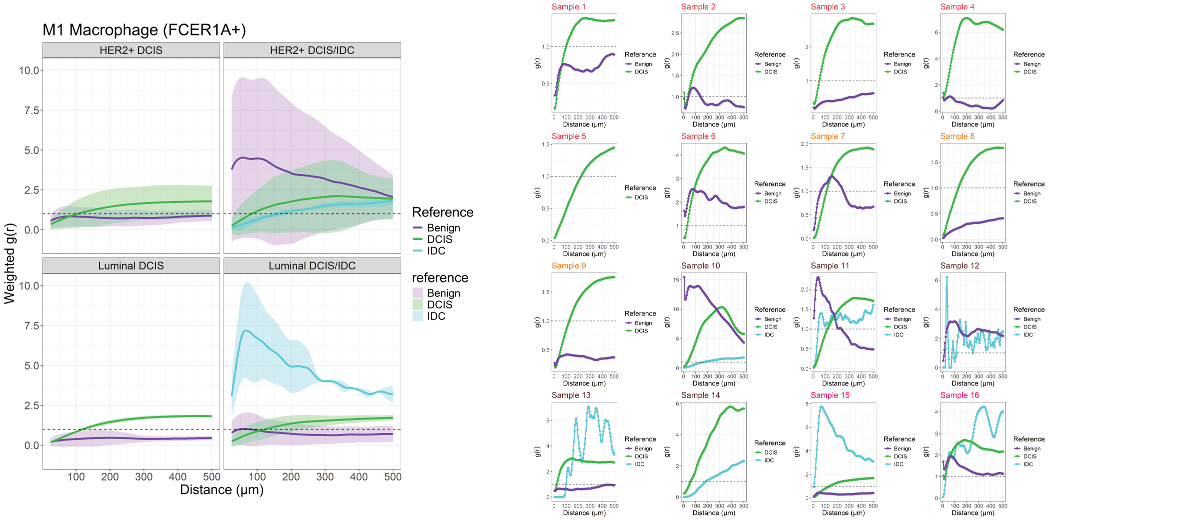

### Supplementary Figure 7

m

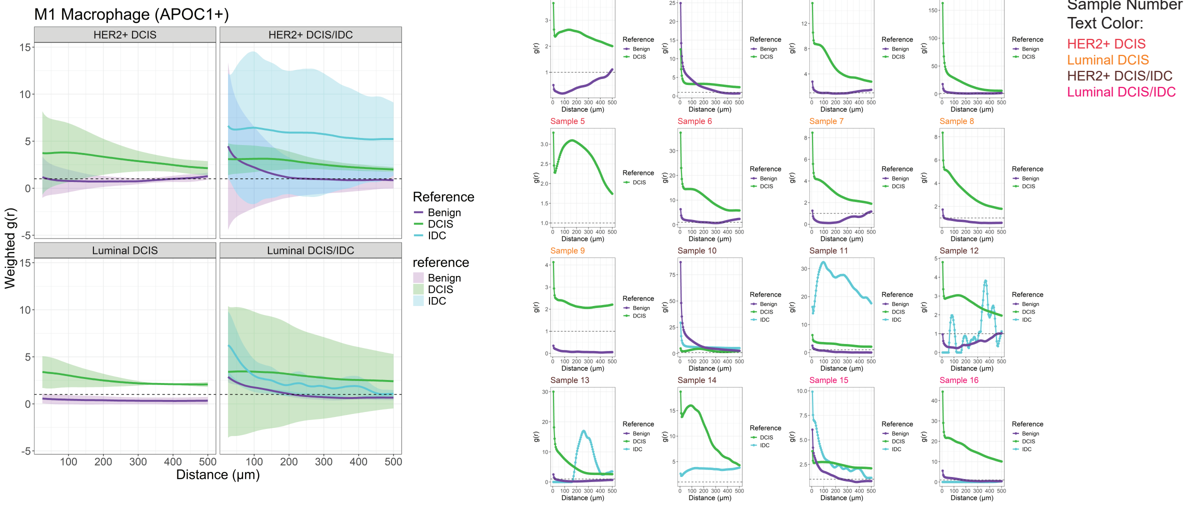


n

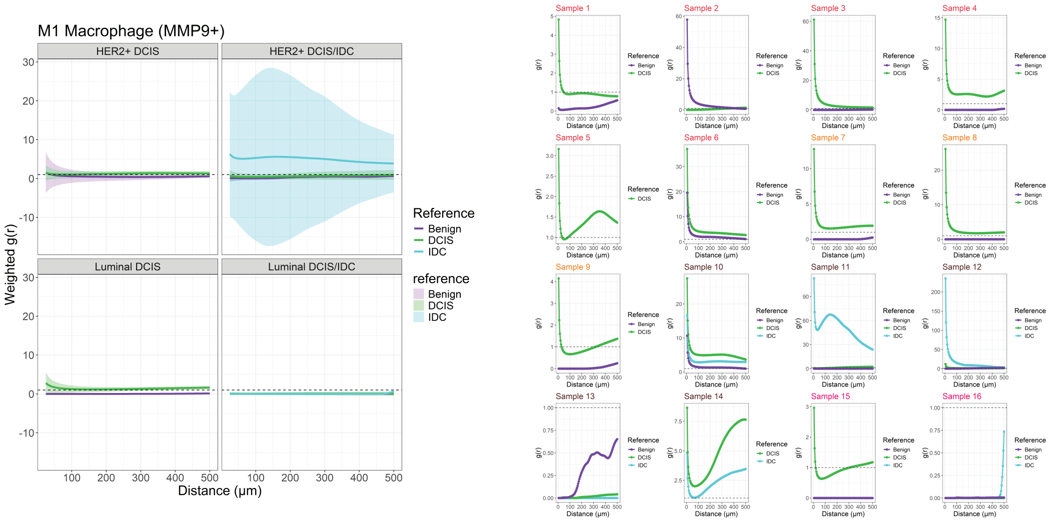


o

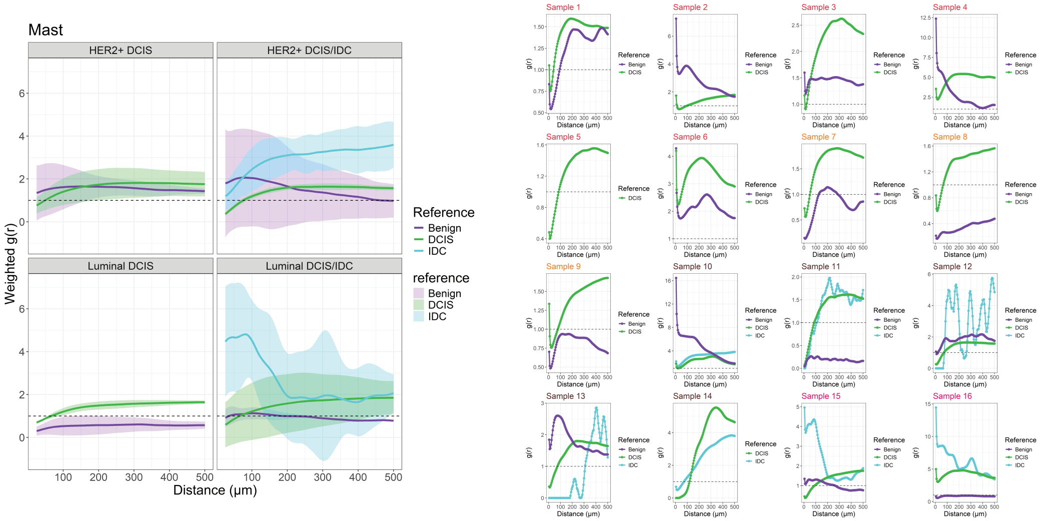

### Supplementary Figure 7

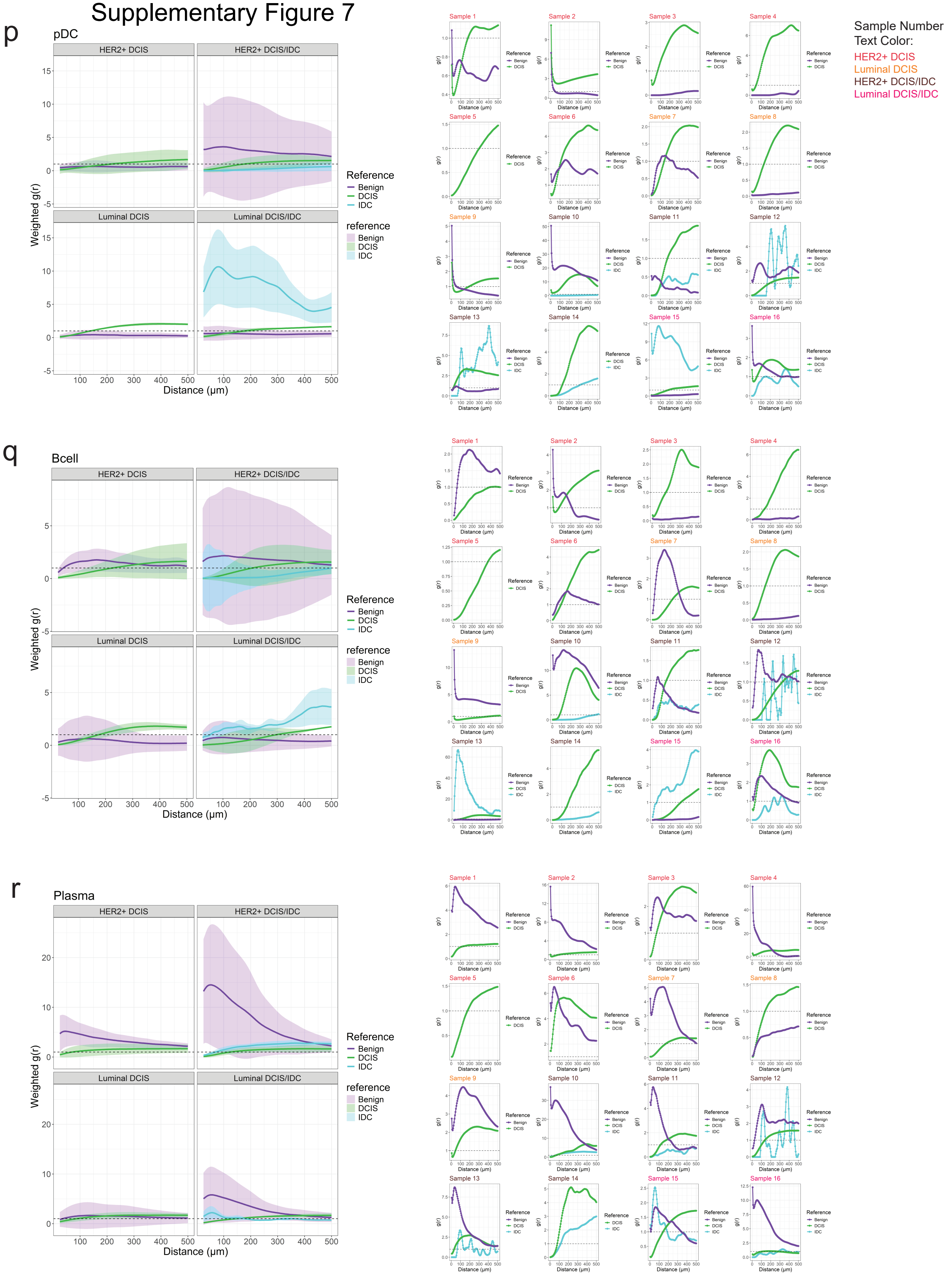

### Supplementary Figure 7

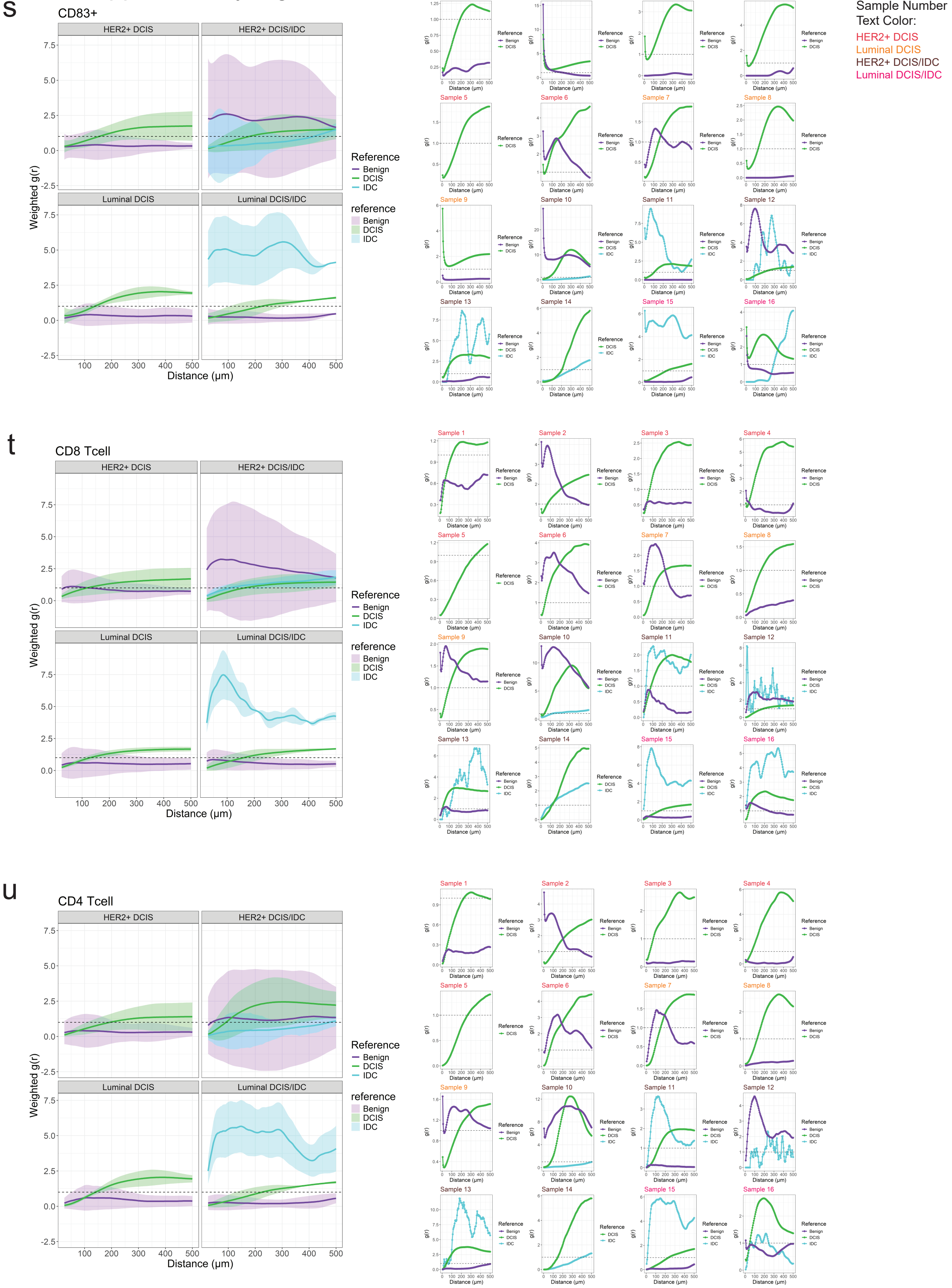

### Supplementary Figure 7

V

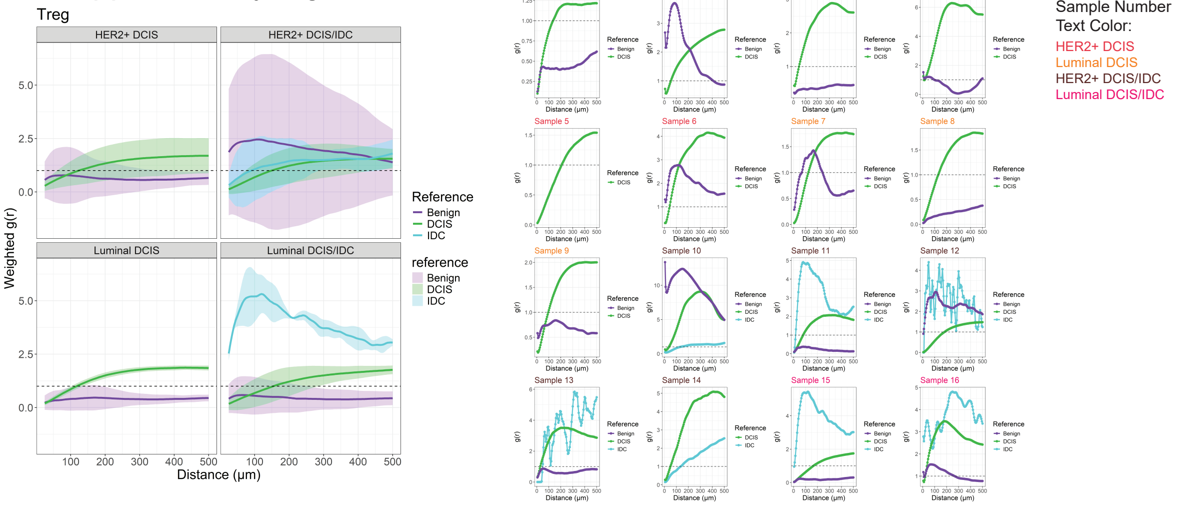


W

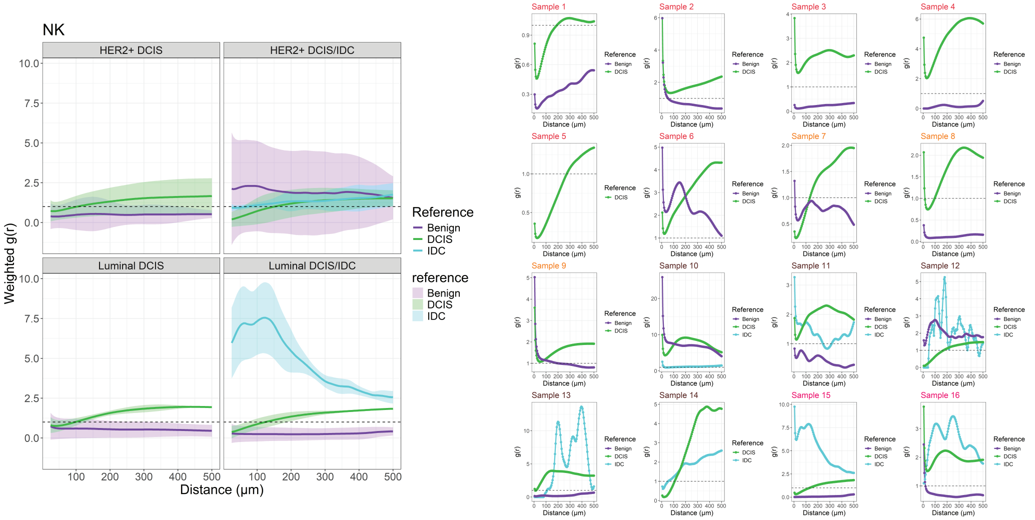

**Supplementary Figure 7 | Tumor microenvironment cross-type pair correlation functions in all samples for all cell types (a-w).** Each panel represents an evaluated cell type. **The left plot** of each panel shows the weighted average cross-type pair correlation function, $g(r)$ (y-axis), for HER2+ DCIS, HER2+ DCIS/IDC, Luminal DCIS, and Luminal DCIS/IDC samples. We evaluated $g(r)$ for the cell type of interest in the panel and either benign epithelium, DCIS cells, or IDC cells, represented by the different color lines (Benign = purple, DCS = green, IDC = blue). Shaded regions indicate 95% confidences intervals. Distances plotted range from 20 µm to 500 µm by increments of 5 µm (x-axis). **The right plot** of each panel shows the cross-type pair correlation function (y-axis) for each sample individually, for the same cell type of interest and one of the three types of epithelial cells. Distances plotted range from 5 µm to 500 µm by increments of 5 µm (x-axis). Sample labels are colored by their respective group (red=HER2+ DCIS, orange=Luminal DCIS, brown=HER2+ DCIS/IDC, pink=Luminal DCIS/IDC). A $g(r) = 1$ indicates independence of the two cell types, meaning the cell types are randomly distributed; $g(r) > 1$ indicates there is more clustering of the two cell types than expected by random chance; $g(r) < 1$ indicates the two cell types are more dispersed from each other than expected by random chance. The horizontal dashed line is at $g(r) = 1$.
*Sample 12 only has one IDC cell. Sample 13 only has 4 IDC cells.
*There are only two Luminal DCIS/IDC samples and can result in unstable weighted average $g(r)$.

Supplementary Figure 8

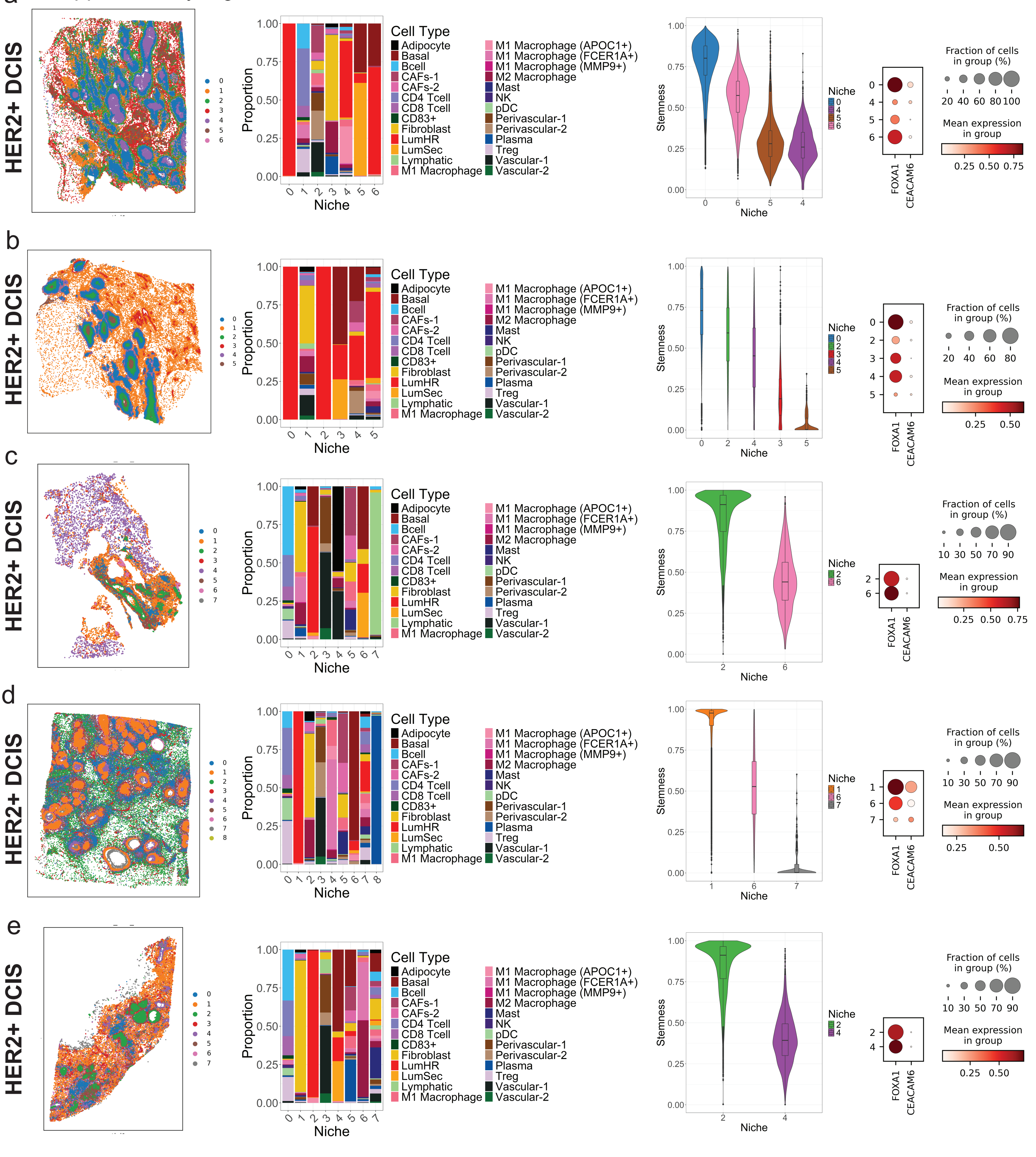

Supplementary Figure 8

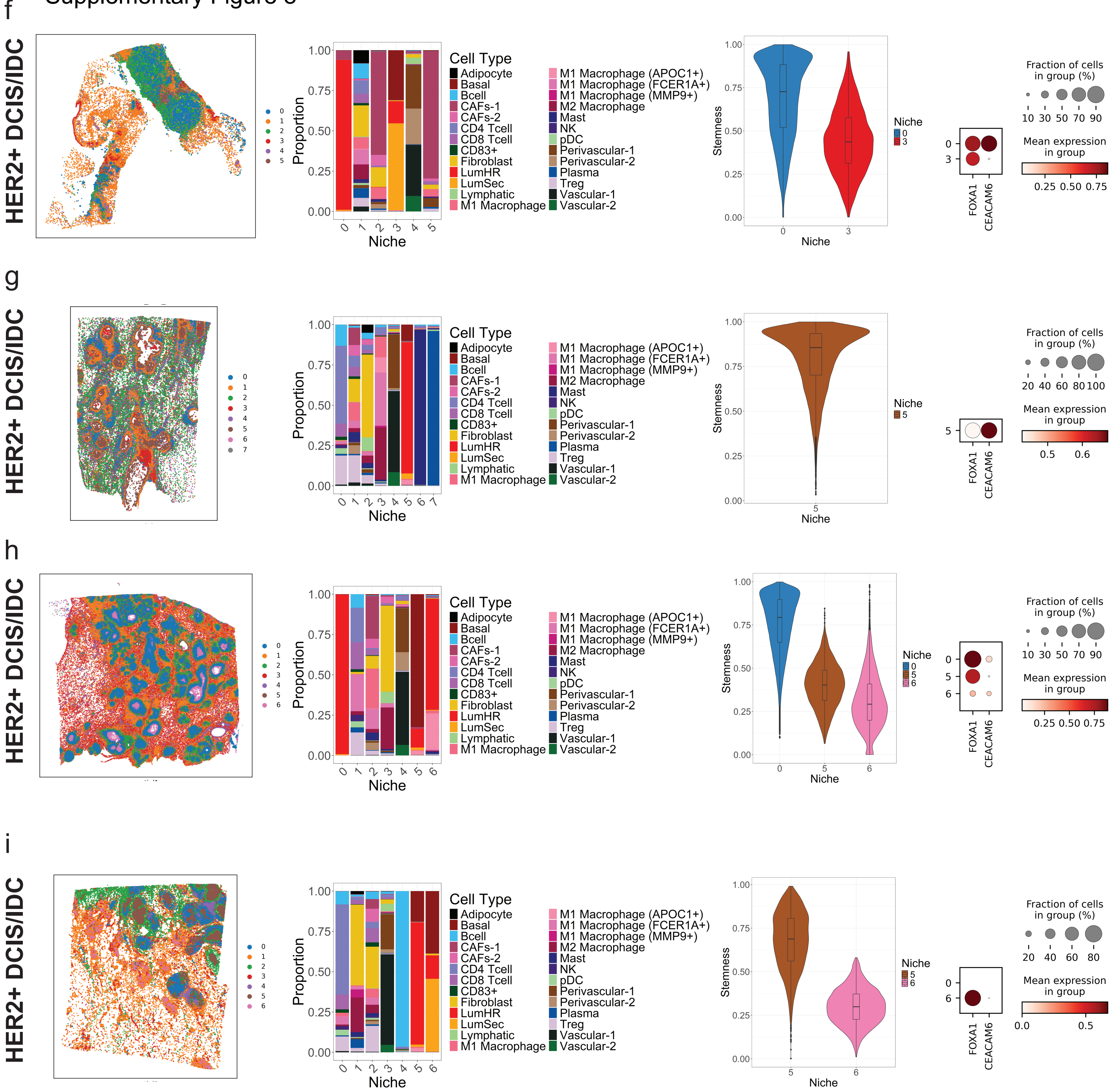

Supplementary Figure 8

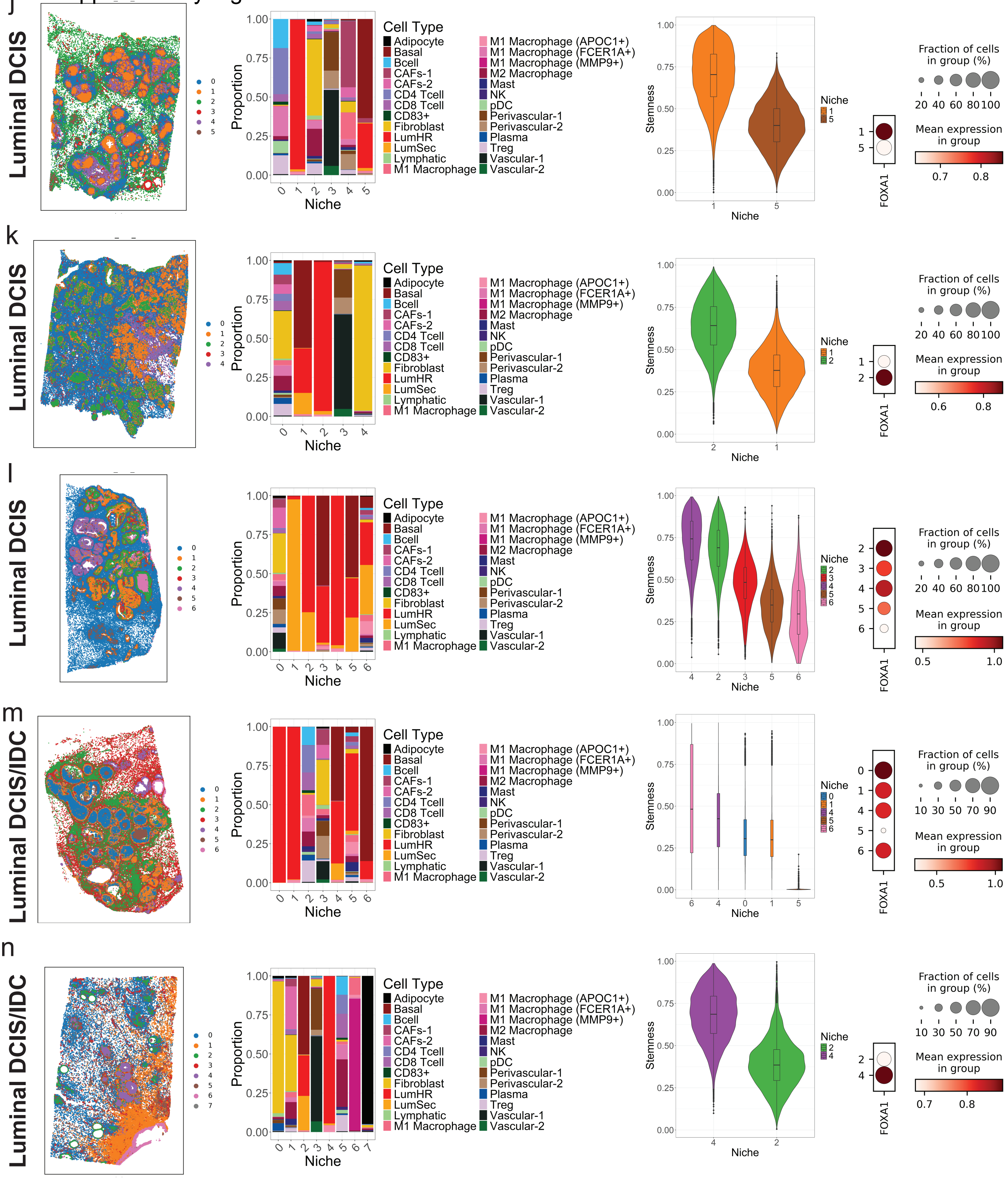

**Supplementary Figure 8 | Sample-Wise Niche Analysis Integrating Spatial Organization, Cell Composition, Stemness, and Marker Expression**

**a-n.** Niche analysis results for all samples, each row shows the results for one sample. From left to right, spatial image with the cells colored in by niche, bar plot showing cell proportion in each niche, stemness score for LumHR cells in niches, dot plot showing FOXA1 and CEACAM6 expression in LumHR cells (**a-i**), or just FOXA1 expression (**j-n**).

Supplementary Figure 9

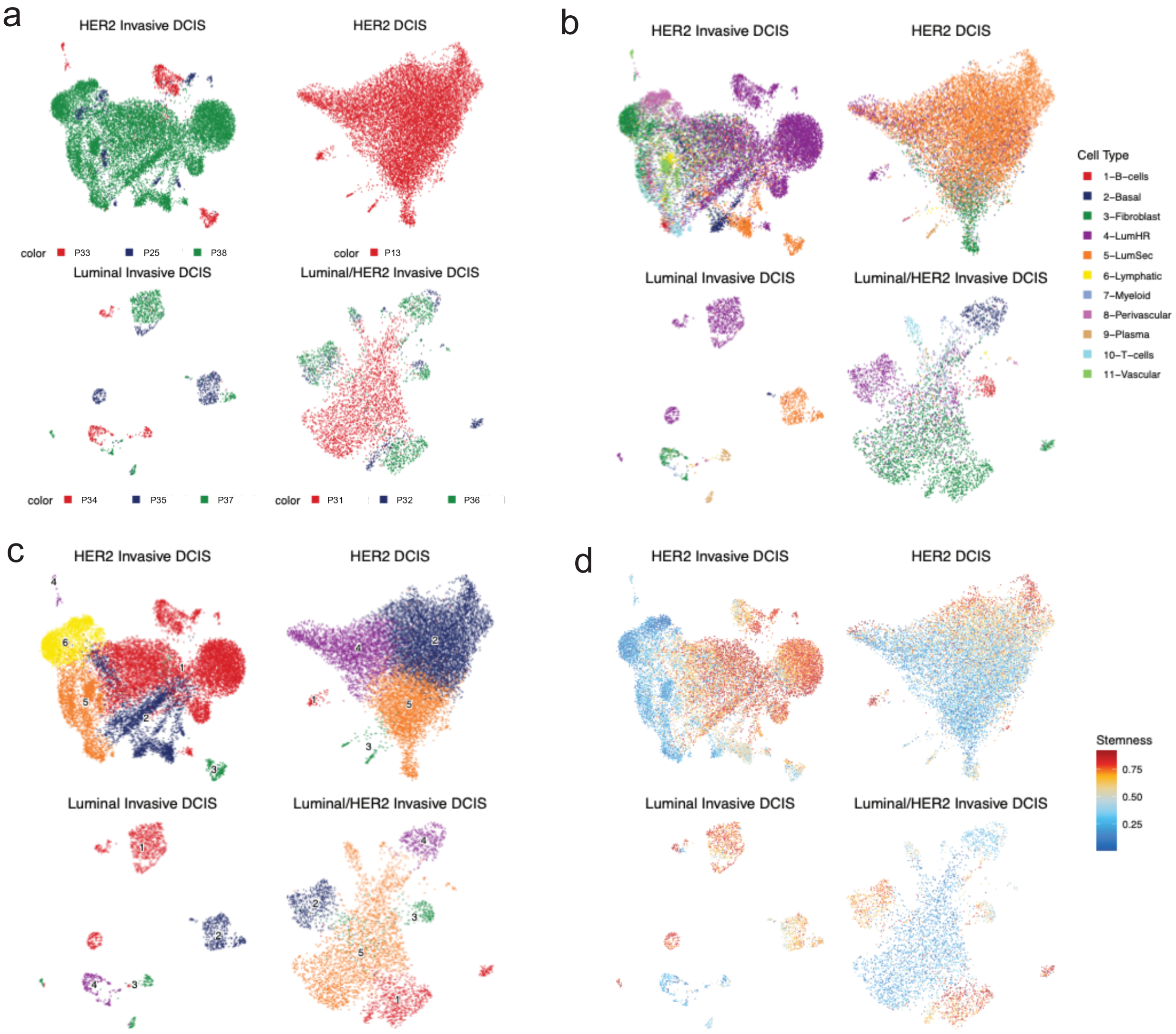

**Supplementary Figure 9 | UMAPs of scATAC-sequencing for each group of samples**

**a**, Colored in by sample ID

**b**, Colored in by cell type

**c,** Colored in by cluster

**d,** Colored in by stemness score
